# Exploration of the influence of environmental changes on the conformational and amyloidogenic landscapes of the zinc finger protein DPF3a by combining biophysical and molecular dynamics approaches

**DOI:** 10.1101/2025.02.24.640013

**Authors:** Julien Mignon, Tanguy Leyder, Antonio Monari, Denis Mottet, Catherine Michaux

## Abstract

In the past few years, the double PHD fingers 3 (DPF3) protein isoforms (DPF3b and DPF3a) have been identified as new amyloidogenic intrinsically disordered proteins (IDPs). Although such discovery is coherent and promising in the light of their involvement in proteinopathies, their amyloidogenic pathway remains largely unexplored. As environmental variations in pH and ionic strength are relevant to DPF3 pathophysiological landscape, we therefore enquired the effect of these physicochemical parameters on the protein structural and prone-to-aggregation properties, by focusing on the more disordered DPF3a isoform. In the present study, we exploited *in vitro* and *in silico* strategies by combining spectroscopy, microscopy, and all-atom molecular dynamics methods. Very good consistency and complementary information were found between the experiments and the simulations. Acidification unequivocally abrogated DPF3a fibrillation upon maintaining the protein in highly hydrated and expanded conformers due to extensive repulsion between positively charged regions. In contrast, alkaline pH delayed the aggregation process due to loss in intramolecular contacts and chain decompaction, the extent of which was partly reduced thanks to the compensation of negative charge by arginine side chains. Through screening attractive electrostatic interactions, high ionic strength conditions (300 and 500 mM NaCl) shifted the conformational ensemble towards more swollen, heterogeneous, and less H-bonded structures, which were responsible for slowing down the conversion into β-sheeted species and restricting the fibril elongation. For defining the self-assembly pathway of DPF3a, we unveiled that the protein amyloidogenicity intimately communicates with its conformational landscape, which is particularly sensitive to modification of its physicochemical environment. As such, understanding how to modulate DPF3a conformational ensemble will help designing novel protein-specific strategies for targeting neurodegeneration.

**Highlights:**

- DPF3a is a polyampholyte IDP, structurally sensitive to environmental changes.
- DPF3a amyloid pathway and propensity can be modulated by pH and ionic strength.
- Acidic condition inhibits fibrillation and maintains DPF3a in an extended state.
- Alkaline pH and ionic strength delay fibrillation by reducing structure collapse.
- DPF3a fibrils exhibit condition-dependent optical-morphological properties.

## 1. Introduction

Amongst the increasing number of health-concerning human diseases, cancer and neurodegenerative disorders still occupy the top leading causes of death and disability worldwide [1–4]. In such pathologies, intrinsically disordered proteins (IDPs), existing as a dynamical ensemble of interconverting conformations, have become an important class of regulators and promotors that are also prone to aggregate into cytotoxic amyloid assemblies [5–8]. Given their high structural plasticity and heterogeneity, weak intramolecular interactions, as well as their surface area extensively exposed to the solvent, the conformational ensemble of IDPs is highly sensitive and adaptative to their physicochemical environment. Therefore, function- and disease-driven variations in pH, temperature, ionic strength, membrane potential, salt type, redox balance, metallic homeostasis, or osmolyte concentration are defining factors in modulating the global and local structures of IDPs, displacing their (un)folded state equilibrium towards functional or pathological conformers [9–11].

For example, disruption of the balance of sodium, potassium, calcium, zinc, copper, and chloride ions promotes aberrant cell proliferation, tumoral mutagenesis, carcinogenesis, as well as cancer metastasis notably through chromatin decompaction and gene overexpression [12,13]. Growth of cancerous cells is advantageously impacted in hypoxia-induced acidic conditions. Indeed, tumour hypoxia and high glycolytic activity are known as acidosis inducers [14,15]. Moreover, balance between intra- and extracellular pH is highly relevant to carcinogenic pathways. For example, the extracellular pH is maintained low while the intracellular is alkalinised in colorectal cancer, which is highly favourable to tumour progression and invasiveness [16]. Regarding neurodegeneration processes, alteration in metallic homeostasis particularly affects the progression of Alzheimer’s (AD) and Parkinson’s diseases (PD) by (de)stabilising aggregation-prone protein folds upon binding [17–19]. Brain cells in patients suffering from AD locally display lower pH and increased ionic strength, conditions that respectively stabilise the assembly of amyloid β (Aβ) fibrils due to histidine protonation and enhance primary and secondary nucleation through screening the repulsion between Aβ negative charges [20,21]. Such physicochemical parameters are initiators and modulators of amyloid polymorphism that is not only encountered in *in vitro* experiments but also as a hallmark for defining different types of amyloidosis [22,23]. In the case of PD, studies have highlighted that slight modifications of pH and salt concentration modulate the nucleation and elongation rate of α-synuclein (α-syn), the diversity of its fibrillar repertoire, as well as its ability to efficiently bind anti-amyloid molecules [24–26].

The double plant homeodomain (PHD) fingers 3 (DPF3) is a human nuclear zinc finger protein belonging to the BRM/BRG1-associated factor (BAF) multiprotein chromatin remodelling complex, in which it acts as an epigenetic regulator by recognising modified motifs on histone tails [27–29]. Recently, a non-canonical BAF-independent role of DPF3 has been uncovered in mitosis and ciliogenesis. DPF3 depletion associated to the induction of kinetochore fibre instability and chromosome alignment defects that trigger apoptotic death, making it essential for proper cell division [30]. In previous research works, we unravelled the structural properties of DPF3 protein isoforms, DPF3b and DPF3a, which were revealed to be amyloidogenic IDPs [31–33]. Indeed, full-length and C-terminal variants of both isoforms were demonstrated to exhibit a high degree of intrinsic disorder and to spontaneously rearrange into β-sheeted high-order oligomers that elongate to form amyloid fibrils, features that are highly relevant to DPF3 physiopathology [34]. Upregulation of its gene expression profile is a hallmark in the Tetralogy of Fallot (TOF), a congenital cardiac hypertrophy, as well as in lymphocytic leukaemia, a blood and bone marrow cancer, in which it also promotes the proliferation of malignant cells [35–37]. DPF3 also connects to the motility of breast cancer cells, the stemness conservation of glioma-initiating cells, the prognosis of lung cancer, the prostate carcinogenesis, as well as in signalling pathways and apoptosis inhibition in renal cell carcinoma [38–43]. Furthermore, genomic and bioinformatic evidence suggest that DPF3 participates in the development and progression of AD and PD [44–46]. It can be pointed out that DPF3 is a metalloprotein, containing a conserved Krüppel-like C_2_H_2_ zinc finger (ZnF) binding two Zn^2+^ ions, as well as a PHD tandem for DPF3b coordinating four additional Zn^2+^ cations. We demonstrated that divalent metallic cations relevant to neurodegeneration were able to differently affect DPF3 aggregation propensity and mechanism depending on the isoform. While Mg^2+^ and Zn^2+^ cations reduced the protein fibrillation lag phase and promoted amyloid aggregation of DPF3a, Cu^2+^ and Ni^2+^ inhibited the fibril elongation and maturation steps [33].

In the light of its pathological repertoire, intrinsically disordered behaviour, and metal-impacted aggregation, the present study aims at expanding the knowledge on the amyloidogenic pathway of DPF3, as well as its conformational sensitivity to variations in the environmental pH and ionic strength, by focusing on the full-length DPF3a isoform. Drawing from our *in vitro* and *in silico* expertise, we combine experimental and computational approaches to decipher the factor-dependent structural properties of DPF3a, by confronting spectroscopic and microscopic data with all-atom classical molecular dynamics (MD) simulations.

## 2. Materials and methods

### 2.1. Protein overexpression and purification

Human recombinant full-length DPF3a protein was transformed in *E. coli* BL21 Rosetta (DE3) strains using a pET-like plasmid vector, containing the sequence for a GST fusion tag at the N-terminus position and a TEV-specific cleavage site. DPF3a-transformed bacteria were cultured in 20 g/L lysogeny Lennox broth (LB) medium supplemented with 0.36 mM ampicillin for approximately 16 h at 37 °C. Fractions of preculture were transferred in 20 g/L LB with 0.14 mM ampicillin and allowed to grow up to reaching an optical density at 600 nm comprised between 0.5 and 0.8. Isopropyl β-D-1-thiogalactopyranoside at a concentration of 0.5 mM was added to the media to induce protein expression for 4 h at 37 °C. Cultures were centrifuged, pelleted, and stored at −20 °C before purification. Bacterial pellets thawed at room temperature were suspended in lysis buffer (phosphate-buffered saline (PBS) pH 7.3, 0.5% Triton X-100, 200 mM KCl, 200 µM phenylmethylsulphonyl fluoride), sonicated in an ice-water bath, and centrifuged. For the purification procedure, collected supernatants containing GST-tagged DPF3a proteins were bound to a 5 mL pre-packed GSTrap FF column (Cytiva) with the binding buffer (PBS pH 7.3, 200 mM KCl), using an Äkta Purifier fast protein liquid chromatography system. Cleavage of the affinity GST tag was performed for 2 h at 30 °C with a GST-fused TEV protease (Sigma) and on column, previously equilibrated in Tris-buffered saline (TBS; 50 mM Tris-HCl pH 8.0, 150 mM NaCl). Resulting tag-free DPF3a proteins were eluted and retrieved in TBS. Dodecyl sulphate polyacrylamide gel electrophoresis and mass spectrometry ascertained the sample purity for the biophysical assays.

### 2.2. Protein concentration determination

After protein isolation and purity assessment, DPF3a eluates were concentrated in TBS using a dialysis membrane with a 6-8 kDa cut-off wrapped in PEG-20000 flakes for buffer absorption. Final protein concentration was determined at 214 nm, knowing the associated molar extinction coefficient of DPF3a (ε = 622615 M^−1^.cm^−1^) calculated with the method described by Kuipers B. and Gruppen H. [47]. Absorbance values at 214 nm were extracted from UV-visible absorption spectra recorded between 200 and 400 nm by 1.0 nm increment with a UV-63000PC spectrophotometer (VWR), using a 10 mm optical pathlength quartz QS cell (Hellma).

### 2.3. Protein sample preparation

In order to assess DPF3a susceptibility to pH and ionic strength, the protein was subjected to four distinct conditions in addition to the purified and concentrated material, i.e. pH 8 at 150 mM NaCl, taken as the reference: pH 2 and 12 at 150 mM NaCl, as well as pH 8 at 300 and 500 mM. Acidification, alkalinisation, and salting were achieved using 1 M HCl, 1 M NaOH, and 2 M NaCl stock solutions, respectively. In each sample, the working protein concentration amounted to ∼4.5 µM (0.18 mg/mL). For the aggregation monitoring, samples were subsequently incubated at ∼25 °C for seven days.

### 2.4. Far-UV circular dichroism spectroscopy

Far-UV circular dichroism (CD) spectra were recorded with a MOS-500 spectropolarimeter at ∼20 °C, using a 1 mm optical pathlength quartz Suprasil cell (Hellma) and the following acquisition parameters: 30 nm/min scanning rate, 2 nm bandwidth, 0.5 nm data pitch, and 1.0 s digital integration tome. For each tested condition, the buffer baseline was subtracted, and four scans were averaged and smoothed. CD data are presented as the mean residue ellipticity ([Θ]_MRE_), determined with the following equation: [Θ]_MRE_ = (Mθ)/(N-1)(10γl), where M is the protein molecular mass (Da; M_DPF3a_ = 40244 Da), θ is the measured ellipticity (mdeg), N is the protein sequence length, γ is the protein mass concentration (mg/mL), and l is the cell optical pathlength (cm). At different time points for all conditions, averaged spectra were fitted in the 195-250 nm range using the Beta Structure Selection (BeStSel) online tool to estimate the content in different secondary structures [48].

### 2.5. Steady-state intrinsic fluorescence spectroscopy

For all the fluorescence procedures, measurements were performed with an Agilent Cary Eclipse fluorescence spectrophotometer at ∼20 °C, using a 10 nm optical pathlength quartz QS cell (Hellma). Intrinsic phenylalanine (IPF), tyrosine (ITyrF), and tryptophan (ITF) fluorescence spectra were recorded from their respective excitation wavelength (260, 275, and 295 nm for IPF, ITyrF, and ITF, respectively) up to 600 nm with the following parameters: 1.0 nm data pitch, 0.1 s averaging time, 10 nm excitation-emission slit width (sw), 600 V photomultiplier tube (PMT) voltage, and 600 nm/min scanning rate. The same parameters were used for the acquisition of emission (λ_ex_ of 340 and 400 nm) and excitation (λ_em_ of 420 and 460 nm) autofluorescence (AF) spectra at the wavelengths mentioned in brackets. Complementarily, excitation-emission matrices (EEM) were recorded and constructed in each tested condition, using the parameters set as follows: 200-500 nm excitation range, 200-600 nm emission range by 5.0 nm λ_ex_ increment, 2.0 nm data pitch, 0.0125 s averaging time, 10 nm excitation-emission sw, 600 V PMT voltage, and 9600 nm/min scanning rate. For assessing the optical properties of DPF3a samples aged on long timescale, additional EEM scans were collected after six weeks of incubation at ∼25 °C. In all measurements, Raman scattering of water molecules was removed by applying appropriate excitation and emission wavelength filters. IPF, ITyrF, ITF, and AF data are presented as smoothed spectra, while EEM as colour-coded contour maps.

### 2.6. Transmission electron microscopy

After seven days of incubation at ∼25 °C, DPF3a aggregates were visualised by negative staining transmission electron microscopy (TEM) with a PHILIPS/FEI Tecnai10 electron microscope operating at a voltage of 80 kV. Beforehand glow-discharged formvar/carbon-coated copper grids were set on top of 10 µL droplets of protein material for 3 min, and the excess was soaked up afterwards with a blotting paper. Grid were stained with 0.5% (w/v) uranyl acetate as the contrasting agent for 1 min and air-dried for 5 min before being inserted into the microscope.

### 2.7. System preparation for molecular dynamics simulations

The starting tertiary structure of full-length DPF3a was generated from its amino acid sequence (Uniprot ID: Q92784-2) using the combined approach MMseqs2-AlphaFold2 available on the ColabFold platform and selecting the resulting first ranked and relaxed model [49,50]. As the presence of ions in metal-binding folded domains was not considered during the structure prediction, Zn^2+^ cation was manually placed in the C_2_H_2_ ZnF of DPF3a. Given the very high conservation of such Krüppel-like motif amongst the family of DPF proteins, the crystallographic structure of DPF2 C_2_H_2_ (PDB ID: 3IUF) [51] was aligned to that of DPF3a model, Zn^2+^ alignment was saved, and the protonation state of cysteine (CYM) and histidine (HID) residues involved in the metal coordination was adequately edited using the PyMOL software [52]. Extreme pH conditions tested *in vitro* (pH 2 and 12) were accounted for statically by estimating the pKa of ionisable groups from the three-dimensional model of DPF3a with PROPKA 3.2 [53,54], and by (de)protonating aspartate (ASH), glutamate (GLH), lysine (LYN), and cysteine (CYM) residues, accordingly. Protonation of the backbone amide and other sidechain atoms was carried out with the ProteinPrepare tool implemented in the PlayMolecule server [55]. Exploiting the *tleap* module of AmberTools, each differently protonated DPF3a model containing one coordinated Zn^2+^ cation was centred in a truncated octahedron water box. From each edge, a buffer of 10 Å of water molecules was added and the systems were neutralised with the required minimal amount of Na^+^ or Cl^−^ ions. For mimicking the experimental ionic strength conditions, 150, 300, and 500 mM NaCl concentrations were enforced by randomly adding an equivalent quantity of Na^+^ and Cl^−^ calculated with respect to the volume of the simulation box (Table S1).

### 2.8. Molecular dynamics simulation and trajectory analysis

All-atom classical MD simulations of full-length DPF3a in different pH and ionic strength environments were carried out in independent triplicates with the GROMACS 2023.1 suite [56,57]. For each system, the protein was modelled with the atomistic AMBER ff14SB force field [58]. Additionally, grid-based energy correction map (CMAP) parameters, compatible with the AMBER FF14SB force field, were added to identified DPF3a IDRs, i.e. residues 90-199, 222-260, and 293-357, to more accurately depict its intrinsic disorder [59]. Given the importance and performance of IDP-oriented solvation models, water molecules were explicitly considered and consequently described with the TIP4P-D model for a better sampling of the protein conformational space [60–62]. Along with RATTLE and SHAKE constraints, the use of hydrogen mass repartitioning (HMR), consisting in the redistribution of the mass of non-solvent hydrogen atoms to reduce high-frequency motion, enabled to increase the timestep from 2 to 4 fs for integrating the Newton’s equations of motion [63,64]. To this end, initial topologies were modified to HMR via the *ParmEd* package available in AmberTools [65]. The structure of the simulated systems was minimised for a maximum of 50000 steps using the steepest decent algorithm [66] before they have been thermalised, equilibrated, and propagated in the isothermal and isobaric (NPT) ensemble (Table S1). The Parrinello-Rahman barostat and a modified Berendsen thermostat (velocity rescaling method) were used to maintain a constant pressure of 1 bar and temperature of 300 K, respectively [67,68]. While long-range coulombic interactions were accounted for via the Particle Mesh Ewald (PME) summation, short-range non-bonded interactions were computed with a distance cut-off of 1.0 nm [69]. In combination with the leap-frog integrator for the solving the equations of motion, bonds were constrained to their proper lengths with the LINCS algorithm [70,71]. Production MD replicates were propagated in periodic boundary conditions (PBC) in all three dimensions for 1 µs each. Atom coordinates and energies were saved every 40 ps.

Trajectories were analysed with modules straightforwardly invoked from the GROMACS suite or with in-house Python scripts. Apart from clustering, contact maps, and captured snapshots, time-dependent variables were calculated and averaged from the resulting triplicates for each system. From the initially minimised and equilibrated structure, root-mean square deviation (RMSD) was computed solely from the backbone atoms. Protein backbone flexibility was assessed with the root-mean square fluctuation (RMSF) after alignment of the time-averaged structure as a reference using the following mathematical expression: 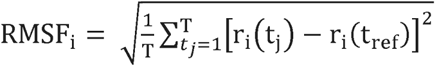 where T corresponds to the analysed time frame and r_i_(t) is the position of a given atom i at time t. With respect to the protein centre of mass, as well as N- and C-termini Cα, radius of gyration (R_g_) and end-to-end distance (d_ee_) were determined over the course of the simulation and plotted against each other into two-dimensional density graphs using the bivariate Kernel density estimator. The solvent accessible surface area (SASA) and number of contacts within 5 Å were calculated for the protein as a whole. The number of intramolecular protein and backbone H bonds was also computed, the latter being defined by a maximum distance cut-off of 3.5 Å and angle of 30° between the donor and the acceptor. Trajectories were separately clustered over the 1 µs-replicates using a RMSD cut-off of 5 Å between the nearest structural neighbours and a timestep of 1 ns, totalising an ensemble of 3000 frames in each system. For each condition, the central structure of the most populated cluster amongst the triplicates was extracted to map intramolecular contacts by computing the minimum distances for every residue pairs. The same rationale was applied to the final structure of the selected representative trajectories. The determination of secondary structure was achieved with the STRIDE algorithm in combination with the Timeline VMD plugin [72]. The trajectories were visualised and the structures of the presented snapshots were rendered using the VMD software [73]. Linear distribution of the net charge per residue was computed and plotted over a sliding window of five amino acids with the Classification of Intrinsically Disordered Ensemble Regions (CIDER) webserver [74].

## 3. Results and discussion

### 3.1. Impact of pH and ionic strength on DPF3a structural properties over time

#### 3.1.1. Secondary structure

First, the effect of pH and NaCl concentration on the secondary structures of DPF3a was evaluated over the course of time by far-UV circular dichroism (CD) spectroscopy (Fig. 1). Regarding the referential condition, that is, at pH 8 and 150 mM NaCl, DPF3a signature expectedly transitions from a hybrid intrinsically disordered protein (IDP), presenting random coil and folded substructures, towards a predominantly antiparallel and twisted β-sheet footprint (Fig. 1B). This is noteworthy evidenced by the maximum at 201 nm and the red-shifted negative band centred at 225 nm, as well as supported by the percentages in secondary structures estimated by BeStSel, showing a gradual increase in antiparallel β-sheet content (Table S2).

**Figure 1.**
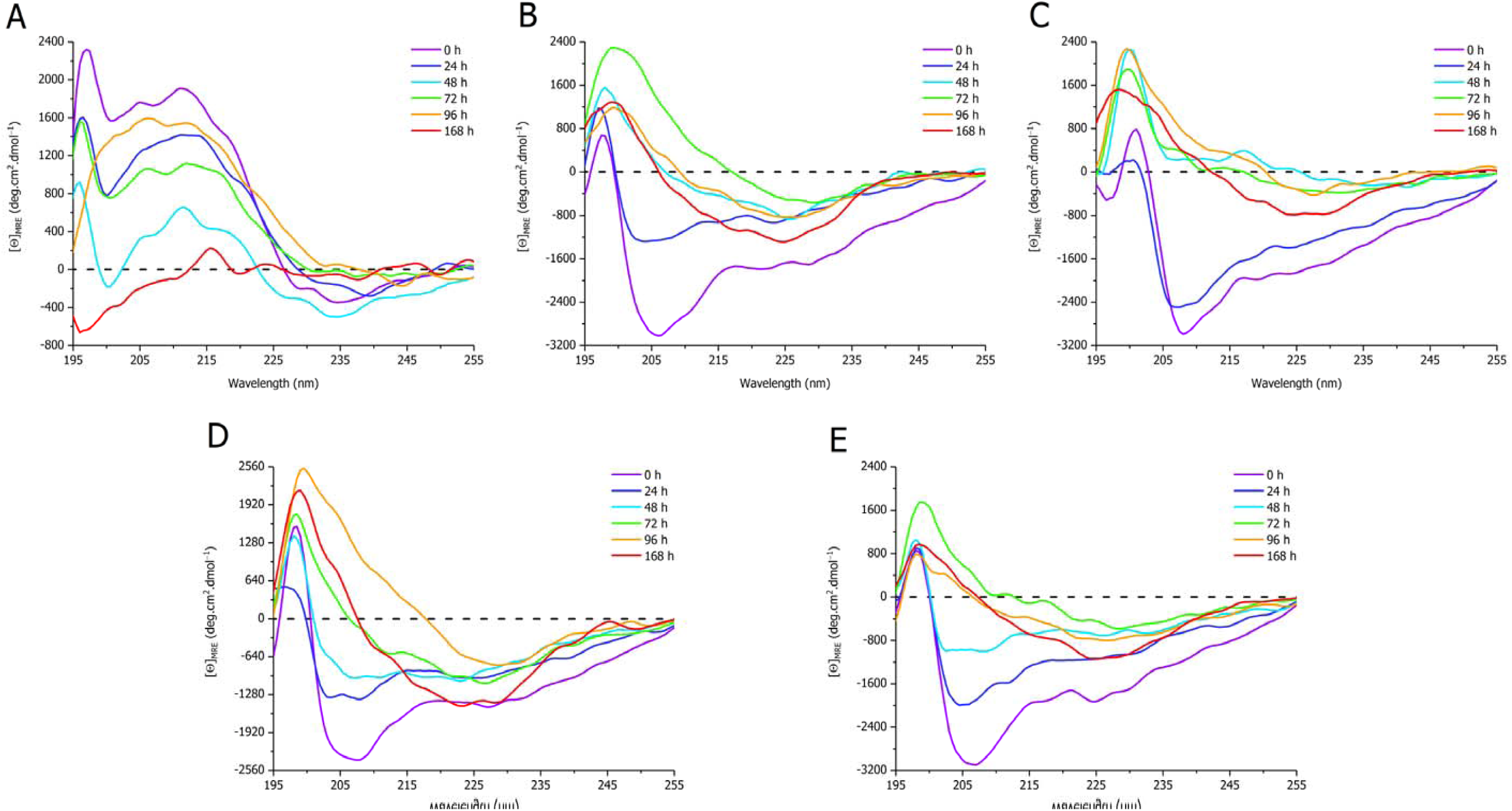
Far-UV CD spectra of full-length DPF3a incubated at ∼25 °C and (A) pH 2, (B) 8, and (C) 12 in 150 mM NaCl, as well as (D) 300 and (E) 500 mM NaCl at pH 8 for 0 h (purple), 24 h (dark blue), 48 h (light blue), 72 h (green), 96 h (orange), and 168 h (red).

Upon acidification to pH 2, the protein undergoes dramatical changes in secondary structure (Fig. 1A). Indeed, in the short wavelength range, a broad positive band spans from 205 to 225 nm, which corresponds to the conversion of random coil and folded domains into turns. This is consistent with the slight negative band close to the baseline in the 230-235 nm region, which is characteristic of β turns [4–6]. Interestingly, acidic conditions prevent the steady conversion into β-sheets over the seven days period of incubation mainly by maintaining DPF3a in a turn-disorder dynamic state, as observed by the fluctuations in band intensity between 195 and 200 nm and in BeStSel turn and coil estimations (Table S2). On the contrary, the main spectral features are conserved at pH 12, with the discernible formation of antiparallel and twisted β-sheets (Fig. 1C). Nevertheless, compared to pH 8, the overall spectrum is red shifted at 0 h, and the early structural conversion events within the first 24 h occur more slowly. This is characterised by a longer retention of helical elements and a high degree of disorder, as seen by the slight hypsochromic shift of the first negative band. Due to the broadening of the positive and negative bands towards higher wavelengths after several days, DPF3a could exhibit an overall higher proportion of turns at basic pH.

Regarding the saline conditions, the spectral profile remains similar to that of pH 8 in 150 mM NaCl at 0 h of incubation in terms of band position and intensity (Figs. 1D and E). Although after 168 h the spectra also exhibit a broad negative band centred at 225 nm, incubation from 0 to 48 h leads to a steadier decrease in the minima intensity, reflecting a slower transformation towards β-sheeted conformers, which appears to be further exacerbated at 500 mM NaCl (Fig. 1E, Table S2).

As such, alkaline pH and high ionic strengths delay the first steps of β conversion through the preservation of α-helix and coil structures, while not impairing the spontaneous rearrangement of DPF3a into β-sheets. The latter is comparatively hindered at acidic pH that favours a conformational ensemble enriched in turn and random coil regardless of the incubation time. While pH can disrupt the formation of H bonds needed for the formation and/or the stabilisation of secondary structure folds, high ionic strength can partly screen attractive interactions involved in the β-sheet transition, not only within but also between protein molecules. High salt concentration likely maintains the protein in a more unfolded state, which has been reported for other proteins [75] and is in very good agreement with the CD data and BeStSel estimates (Table S2), showing an increased coil proportion at 300 and 500 mM compared to 150 mM along the incubation period.

#### 3.1.2. Tertiary structure

Secondly, the intrinsic fluorescence of DPF3a aromatic residues was exploited to evaluate how the tested physicochemical factors affect the changes in tertiary structure occurring over the course of aggregation. Starting from the reference condition, intrinsic tryptophan fluorescence (ITF) spectra mainly show time-dependent changes in the emission band intensity rather than position for which maximum is centred towards 332 nm (Fig. 2B). The first 24 h intensity decrease indicates that the rearrangement pathway of the protein likely involves the relocalisation of tryptophanyl moieties to a polar environment, followed by a progressive hydrophobic bury of Trp residues in the next days. As this is not accompanied by a bathochromic shift of the band, rather than high solvent exposure, it could be attributed to packing near polar side chains and/or carbonyl group quenching effects, which are more probable to happen in an unfolded (disordered) or collapsed protein state [76,77]. Formation of such a more expanded intermediate is supported by intrinsic tyrosine fluorescence (ITyrF) spectra at pH 8 in 150 mM NaCl (Fig. 3B), revealing a shoulder in the short-wavelength range, corresponding to solvent exposed and/or Trp-free tyrosyl moieties [78]. Given the maximum emission wavelength on ITyrF spectra (∼329-332 nm), Trp-Tyr Förster resonance energy transfer (FRET) occurs in DPF3a, the efficiency of which appears enhanced over time due to the aggregation-driven rapprochement of Tyr and Trp residues, leading to the concomitant loss of the spectral shoulder observed at 24 h. Intrinsic phenylalanine fluorescence (IPF) spectra coherently display the same tendencies, as well as the fraction of Tyr residues being more exposed at 24 h, evidenced thanks to the persistence of Phe-Tyr FRET (Fig. S2B). Consistent with our previous reports, ITF, ITyrF, and IPF spectra exhibit a second emission band in the visible range appearing over time, a phenomenon which relates to DPF3a amyloid fibrillation and is referred to as autofluorescence (AF) [31,32,79].

**Figure 2.**
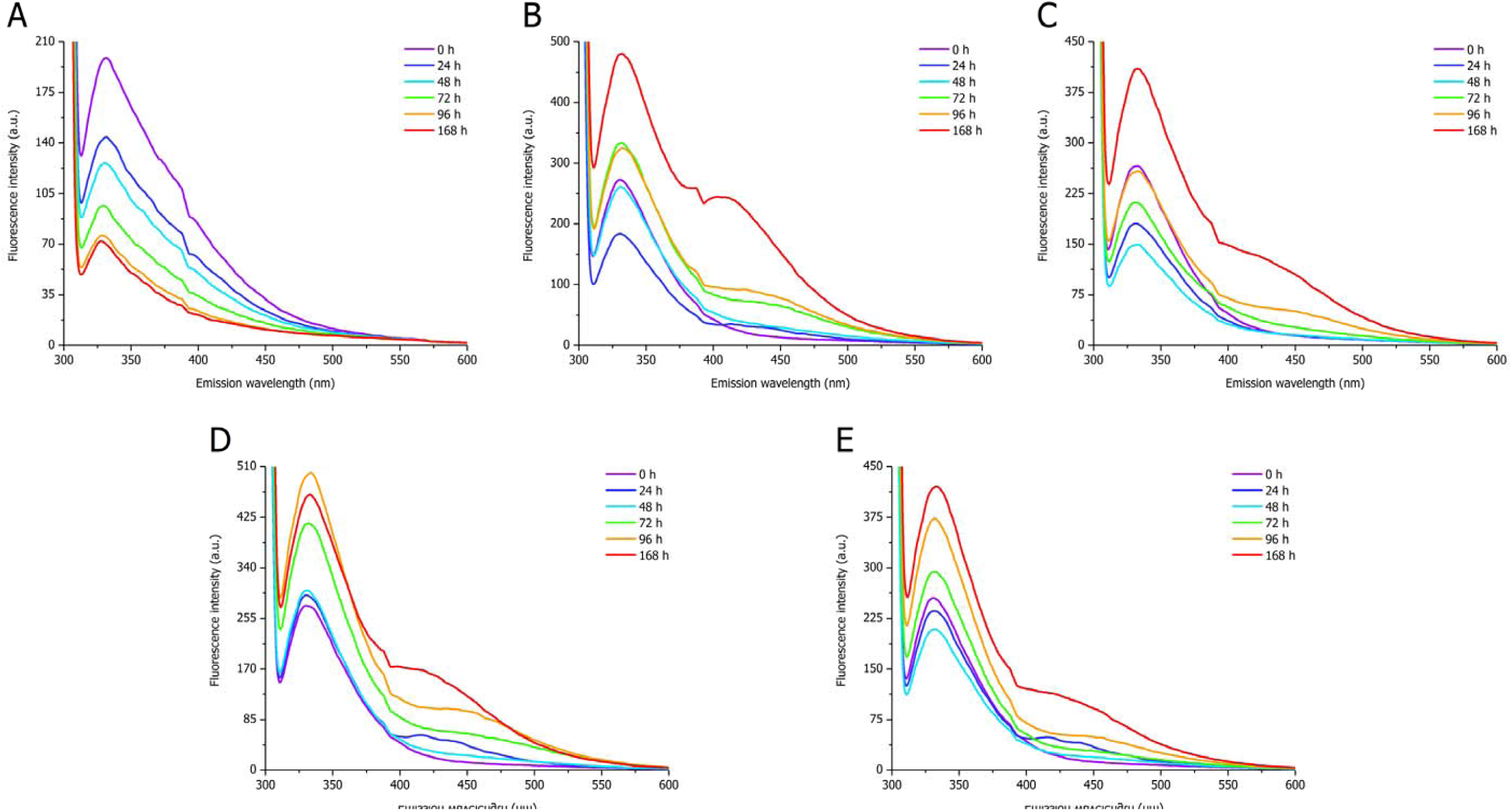
ITF spectra (λ_ex_ = 295 nm, sw = 10 nm) of full-length DPF3a incubated at ∼25 °C and (A) pH 2, (B) 8, and (C) 12 in 150 mM NaCl, as well as (D) 300 and (E) 500 mM NaCl at pH 8 for 0 h (purple), 24 h (dark blue), 48 h (light blue), 72 h (green), 96 h (orange), and 168 h (red).

**Figure 3.**
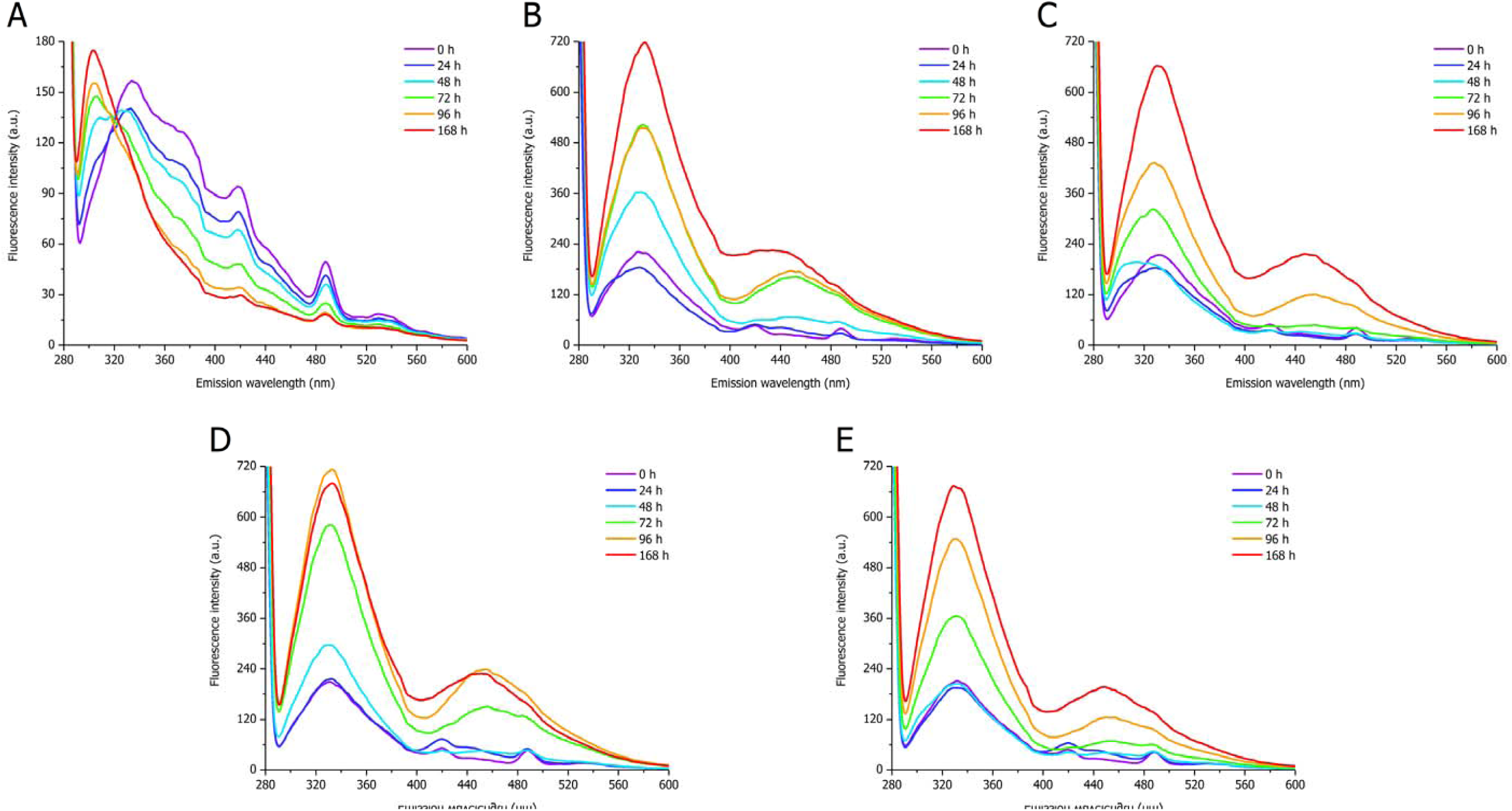
ITyrF spectra (λ_ex_ = 275 nm, sw = 10 nm) of full-length DPF3a incubated at ∼25 °C and (A) pH 2, (B) 8, and (C) 12 in 150 mM NaCl, as well as (D) 300 and (E) 500 mM NaCl at pH 8 for 0 h (purple), 24 h (dark blue), 48 h (light blue), 72 h (green), 96 h (orange), and 168 h (red).

Although the band position is mostly unaltered with a maximum at ∼330-332 nm in acidic pH, the ITF emission intensity is significantly decreased (Fig. 2A). Strikingly, such extinction carries out with time. While the quenching observed at 0 h can be explained by the reduction of Trp quantum yield at pH 2, it does account for its steady overtime decrease in band intensity [80]. In agreement with the CD analysis, it is therefore proposed that very acidic condition maintains DPF3a in a highly disordered and expanded conformational state that leads to high exposure of tryptophanyl moieties to a polar environment and hydrated carbonyl groups. The gradual hypsochromic shift of the Tyr band from 334 to 304 nm, which corresponds to the emission of non-buried Tyr and their distancing from Trp residues [81], resulting in the loss of Trp-Tyr FRET signal, corroborates such hypothesis (Fig. 3A). Similar band blue-shift is observed on IPF spectra (Fig. S2A), ascertaining that acidified DPF3a exists as a highly solvent-exposed conformer, and that Tyr-Phe FRET is still partly preserved. The overall weak ITyrF and IPF signals arise from the acid-mediated extinction of Tyr fluorescence [80]. Furthermore, the dual emission mode is no longer detected at pH 2, supporting the absence of amyloid AF emitters.

At pH 12, no change is observed with respect to the maximum emission wavelength, found between 331-333 nm on ITF spectra (Fig. 2C). However, further decrease of Trp fluorescence intensity is observed compared to pH 8, especially between 24 and 48 h. Such behaviour is particularly structurally relevant, as Trp fluorescence quantum yield is known to be enhanced above pH 10 [80,82]. The hydroxyl moiety of Tyr having a pKa value around 10, ionised tyrosyl has been reported to be a Trp quencher, which can also contribute to the relative intensity decrease [76]. Nevertheless, alkaline pH seemingly delays the conformational rearrangement or changes its pathway, hence maintaining in the 24-48 h time frame the tryptophanyl in a more quenching environment through interaction and/or spatial proximity with polar sidechains and carbonyl groups, before recovering Trp emission intensity through hydrophobic packing through amyloid-prone structural conversion. This is even better evidenced on ITyrF (Fig. 3C) and IPF (Fig. S2C) spectra, with the remarkable emergence of a spectral shoulder between 300 and 310 nm, which subsequently increases in intensity at 48 h before disappearing after 72 h (Fig. 3C), indicating that DPF3a first transitions to a more disordered state. Such a gain in disorder is corroborated by the associated far-UV CD spectra, showing a slight hypsochromic shift of the predominant negative band due to random coil enrichment (Fig. 1C). Further Tyr exposure also suggests distinct or delayed conformation rearrangements and aggregation pathway than that of the referential condition. From 72 h onwards, the emission band intensity is increased due to the burying of Tyr residues allowing Trp-Tyr FRET in the aggregated state. Slower alkaline-mediated fibrillation can be evidenced by the late generation of AF emitters on IPF, ITyrF, and ITF spectra consistently, which is coherent with the deferred β-sheet conversion in far-UV CD (Fig. 1C). Although it is most likely that Trp-Tyr FRET still occurs at pH 12, it cannot be excluded that part of the observed signature around 335 nm arises from tyrosinate emission [83,84].

In both high ionic strength conditions, the ITF band remains once more in the similar emission wavelength range (∼331-333 nm), irrespective of the salt concentration or incubation time (Figs. 2D-E). However, they do not display the same spectral variations over the course of time. In 300 mM NaCl, the intensity of the Trp band progressively increases with time, without passing through a Trp-quenching intermediate (Fig. 2D). Nevertheless, a significant increase in emission intensity is only observed after 72 h incubation due to the slower aggregation-induced formation of more efficient Trp-Tyr FRET conformers, as well as the burying of aromatic residues in a more hydrophobic core (Figs. 2D and 3D). Interestingly, 500 mM NaCl exhibits a similar overtime tendency to that of pH 12. Indeed, not only the Trp emission intensity is gradually decreased from 0 to 48 h before increasing (Fig. 2E), but also ITyrF spectrum presents after 48 h a shoulder towards 305 nm associated to exposed Tyr residues (Fig. 3E). IPF spectrum at 48 h also displays such hypsochromic band broadening due to Tyr-Phe FRET (Fig. S2E). Whilst the structural transformation locally differs between pH 12 and 500 mM NaCl due to the extent of each effect, the constated similarity reinforces that both conditions still partly hinder DPF3a aggregation through the permanence of expanded and hydrated conformers on a longer timescale. On the other hand, in 300 mM NaCl the lagged β-sheet enrichment does not seem to arise from the expansion of DPF3a structure, which is supported by the absence of short-wavelength shoulder on ITyrF (Fig. 3D) and IPF (Fig. S2D) spectra, suggesting a different rearrangement pathway compared to 150 and 500 mM. AF-related emission bands are also detected upon DPF3a aggregation regardless of the ionic strength and excited aromatic amino acid. As such, concomitant aromatic residues FRET enhancement and appearance of AF second emission band is a coherent indicator of DPF3a aggregation.

High ionic strength is known to screen long- and short-range interactions, either repulsive or attractive [11]. In the context of DPF3a, the protein displays a relatively low mean net charge at pH 8, and the charged residues are rather well scattered along the sequence or sequestered in small clusters of opposite charges (Fig. S1A). This is reflected by its moderate charge patterning κ value of 0.27, from which DPF3a is expected to occupy locally and/or globally collapsed conformations [85,86]. Therefore, high ionic strength is likely to screen attractive interactions between such distributed charged patches in DPF3a, leading the protein to occupy a more expanded, and thus, less compact conformational subspace with fewer intra- and intermolecular contacts, hence slowing down its amyloid rearrangement. Extreme pH conditions instead lead to sidechain (de)protonation that dramatically transform the overall charged state and charge distribution within DPF3a (Figs. S1B-C). While electrostatic repulsion between positively charged residues dominates at pH 2, resulting in the protein adopting a polyelectrolyte-like and solvent-exposed conformation, persistence of arginine (Arg) positive charge partly balances the more negative charge distribution at pH 12, thus not precluding aggregation.

### 3.2. Modulation of amyloid-driven autofluorescence

#### 3.2.1. Overtime evolution of autofluorescence

As evidenced by the previous spectra, DPF3a typically exhibits a dual emission band appearing over the course of time, which is intrinsically associated to the amyloid propensity of the protein [32,33,79]. In a more recent investigation work, we were able to better appreciate the origins of such photoluminescence in DPF3 proteins and the involvement of pH-mediated proton transfer and charge recombination, as well as to discriminate different autofluorescence (AF) modes according to their emission range, speculated to prove relevant to the inner and macroscopic structure of fibrils [79]. In the same research work, it was also shown that mature fibrils of DPF3a formed in 150 mM NaCl at pH 8 were characterised by an emission maximum centred at 415 nm, designated as violet autofluorescence (vAF), upon illumination at 340 nm.

Drawing from our knowledge of DPF3a AF properties, emission spectra were first monitored over time after excitation at 340 nm (Fig. 4). In the reference condition, the AF fingerprint expectedly appears and increases in intensity along the protein aggregation, reaching an emission maximum at 415 nm after 168 h (Fig. 4B). Remarkably, a gradual shift from the deep-blue autofluorescence (dbAF, λ_em_ of 445 nm) to the vAF (λ_em_ of 415 nm) region is observed over the course of time. Moreover, the three conditions at pH 8 exhibit a common dual emission band at 24 h, presenting two maxima at ∼418 and ∼440 nm, respectively (Figs. 4B, D-E). This is interesting because they correspond to two different AF modes. While dbAF becomes dominant after 48 h, it progressively shifts towards the vAF mode. As it is no longer detected upon pH variation, it informs that generation of such vAF-dbAF intermediates is only sensitive to the modification of the protein charged and protonated state. Taken together, these spectral fluctuations reveal that the photophysics of DPF3a is quite dynamic during its aggregation. At the beginning of the structural rearrangement, weak vAF and dbAF fluorophores coexist, the latter being favoured in the first self-assembly steps, before being supplanted by strongly vAF-emitting fibrils after elongation. Whilst the optical trend appears similar, the ionic strength affects the position of the AF bands. After 168 h, the major vAF band is found at 425 and 428 nm in 300 and 500 mM, respectively (Figs. 4D-E). Furthermore, during the first aggregation steps, the dbAF maximum is also red shifted compared to 150 mM NaCl, at around 450 and 455 nm for 300 and 500 mM, respectively. Such differences are likely associated to distinct oligomeric and fibrillar structures defined by ionic strength. Nonetheless, discrepancy in AF emission intensity, particularly after 168 h of incubation, reinforces that the formation of vAF-emitting species, associated to the elongation and maturation of fibrils, is hindered as the ionic strength increases (Figs. 4D-E). Regarding the pH conditions, acidification results in no AF signal whatsoever (Fig. 4A), as only weak and artefactual UVA signals that cannot be attributed to AF are detected. In contrast, pH 12 AF spectra show a gradual increase in emission intensity from 72 h onwards, accompanied by the hypsochromic shift of the band position towards the vAF region, supporting that DPF3a fibrillation is delayed upon alkalinisation (Fig. 4C). While the excitation spectra of the vAF mode were also analysed in each tested condition (Fig. S3), they did not show any significant variations at the level of the main AF contributor, at the exception of pH 2 for which it was accordingly completely extinguished (Fig. S3A).

**Figure 4.**
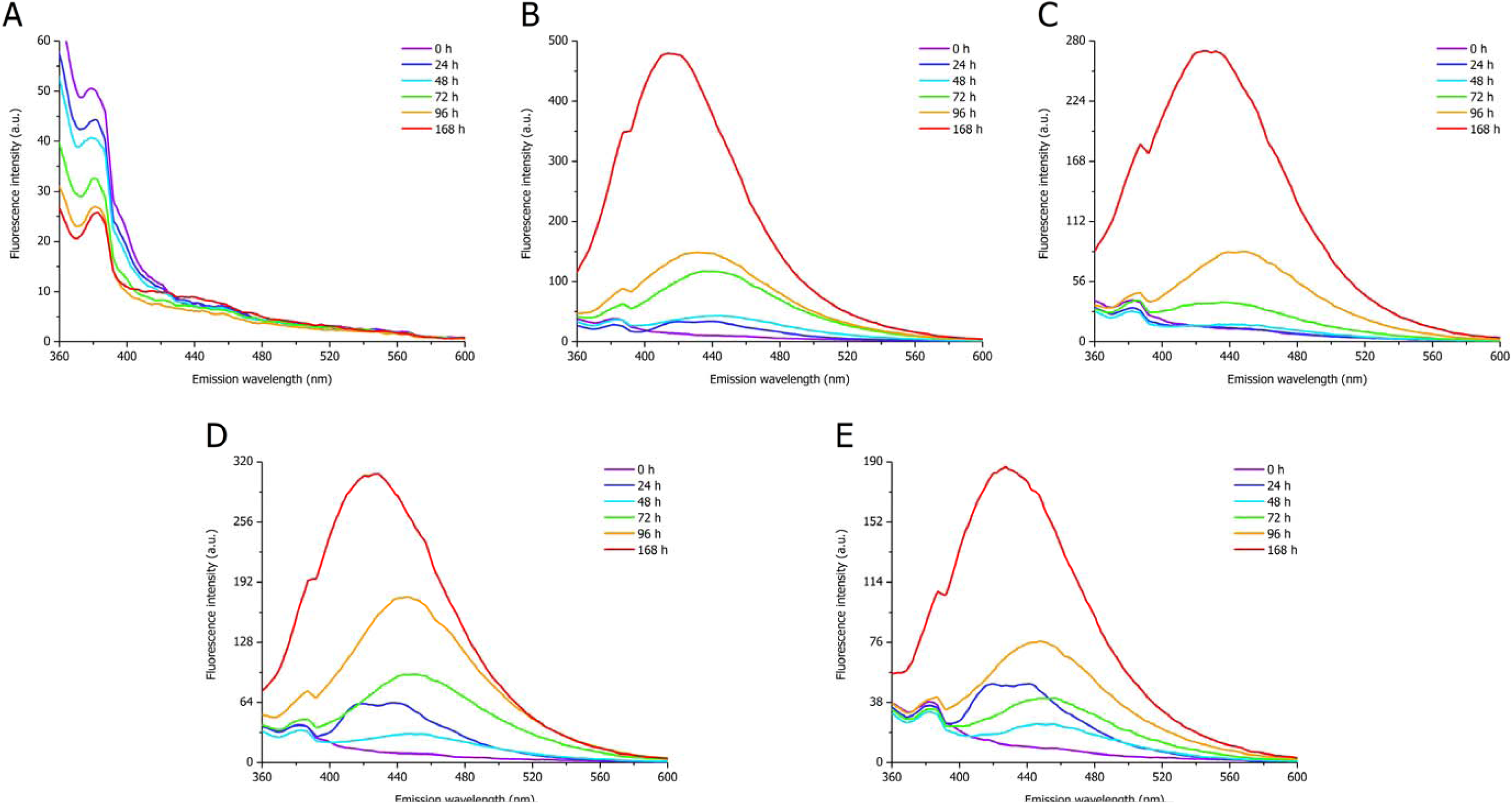
AF spectra (λ_ex_ = 340 nm, sw = 10 nm) of full-length DPF3a incubated at ∼25 °C and (A) pH 2, (B) 8, and (C) 12 in 150 mM NaCl, as well as (D) 300 and (E) 500 mM NaCl at pH 8 for 0 h (purple), 24 h (dark blue), 48 h (light blue), 72 h (green), 96 h (orange), and 168 h (red).

Scanning at longer excitation wavelength, i.e. 400 nm, reveals at first glance that dbAF emission properties are far less sensible to changes in the protein environment compared to the vAF populations, which is reflected by the absence of any band shift (Fig. S4). Although intensity discrepancies are observed between the alkaline and saline conditions, an overall increase of the band located at around 460 nm is observed over time (Figs. S4B-E). However, examination of their associated excitation spectra, by fixing the emission wavelength at 460 nm, unveiled interesting trends, allowing to discriminate the effect between conditions (Fig. S5). Indeed, while the main long-wavelength dbAF contribution is situated at 400 nm after 48 h, a second absorption band appears from 72 h at 350 nm and increases in intensity to finally be the dominant band (Fig. S5B). Comparatively, emergence of the 350 nm contributor is delayed upon increasing the pH or the ionic strength, only becoming distinguishable after 168 h of incubation (Figs. S5C-E). Consistent with the data presented so far, formation of vAF emitters and appearance of the band at 350 nm are indicative of fibril maturation. Neither of the identified excitation contributors was detected at pH 2 (Fig. S5A). While not further detailed hereafter, it can be pointed out that, in every excitation spectra (Figs. S3 and S5), slight changes are also observed at the level of the peptide bond absorption band (λ_ex_ range of 230-240 nm) and excitation contribution of aromatic residues (λ_ex_ range of 260-295 nm), which participate in defining the AF modes and are of significance to the factor-altered optical activity of fibrils.

#### 3.2.2. Autofluorescence landscape

Other AF modes, such as dbAF and blue-green (bgAF) autofluorescence, have also been identified in DPF3a fibrils and can be photoselected at long-range excitation wavelength. In this regard, the photoluminescence landscape of DPF3a was further explored by recording excitation-emission matrix (EEM) at different time points and represented as coloured contour plots (Figs. 5, S6, and S7). Overall, the matrices unveiled evocative dynamics between AF populations, as well as parameter-dependent changes in the optical response of DPF3a. At pH 8 and 150 mM NaCl, fibril growth is characterised by a progressive shift from a continuous spectral subspace overlapping dbAF and vAF regions with a weak maximum at λ_ex_/λ_em_ of 347/436 nm (Fig. S6B) towards a more intense and dominant vAF population centre at λ_ex_/λ_em_ of 340/420 nm (Fig. 5B). At pH 2, matrices were unambiguously devoid of any AF emitters, even after one week of incubation (Figs. 5A and S6A), which is consistent with the previous CD and intrinsic fluorescence spectra, advocating for the acidic abrogation of the fibrillation process. Interestingly, only the dbAF mode is detected at λ_ex_/λ_em_ of 393/457 nm after 96 h upon alkalinisation (Fig. S6C), and vAF-emitting species appear later in the aggregation process through the extension of the AF space up to a distinct maximum at λ_ex_/λ_em_ of 345/430 nm (Fig. 5C). Similar evolutions are observed at high ionic strength for which dbAF emitters arise first and foremost at maximum around λ_ex_/λ_em_ of 400/457 nm (Figs. S6D-E) before transitioning to vAF fluorophores near λ_ex_/λ_em_ of 345/425 nm (Figs. 5D-E). One noticeable difference consists in the absence of intensity discrepancy in the AF subspace after 168 h in 300 mM NaCl, as both vAF and dbAF populations coexist in the same proportions (Fig. 5D), meaning that such condition retains and stabilises more dbAF-fluorescing assemblies. In 500 mM NaCl, the overall spectral intensity is also decreased compared to 150 and 300 mM (Fig. 5E), reinforcing that fibrillation is further hindered. The consistently observed delay in the vAF formation once more advocates for alkaline and high ionic strength partly hindering DPF3a amyloid fibrillation due to less favourable charge repulsion and screening of attractive interactions, respectively. Furthermore, dbAF emitters seem to correspond to transient species, which could also be fibrillar, as it was shown that fibrillation is a highly polymorphic, dynamic, and evolving process engaging kinetically and thermodynamically favoured aggregated states, communicating with and cannibalising each other over the course of time [87]. It can be also pinpointed that only very weak bgAF exists, so much so that it is barely detected on the EEM. As such, pH and ionic strength conditions do not seem to promote the formation of bgAF emitters, fibrils only existing in a dbAF-vAF continuum.

**Figure 5.**
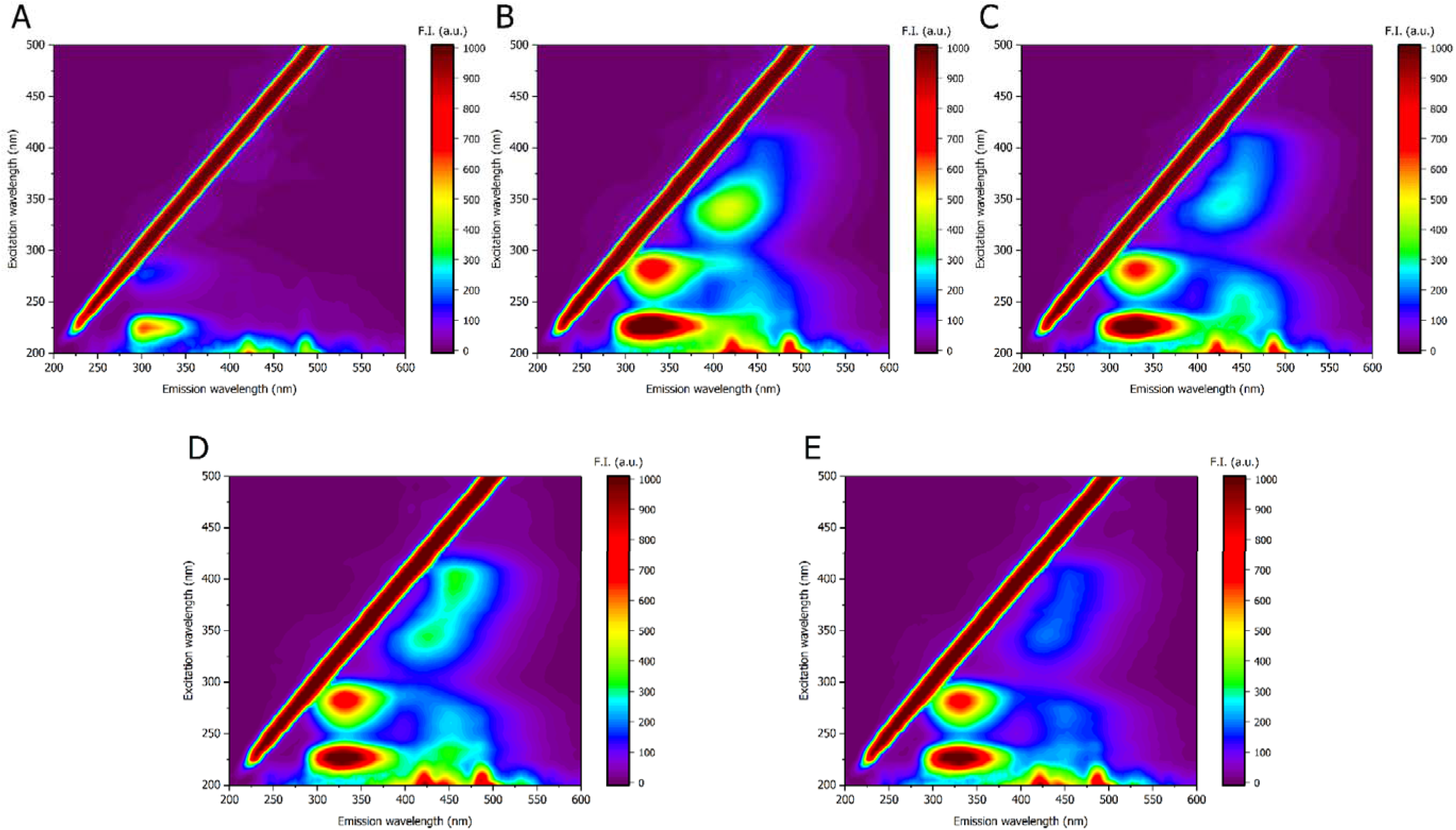
EEM (sw = 10 nm) of full-length DPF3a incubated for 168 h at ∼25 °C and (A) pH 2, (B) 8, and (C) 12 in 150 mM NaCl, as well as (D) 300 and (E) 500 mM NaCl at pH 8.

**Figure 6.**
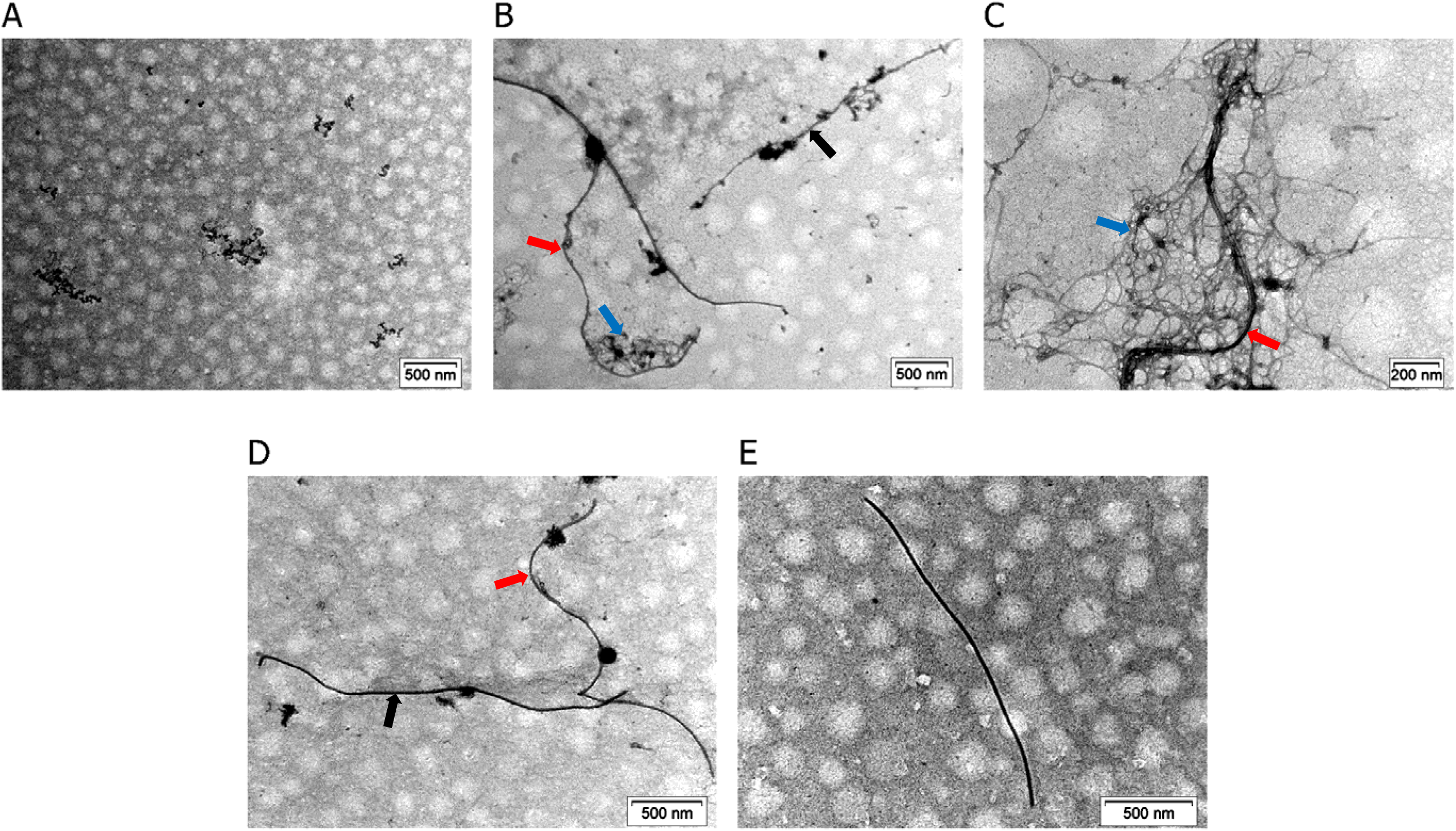
Negatively stained TEM micrographs of full-length DPF3a incubated for 168 h at ∼25 °C and (A) pH 2, (B) 8, and (C) 12 in 150 mM NaCl, as well as (D) 300 and (E) 500 mM NaCl at pH 8. (A) Amorphous phases. (B) Coexistence of straight (black arrow) and curved (red arrow) fibrils entangled with thin cobweb-like fibrillar networks (blue arrow). (C) Curved and undulating fibrils (red arrow) entangled within cobweb-like fibrils (blue arrow). (D) Coexistence of straight (black arrow) and curved (red arrow) fibrils. (E) Short and isolated straight fibril. The scale bar is provided at the bottom right of each micrograph.

Long-term incubation uncovered very interesting changes in AF matrices, all presenting the same excitation-emission landscape with a neat maximum in the vAF mode of comparative importance (Fig. S7). More precisely, while the initial dbAF population at λ_ex_/λ_em_ of 400/457 nm is no longer present, vAF emitters slightly shift towards λ_ex_/λ_em_ of 340/410 nm and consequently increase in intensity after 6 weeks of incubation (Fig. S7B), which is attributed to the accumulation of fibrillated material over time. The same behaviour is remarkably observed at pH 12, as well as in 300 and 500 mM NaCl (Figs. S7C-E). However, in 300 mM, a fraction of dbAF-emitting aggregates persists over 6 weeks (Fig. S7D), a population that is completely cannibalised by the predominant vAF-emitting fibrils in the other conditions. Compared to 150 mM NaCl, higher ionic strength leads to an overall reduction of the vAF intensity, which signifies that a lower proportion of mature fibrils is formed and/or arises from fibril shortening due to elongation impairment [33,88,89].

Taken together, these results suggest that alkaline and high ionic strength conditions alter the kinetically favourable amyloidogenic pathway of DPF3a, whereas over time the equilibrium between different fibrillated subpopulations is displaced towards the formation of a common and presumably more thermodynamically favourable state, leading to the same dominant vAF emitter regardless of the condition the protein was initially aged in. Also, it seems to indicate that an overall similar pathway could be taken by DPF3a in different conditions as the same succession of AF fluorophores is formed throughout the fibrillation, although with delayed times in high pH and salt concentration, and all finishing with the same optical properties, which could indicate comparable morphologies. Nevertheless, subtle changes are still detected at the levels of the peptide bond and aromatic absorption bands, which could reflect subtle differences in the inner fibril structure between conditions. In good agreement with CD, ITF, ITyrF, and IPF spectra, conspicuous absence of any autofluorescence emitters at pH 2, even after 6 weeks of incubation (Fig. S7A), further ascertains that not only very acidic environment completely inhibits the protein amyloidogenic pathway, but also AF intimately arises from DPF3a amyloid fibrillation.

### 3.3. Parameter-induced morphological diversity of DPF3a fibrils

The morphological features of DPF3a aggregates and their correlation with AF signatures were assessed by negative staining transmission electron microscopy (TEM, Fig. 6). Coherently, none fibrillar structure was found at pH 2, with only small amorphous phases and clusters sporadically scattered on the grid (Figs. 6A and S8). In the reference condition, typical 18-30 nm wide straight fibrils are mainly found along with curved and undulating fibrils of the same width (Figs. 6B and S9A-B). Most of the observed fibrils exhibit very extended length due to non-limited elongation step and favourable conversion of free monomers at fibril extremities (Figs. S9B, D-E). Intriguingly, some fibrils are localised close to or entangled within dense fibrous patches resembling scrambled cobwebs. Upon increased magnification, they revealed to be networks of thinner fibrils, between 5 and 10 nm in width, some of them associating into (Fig. S9C) or with larger ones (Fig. S9F). Such structure was never reported for DPF3a and could either correspond to a new polymorph or to the transient dbAF-emitting fibrillar intermediates, 18-30 nm wide straight or curved fibrils rather exhibiting vAF emission. Cobweb-like fibrillar networks are also found in great number at pH 12 (Figs. 6C and S10A-B), associating with (Fig. S10C) or localising next to mature fibrils (Fig. S10F). While the typical straight and curved fibrils of comparable morphology to that of pH 8 are spotted (Figs. 6C and S10D-E), they are statistically of shorter length and in lower quantity. The same trend is observed at high ionic strength, the fibrillar populations becoming shorter and in lesser number the higher the salt concentration (Figs. 6D-E, S11A-B, S11E, S12A-B, and S12F). Moreover, similar undulating fibrils are still present, although in a reduced occurrence (Figs. S11C and S12C). Albeit not gathered in cobweb-like networks, thin fibrils associating into wider ones are also visualised in 300 (Fig. S11F) and 500 mM NaCl (Figs. S12D-E).

TEM micrographs are very consistent with the previously discussed spectroscopic data, demonstrating that DPF3a fibrillar and/or amorphous aggregates exhibit distinctive optical-structural features, as hypothesised in our previous investigation [79]. Indeed, high pH and ionic strength hindrance effect on DPF3a aggregation and fibril elongation, due to charge repulsion and screening of attractive interactions, not only manifests through late amyloid-driven spectral changes but is also reflected in the low proportion of mature fibrils of significant shorter length. In the light of delayed fibrillation, thin fibrils often found in cobweb-like networks are likely associated to early aggregation dbAF emission, and are, as such, transient species clustering or morphing into wider straight or curved fibrils fluorescing in the vAF region. Furthermore, absence of fibrillar or amyloid-like assemblies, conjugated with the inhibition of β-sheets and AF emitters formation, substantiates that very acidic environment completely suppresses the amyloidogenic pathway of DPF3a.

Compared to alkaline conditions, such behaviour informs that disruption of DPF3a fibrillation process is more sensitive to protonation and the generation of positive charge, as the protein net charge changes from −5 (pH 8) to +59 (pH 2) (Table S1). In this regard, highly positively charged DPF3a monomers are not able to interact and nucleate into amyloid precursors, resulting in photoinactive species and amorphous phases. On the other hand, alkaline-mediated deprotonation, albeit unfavourable due negative charge repulsion, could be compensated by the formation of non-native intra- and/or intermolecular disulphide bridges, enabling another pro-amyloid pathway, resulting for example in the increased proportion of cobweb-like fibrous networks. Indeed, thiolate moieties are tenfold more reactive than their thiol form, and the thiol-disulphide exchange reaction is facilitated in alkaline medium [90,91]. In addition, DPF3a is particularly enriched in cysteine (Cys) residues accounting to a total of twelve, representing 3.4% of the total sequence content, which is greater than what is usually found across the eukaryote proteome (2.2%) [92]. Two Cys residues directly bind one Zn^2+^ cation in the C_2_H_2_ domain, whereas the other ten sulphydryl moieties can be available to undergo oxidation. Various amyloidogenic proteins have been reported to oligomerise and fibrillate thanks to the reactivity of their Cys sidechains. For example, interchain disulphide bonds are well-known to be conserved in the fibrillated state of insulin molecules [93]. While intramolecular disulphide bonds are stabilising protein into monomeric and non-amyloid conformers, intermolecular bridges enhance interactions and can be responsible for amyloidosis [94]. One can speculate that Cys residues found in DPF3a intrinsically disordered regions (IDRs), due to their availability and exposure, are more sensitive to oxidation and could drive disulphide bridging between DPF3a monomers, favouring self-assembly.

### 3.4. Molecular dynamics simulations

#### 3.4.1. Context-defined protein chain flexibility, compaction, and hydration

To deeper unravel the obtained *in vitro* results and to gain further insight into the effect of each physicochemical factor on defining the conformational properties of DPF3a at the molecular level, all-atom classical molecular dynamics (MD) simulations were performed on the microsecond timescale under pH and ionic strength parameters mimicking the experimental conditions. First, the overall conformational stability was assessed by analysis of the root-mean square deviation (RMSD) evolution with time (Figs. 7A-B). Regardless of the condition, a very pronounced and nanometre-scale RMSD variation is observed, which is typical of IDPs and indicative of a highly dynamic system. At pH 8 in 150 mM NaCl, DPF3a appears to converge towards a stabilised conformation beyond 400 ns (Fig. 7A). Upon alkalinisation, the conformational ensemble appears more complex, with significant fluctuations in RMSD values and high deviation between replicates all along the simulation frame, a behaviour that is further exacerbated at low pH. Compared to pH variations, increased ionic strength acts as a structural stabilising factor, each curve converging to a plateau with less variability between the replicates (Fig. 7B). However, the curve corresponding to 500 mM salt concentration presents non-negligeable deviation, especially towards the end of the simulation. As such, changes in the system environment reflecting into the protein charged state result in a significant increase in the conformational heterogeneity of DPF3a.

**Figure 7.**
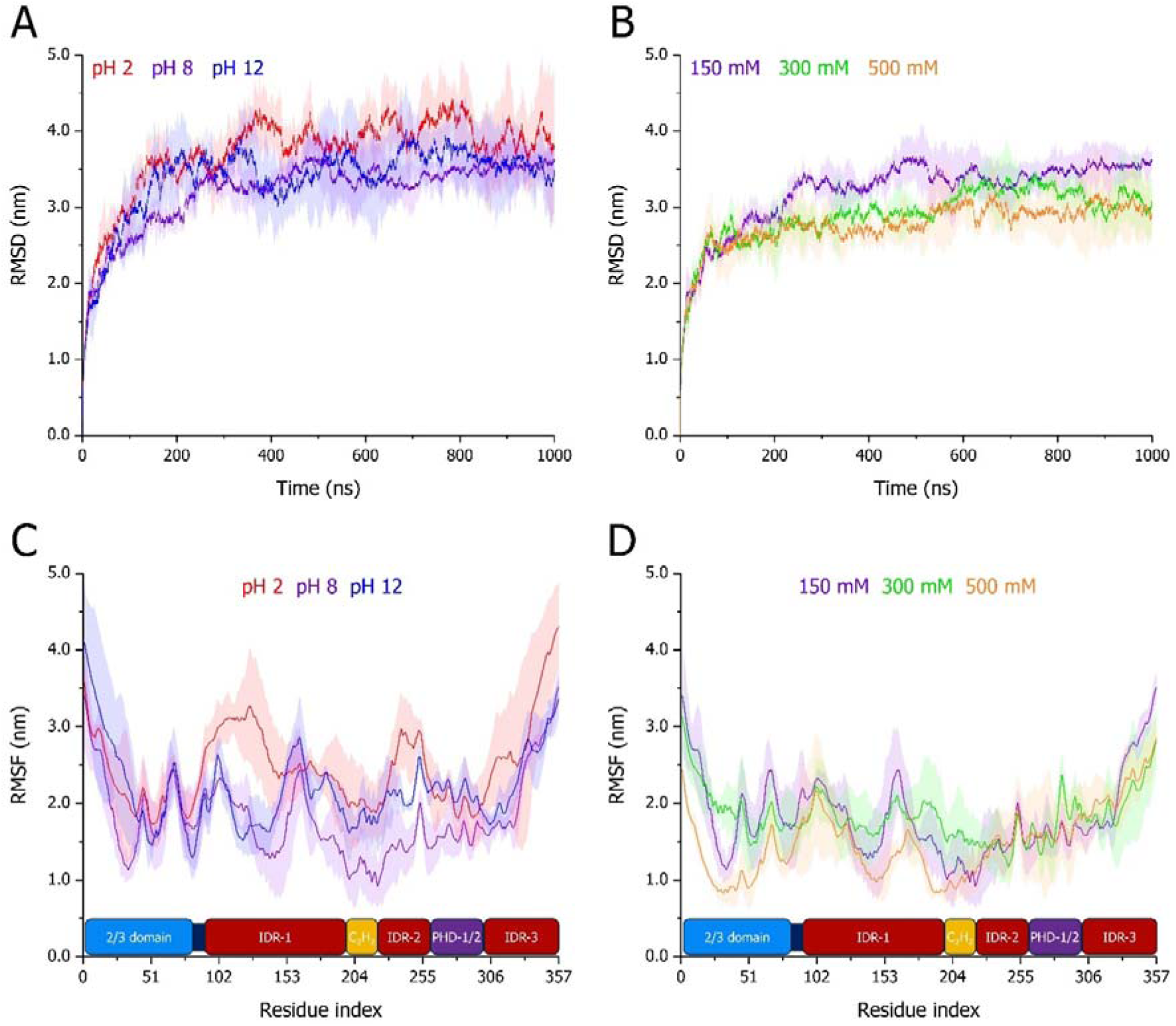
Time-evolution of (A-B) RMSD and (C-D) backbone RMSF of full-length DPF3a simulated for 1 µs at (A,C) pH 2 (red), 8 (purple), and 12 (blue) in 150 mM NaCl, as well as (B,D) in 150 (purple), 300 (green), and 500 mM NaCl (orange) at pH 8. For each condition, curves correspond to the average of triplicates with the standard deviation represented as a trace (shaded area) in the condition-associated colour. At the bottom of each RMSF plot, the sequence organisation of DPF3a is displayed according to its constitutive domains: the N-terminal 2/3 domain (blue), intrinsically disordered regions (dark red), the Krüppel-like C_2_H_2_ zinc finger (yellow), and truncated PHD-1/2 (purple).

Examination of the backbone atoms flexibility through the root-mean square fluctuation (RMSF) averaged over the complete simulation time frame shows that, i) unsurprisingly, identified IDRs display high RMSF values, especially towards the C-terminal region with a gradual increase in flexibility, and ii) the C_2_H_2_ ZnF is the most rigid domain due to the permanence of the fold stabilised by Zn^2+^ coordination (Fig. 7C). Interestingly, while the start of 2/3 domain (residue 1-40), predicted to have an α-helix fold, shows a decreased flexibility, its end, from residue 50 to 80, presents high RMSF, comparable to that of an IDR, suggesting that it might be disordered as well. This region is also far more sensitive to ionic strength, becoming more rigid at higher the salt concentrations (Fig. 7D). Upon acidification, every residue within DPF3a increases its flexibility, especially for the residues belonging to the three IDRs (Fig. 7C). Although to a lesser extent, higher RMSF values are also observed at pH 12, more specifically from IDR-1 to PHD-1/2, as well as for the first 50 N-terminal amino acids, which could evocate the unfolding of the N-terminal α-helix region. Regarding ionic strength, the variations are more subtle (Fig. 7D). Nonetheless, the RMSF profile at 300 mM is overall similar to that of 150 mM, while 500 mM NaCl leads, on average, to a reduction in the backbone flexibility. Most of IDRs are affected differently, although the C-terminal region presents less differences between the saline conditions. Significant deviation is observed for 300 and 500 mM at the level of the IDR-3 and the truncated PHD-1/2 domain, respectively. Therefore, while extreme pH, especially acidic, leads to a global increase of the protein chain flexibility, the effect of ionic strength manifests more locally with some regions presenting either higher or lower backbone flexibility depending on the salt concentration. Such tendency is in good agreement with the variations observed on the RMSD plots and is also well described by the protein compactness.

Indeed, the radius of gyration (R_g_) of DPF3a, whereas extending during the first 100 ns of the simulation, progressively converges to a low value at pH 8 (Fig. 8A, Table 1). This tendency is also evidenced by a relatively narrower R_g_ distribution comprised between 2.5 and 4.5 nm (Fig. 8B). Such behaviour is very coherent with the DPF3a sequence-embedded charge patterning (Fig. S1A), as medium-to-high κ proteins (κ_DPF3a_ = 0.27) tend to compact into low-R_g_ structures compared to low-κ IDPs [85,95]. The collapse of the protein structure is no longer observed at low and high pH (Fig. 8A, Table 1), and, on the contrary, the protein chain gradually but significantly expands over the course of time. Albeit the R_g_ value seemingly stabilises towards the end of the simulation under alkaline condition at an intermediate R_g_ value, both pH conditions exhibit particularly large deviation throughout the trajectory compared to pH 8. Accordingly, their associated R_g_ distributions are broadened and occupy a more heterogeneous and extended span, ranging from 3.0 to 6.5 nm (Fig. 8B). While at first a similar decrease in the radius of gyration can be observed at high ionic strength, the protein adopts a looser structure from 500 ns onwards, characterised by an increase in R_g_, as well as a dramatic variability between the replicates. This effect is most noticeable in 500 mM NaCl for which R_g_ values reache up to 5.0 nm (Fig. 8C, Table 1). Such conformational expansion in the second half of the simulation results in a shift of the distribution towards higher R_g_ values observed for 300 mM and even the population of very high-R_g_ conformations comprised between 4.0 and 5.5 nm for 500 mM (Fig. 8D). In this regard, extreme pH and increase of the ionic strength uncompact DPF3a structure, with charge repulsion, i.e. change in the protonation state, having larger effect than the screening of attractive electrostatic intramolecular interactions dependent on the ionic strength.

**Figure 8.**
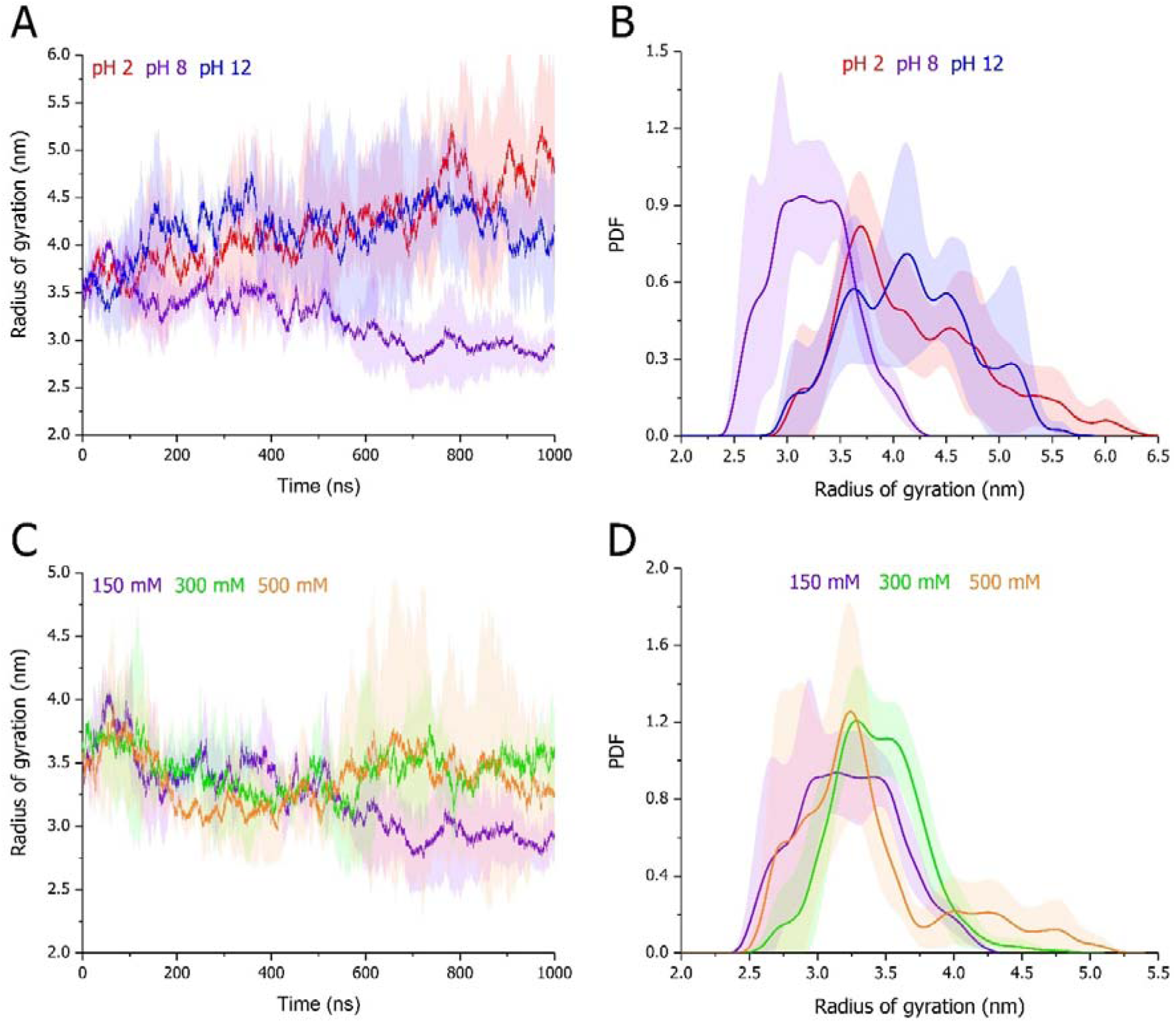
Radius of gyration (A,C) evolution over time and (B,D) distribution over the trajectories of full-length DPF3a simulated for 1 µs at (A-B) pH 2 (red), 8 (purple), and 12 (blue) in 150 mM NaCl, as well as (C-D) in 150 (purple), 300 (green), and 500 mM NaCl (orange) at pH 8. For each condition, curves correspond to the (A,C) time-evolution and (B,D) probability density function (PDF) average of triplicates with the standard deviation represented as a trace (shaded area) in the condition-associated colour.

**Table 1.** Mean values and associated standard deviation of MD variables extracted from the averaged trajectory over triplicates of full-length DPF3a simulated for 1 µs in selected pH and ionic strength conditions. Indicated standard deviation values correspond to that of the averaged sample. Percentages of secondary structure types were determined with the STRIDE algorithm. R_g_, radius of gyration; d_ee_, end-to-end distance; SASA, solvent accessible surface area; Contact, number of intramolecular contacts < 5 Å; CPA, number of intramolecular contacts per protein atom; H bond, number of intramolecular protein H bonds < 3.5 Å and 30° angle; H bond D, number of H bond donors; H bond A, number of H bond acceptors; Total D-A, total number of H bond donors and acceptors; H bond ratio, ratio of H bond against Total D-A.

|  | pH 2<br>150 mM NaCl | pH 8<br>150 mM NaCl | pH 12<br>150 mM NaCl | pH 8<br>300 mM NaCl | pH 8<br>500 mM NaCl |
| --- | --- | --- | --- | --- | --- |
| R<sub>g</sub> (nm) | 4.2 ± 0.4 | 3.2 ± 0.3 | 4.2 ± 0.3 | 3.4 ± 0.2 | 3.4 ± 0.2 |
| d<sub>ee</sub> (nm) | 5.9 ± 0.8 | 5.4 ± 0.8 | 5.5 ± 0.8 | 5.3 ± 0.8 | 5.2 ± 0.7 |
| SASA (nm<sup>2</sup>) | 357 ± 7 | 333 ± 16 | 353 ± 8 | 343 ± 13 | 338 ± 13 |
| Contact (.10<sup>3</sup>) | 185.4 ± 1.5 | 187.1 ± 3.1 | 181.0 ± 1.6 | 185.2 ± 2.2 | 186.6 ± 2.5 |
| CPA | 33.2 ± 0.3 | 33.9 ± 0.6 | 33.0 ± 0.3 | 33.5 ± 0.4 | 33.8 ± 0.5 |
| H bond | 139 ± 7 | 164 ± 7 | 140 ± 8 | 152 ± 6 | 157 ± 6 |
| H bond D | 622 | 558 | 558 | 558 | 558 |
| H bond A | 1104 | 1104 | 1104 | 1104 | 1104 |
| Total D-A | 1726 | 1662 | 1662 | 1662 | 1662 |
| H bond ratio | 0.081 ± 0.004 | 0.099 ± 0.004 | 0.084 ± 0.005 | 0.092 ± 0.004 | 0.094 ± 0.004 |
| Turn (%) | 33.0 ± 3.7 | 33.8 ± 3.3 | 37.1 ± 4.2 | 36.1 ± 3.9 | 36.0 ± 3.5 |
| β-sheet (%) | 3.9 ± 2.0 | 3.4 ± 1.4 | 1.8 ± 1.3 | 3.5 ± 1.4 | 4.1 ± 1.5 |
| β-bridge (%) | 1.3 ± 0.8 | 2.1 ± 0.7 | 1.6 ± 0.8 | 2.1 ± 0.9 | 1.8 ± 0.8 |
| α-helix (%) | 18.6 ± 4.0 | 16.7 ± 2.7 | 11.5 ± 3.9 | 14.9 ± 3.4 | 17.0 ± 2.7 |
| 3<sub>10</sub>-helix (%) | 2.8 ± 1.3 | 3.1 ± 1.4 | 4.9 ± 1.8 | 2.8 ± 1.4 | 3.2 ± 1.5 |
| π-helix (%) | 0.0 ± 0.0 | 0.0 ± 0.1 | 0.1 ± 0.3 | 0.2 ± 0.4 | 0.0 ± 0.1 |
| Coil (%) | 40.4 ± 4.2 | 40.9 ± 3.3 | 43.1 ± 4.2 | 40.4 ± 3.7 | 37.9 ± 3.1 |

Concordantly, DPF3a solvent accessible surface area (SASA) steadily decreases over time at pH 8, whilst it leads to a higher mean value and larger oscillations in acidic and alkaline conditions (Fig. 9A, Table 1). More precisely, SASA average value for acidic and alkaline pH conditions deviates significantly from the behaviour of the system at pH 8 from 250 ns onwards. This is even better evidenced on the SASA distribution plot, showing that conformations at pH 2 and pH 12 occupy a more solvated space than pH 8 (Fig. 9B). The extent of the solvent exposure is enhanced upon acidification, which is reflected by a narrower population exhibiting a maximum density near 363 nm^2^, with respect to a broad SASA distribution centred around 355 nm^2^ at pH 12. Less variations in the SASA profile over time can be observed upon increasing the ionic strength. Yet SASA values tend to increase with higher ionic strengths, especially in the second half of the simulation (Fig. 9C, Table 1). Consequently, high ionic strengths displace the SASA distributions to larger values, with two main subpopulations at around 340 and 358 nm^2^ for both 300 and 500 mM NaCl (Fig. 9D), in comparison with the major population centred at 325 nm^2^ for 150 mM NaCl. Furthermore, replicates are quite reproducible at 150 mM, whereas a greater variability characterises the time-evolution and density distribution of SASA at higher ionic strength. In good agreement with the R_g_ data, pH-induced (de)protonation of DPF3a shifts its conformational ensemble towards more swollen and hydrated structures, a fact which is attributed to the repulsion between positively or negatively charged residues at pH 2 and 12, respectively (Figs. S1B-C). To a lesser extent, increased ionic strength leads to a similar effect by screening intramolecular attractive interactions favouring collapsed conformers. This also agrees with the experimental spectra, not only showing an enrichment in random coil but also the adoption of less collapsed tertiary structures with solvent-exposed aromatic residues.

**Figure 9.**
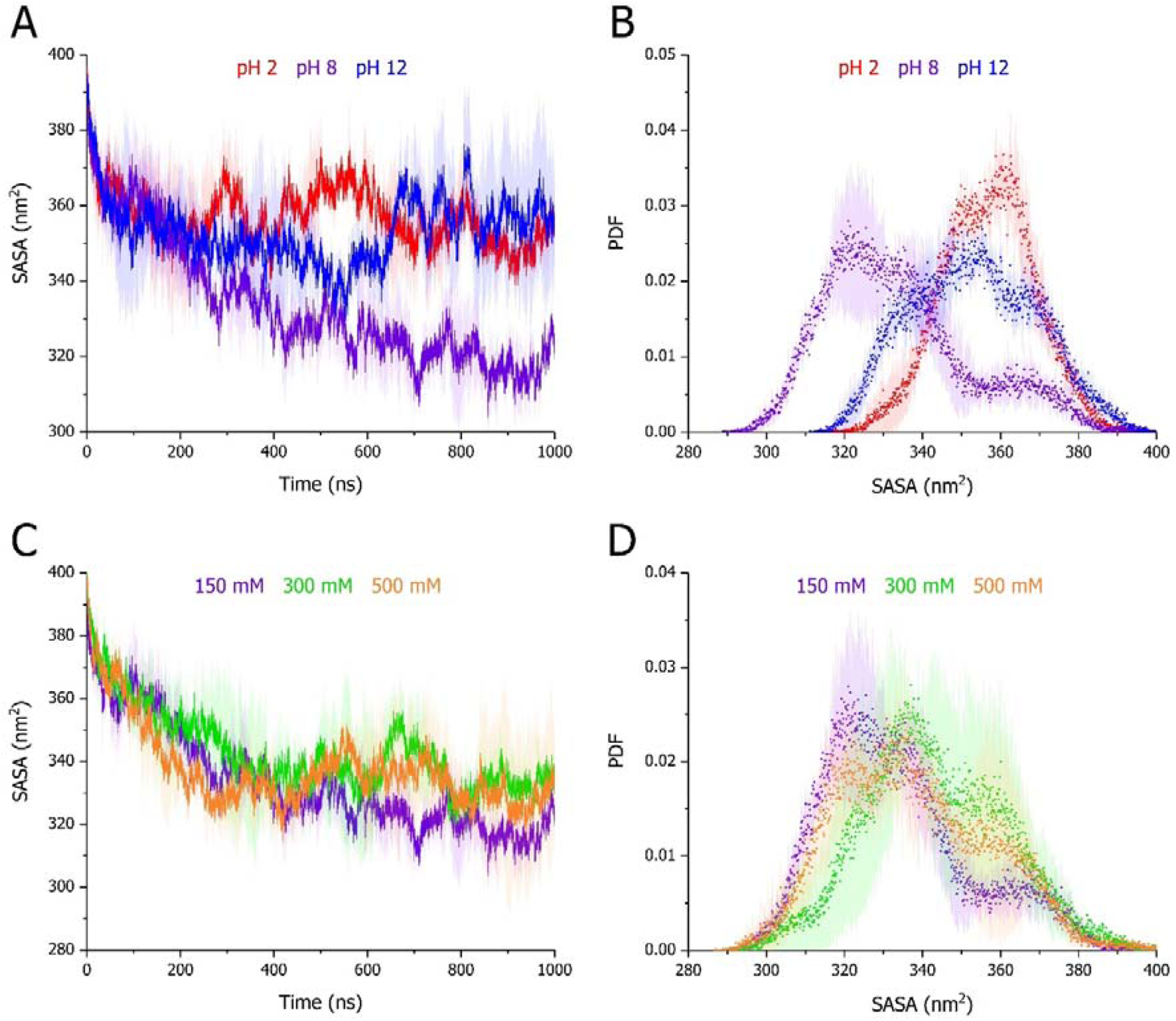
SASA (A,C) evolution over time and (B,D) distribution over the trajectories of full-length DPF3a simulated for 1 µs at (A-B) pH 2 (red), 8 (purple), and 12 (blue) in 150 mM NaCl, as well as (C-D) in 150 (purple), 300 (green), and 500 mM NaCl (orange) at pH 8. For each condition, curves and scattered dots respectively correspond to the (A,C) time-evolution and (B,D) probability density function (PDF) average of triplicates with the standard deviation represented as a trace (shaded area) in the condition-associated colour.

#### 3.4.2. Factors mediating the disruption of intramolecular interactions and secondary structures

As we hypothesise that the expansion of the protein chain is driven by, or results in, the loss of intramolecular interactions, the hydrogen bond (H bond) occurrence and number of contacts within DPF3a in the different tested conditions have been analysed. Predictably, the content in protein H bonds increases with time at pH 8 and 150 mM NaCl along the gradual compaction of its structure, amounting to a mean value of 164 H bonds over the trajectories (Fig. 10A, Table 1). By subjecting DPF3a to extreme pH, the proportion of intramolecular H bonds gradually decreases upon reaching an average value of ∼140 in both acidic and alkaline media (Fig. 10A-B, Table 1). Nonetheless, acidification-driven protonation leads to an increased number of H bond donor sites, thus introducing a population bias. Taking this into account and rationalising the number of H bonds with respect to the proportion of available donors and acceptors actually show that the overall H bond occupancy is higher at pH 12 than pH 2 (Table 1). To a lesser extent, high ionic strength also reduces the formation of H bonds in DPF3a, especially at the end of the simulations (Fig. 10C), which is even better illustrated by their density distribution broadening and shifting towards lower numbers (Fig. 10D). Globally, these tendencies are also well represented and confirmed by the number of intramolecular contacts defined by a 5 Å interatomic cut-off distance (Fig. S13, Table 1).

**Figure 10.**
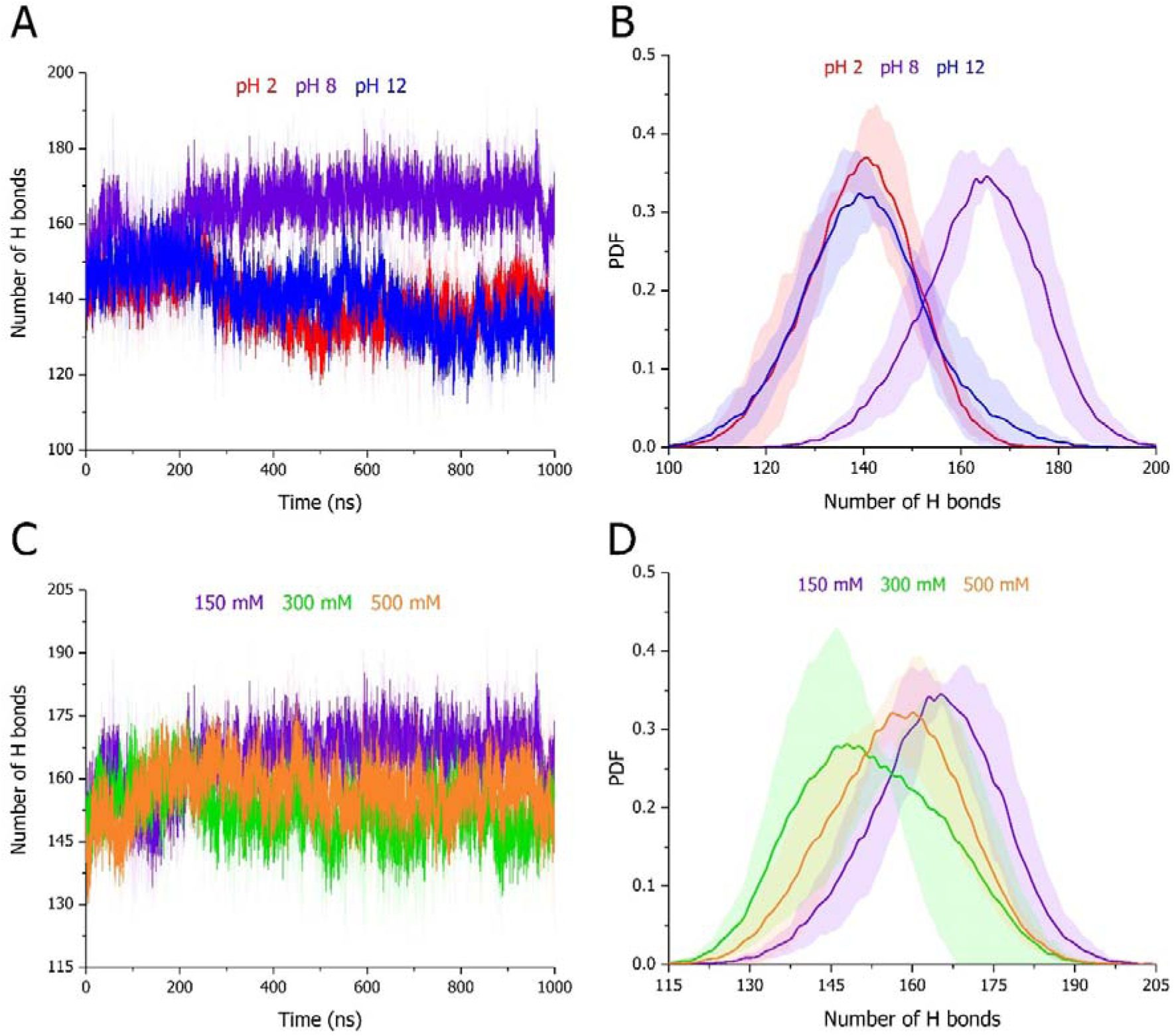
Intramolecular protein H bond (A,C) evolution over time and (B,D) distribution over the trajectories of full-length DPF3a simulated for 1 µs at (A-B) pH 2 (red), 8 (purple), and 12 (blue) in 150 mM NaCl, as well as (C-D) in 150 (purple), 300 (green), and 500 mM NaCl (orange) at pH 8. For each condition, curves correspond to the (A,C) time-evolution and (B,D) probability density function (PDF) average of triplicates with the standard deviation represented as a trace (shaded area) in the condition-associated colour.

Surprisingly, different trends are unveiled for the H bond evolution at the backbone level (Fig. S14). A reduction of the H bond population is observed for all conditions and is attributed to supplementary relaxation of the starting model and folded elements predicted by AlphaFold. The trajectory reaches a more stable phase ranging from 200 ns onwards at pH 8, while a further decrease is observed at both pH 2 and 12 before converging after 400 ns (Fig. S14A). Although the density distribution is centred around the same maximum value at pH 8 and 2, the latter displays two shoulders for lower and higher H bond numbers, respectively (Fig. S14B). Regarding the high-value population, it likely relates to a longer preservation of secondary structure motifs in acidic medium, especially within the first 300 ns of the dynamics. As evidenced by the time-evolution plot and by the lower density distribution at pH 12, the reduction in the backbone H bonds could be related to a greater loss in secondary structure elements. The latter tendency is also observable at pH 12, in line with the high RMSF detected for the N-terminal α-helix in the 2/3 domain for example. The decrease of H bond is achieved on a longer time period in 300 mM NaCl, while in 500 mM NaCl it reaches a plateau, even more rapidly compared to 150 mM, as reflected by a slight shift of the associated distribution towards lower and higher values, respectively (Figs. S14C-D). In this regard, H-bonded and folded structural domains may be (de)stabilised depending on the salt concentration.

As secondary structure elements are formed and stabilise through networks and defined geometries of H bonds engaging backbone amide groups, secondary structure propensities were assessed by analysing the trajectories with the STRIDE algorithm. Secondary structures were distinguished into seven classes: turns (predominantly β type, *i-i+3* turn), extended β-sheet, isolated β-bridge (usually *i-i+>6*), α-helix (*i-i+5* helix), 3_10_-helix (*i-i+3* helix), π-helix (distorted or bulged *i-i+5* helix), and random coil. No noticeable enrichment in β-sheet, β-bridge, or π-helix has been identified throughout the simulations. However, the time-evolution curves first indicate that the initial decrease in the backbone H bonds mainly arises from a loss in α-helical structures in each condition (Fig. 11), which is consistent with the analysis of the CD spectra. Although important variability is observed at pH 2, the helical content, on average, stabilises from 300 ns onwards (Fig. 11A). The helical content is instead further decreased at pH 8 (Fig. 11B), as well as at 300 (Fig. 11D) and 500 mM NaCl (Fig. 11E). The same effect is markedly more pronounced at pH 12, where it is concomitant with the formation of turns, 3_10_-helices, and coil (Fig. 11C). A global increase in coil also occurs at pH 2, 8, and in 300 mM NaCl, whereas turn enrichment is only promoted in alkaline and high ionic strength conditions. Yet, 500 mM does not enrich in coil and mainly favours the conversion of α-helices into turns (Table 1). Compared to pH 2 and 8, a higher proportion of turns is also detected at pH 12 and in 300 mM NaCl (Table 1). The different behaviour described for the backbone H bonds are coherent with the observed evolution in the secondary structures (Figs. 11). Such structural features could also connect to their comparable delaying effect on DPF3a fibrillation experimentally observed, as turn flexibility is a defining factor in the amyloidogenic propensity and mechanism of aggregation. Indeed, the introduction of more constrained or stronger H-bonded β-turns embedded in Aβ monomers has been demonstrated to significantly reduce or even inhibit their fibrillation [96].

**Figure 11.**
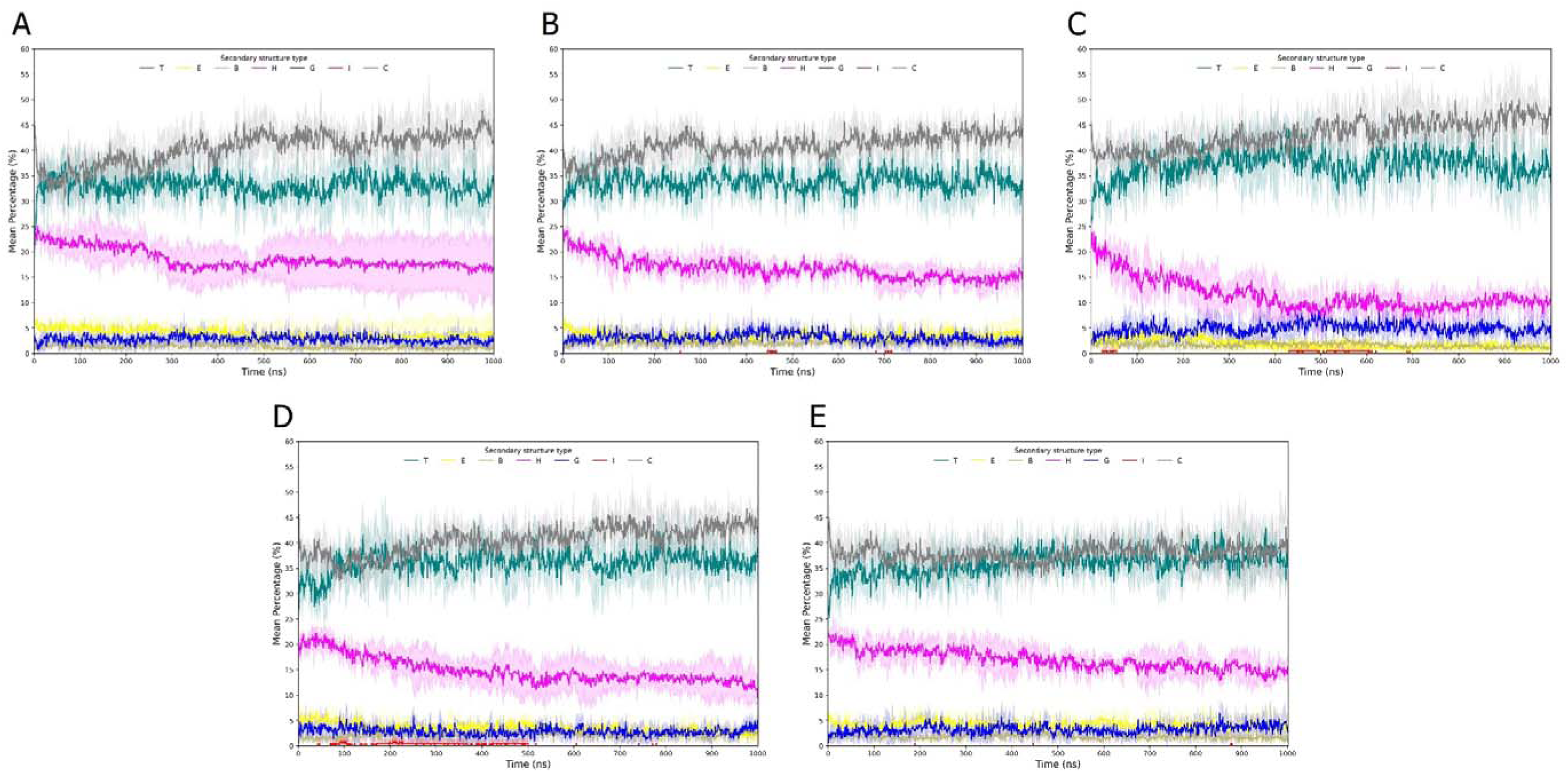
Time-evolution secondary structure content of full-length DPF3a simulated for 1 µs at (A) pH 2, (B) 8, and (C) 12 in 150 mM NaCl, as well as (D) in 300 and (E) 500 mM NaCl at pH 8. The secondary structures are defined into seven classes after STRIDE assignment: turn (T, turquoise), extended β-sheet (E, yellow), isolated β-bridge (B, tan), α-helix (H, pink), 3_10_-helix (G, blue), π-helix (I, red), and random coil (C, grey). For each condition, curves correspond to the average of triplicates with the standard deviation represented as a trace (shaded area) in the condition-associated colour.

By examining the secondary structure occurrence per-residue (Fig. S15), simulations at pH 12 illustrates that the conversion α-helices into turns and 3_10_-helices is predominantly localised in the 2/3 domain and IDR-1 more preponderantly (Fig. S15C). Comparatively, α-helical structures are more persistent in the other conditions, especially at pH 2 (Fig. S15A). The higher permanence of helical and coiled structures is likely unfavourable to the amyloid rearrangement of the protein in acidic condition. More turn occupancy is observed in the three IDRs at high ionic strength compared to 150 mM NaCl (Figs. S15D-E), the latter conserving a higher proportion of coil, especially towards the C-terminus, comprising PHD-1/2 and IDR-3 (Fig. S15B). In each condition the C_2_H_2_ ZnF maintains its characteristic Zn^2+^-coordinated α-helix and antiparallel β-sheet fold. Interestingly, β-sheet- and β-bridge-forming residues are mainly localised at the extremities of DPF3a sequence, that is, in the second half of the N-terminal 2/3 domain, as well as the C-terminal segment spanning from the truncated PHD-1/2 domain to the IDR-3, which presumably correspond to aggregation or amyloid prone regions concordantly with our previous sequence-based predictions [79]. Last but not least, increased β-bridge frequency is only observed for the referential condition in IDR-3, advocating for its higher intramolecular contact propensity, which favours structure collapsing and amyloid conversion.

#### 3.4.3. Environmental dependence of DPF3a conformational landscape

To get an overview of the conformational ensembles adopted by DPF3a, trajectories were examined through plotting the occurrence of R_g_ and end-to-end distance (d_ee_) pairs into Kernel density maps, as well as clustering analysis using a RMSD cut-off of 5 Å between neighbouring structures. First, the R_g_-d_ee_ space occupied by DPF3a is revealed to be very responsive to the physicochemical environmental conditions (Fig. 12). In the reference condition, while the protein is able to visit moderately swollen conformations, the most populated conformational space corresponds to very collapsed and compact structures of low R_g_ and variable d_ee_ values. Upon acidification, the conformational density is not only shifted towards both higher R_g_ and d_ee_, but also spread over a broader range of populated conformations, which are represented by particularly swollen and expanded structures, notably through the extension of the C-terminal region. In comparison, DPF3a also displays more expanded conformations in alkaline condition, albeit some low density collapsed populations are still detected. Nevertheless, the overall landscape is far less diffuse at pH 2, as highlighted by the absence of a very high R_g_ subspace due to partial compaction within the C-terminal domain. In high ionic strength, the protein exhibits a relatively similar conformational landscape between 300 and 500 mM NaCl. It is characterised by a smaller space limited to a narrow R_g_ window of differently collapsed populations, between which the transitions occur through local swelling and/or termini distancing. Although conformationally more restrained in high salt concentration, DPF3a does not populate conformers as compact as those overrepresented in 150 mM NaCl. Coherently with our experimental data, whereas chain extension at pH 2 does not allow the protein to fold into amyloid precursors, the structure swelling observed in the other conditions is less favourable to undergo fibrillation, slowing down the rearrangement and nucleation steps.

**Figure 12.**
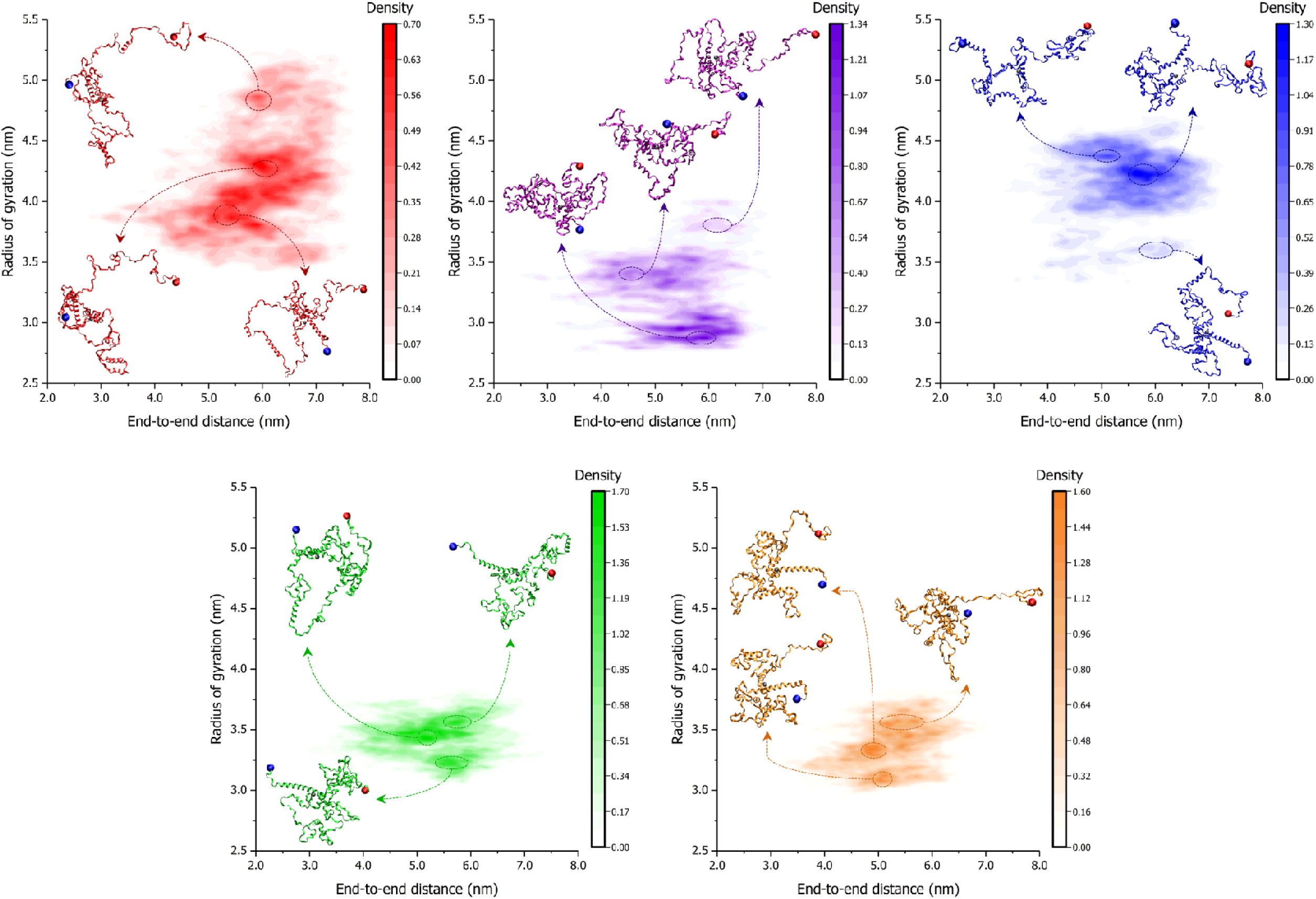
Conformational space of full-length DPF3a simulated for 1 µs at pH 2 (red, upper left), 8 (purple, middle), and 12 (blue, upper right) in 150 mM NaCl, as well as in 300 (green bottom left) and 500 mM NaCl (orange, bottom right) at pH 8. For each condition, radius of gyration and end-to-end distance pairs were averaged between the triplicates and represented on a two-dimensional Kernel density map in the condition-associated colour. On each panel, relevant conformational subspaces are highlighted by dashed ellipses and illustrated by a representative snapshot of the modelled protein structure shown as ribbon in the condition-associated colour with the N- and C-termini pinpointed as blue and red spheres, respectively. The coordinated Zn^2+^ cation is represented as a grey bead.

Through the clustering dimensionality reduction, the previously analysed tendencies are once more corroborated. RMSD distribution profiles show that variation of pH increases the deviation from the initial structure (Fig. S16A) and the conformational heterogeneity along the simulation, as barely any largely populated cluster can be observed compared to pH 8 (Fig. S16C). While the distribution is also shifted towards higher RMSD values in 300 mM NaCl, the same maximum is observed for 500 mM (Fig. S16B). Compared to pH, both ionic strength conditions converge better, but do not stabilise into a well-defined cluster upon the end of the simulation and exhibit significant variability all along the trajectory, contrasting with the consistency between the replicates observed at 150 mM (Fig. S16D). For each simulated system, the minimum distance contact map of the central structure corresponding to the most populated cluster amongst the triplicated trajectories was extracted and compared with respect to pH 8 in 150 mM NaCl (Fig. 13). In the latter, DPF3a adopts a very compact state, in which every domain interacts with the others by establishing multiple and multivalent interfaces thanks to the structure collapse. Because of chain swelling and C-terminal extension, acidification overall leads to a loss in inter-IDR communication through long-range interactions, especially between IDR-1 and IDR-2, as well as IDR-2 and IDR-3 (Fig. 13A). A comparable trend is observed at pH 12, with the 2/3 domain, PHD-1/2, and IDR-3 being isolated, even though most of the new contacts are mediated at the level of the C_2_H_2_ ZnF, around which the protein tends to gain in compactness (Fig. 13B). In this aspect, DPF3a presents a higher degree of compaction in 300 mM NaCl, yet less short- and long-range interactions engaging the 2/3 domain and IDR-1 can be observed, the contacts being concentrated in and between the C-terminal domains (Fig. 13C). While IDR-1 losing inter-domain communication is once more the most affected, a slightly higher proportion of interactions between the 2/3 domain with IDR-2 and PHD-1/2, as well as of intra-IDR-1 contacts is found in 500 mM NaCl (Fig. 13D). The loss of long-range interactions is also very evocative upon comparison of the final state of each simulation (Fig. S17). From pH 2 (Fig. S17A) to 500 mM NaCl (Fig. S17D), DPF3a progressively transitions from a very expanded conformer to a more collapsed but swollen structure. At pH 2, only few contacts between the 2/3 domain with C_2_H_2_ and IDR-2 are observed, which is unsurprising given the rod-like extension of the PHD-1/2 and IDR-3 domains (Fig. S17A). In all the conditions, contacts with IDR-1 are mostly inexistent, while at pH 8 the IDR has multivalent interactions throughout the C-terminal segment, comprising IDR-2, PHD-1/2, and IDR-3 (Figs. S17A-D).

**Figure 13.**
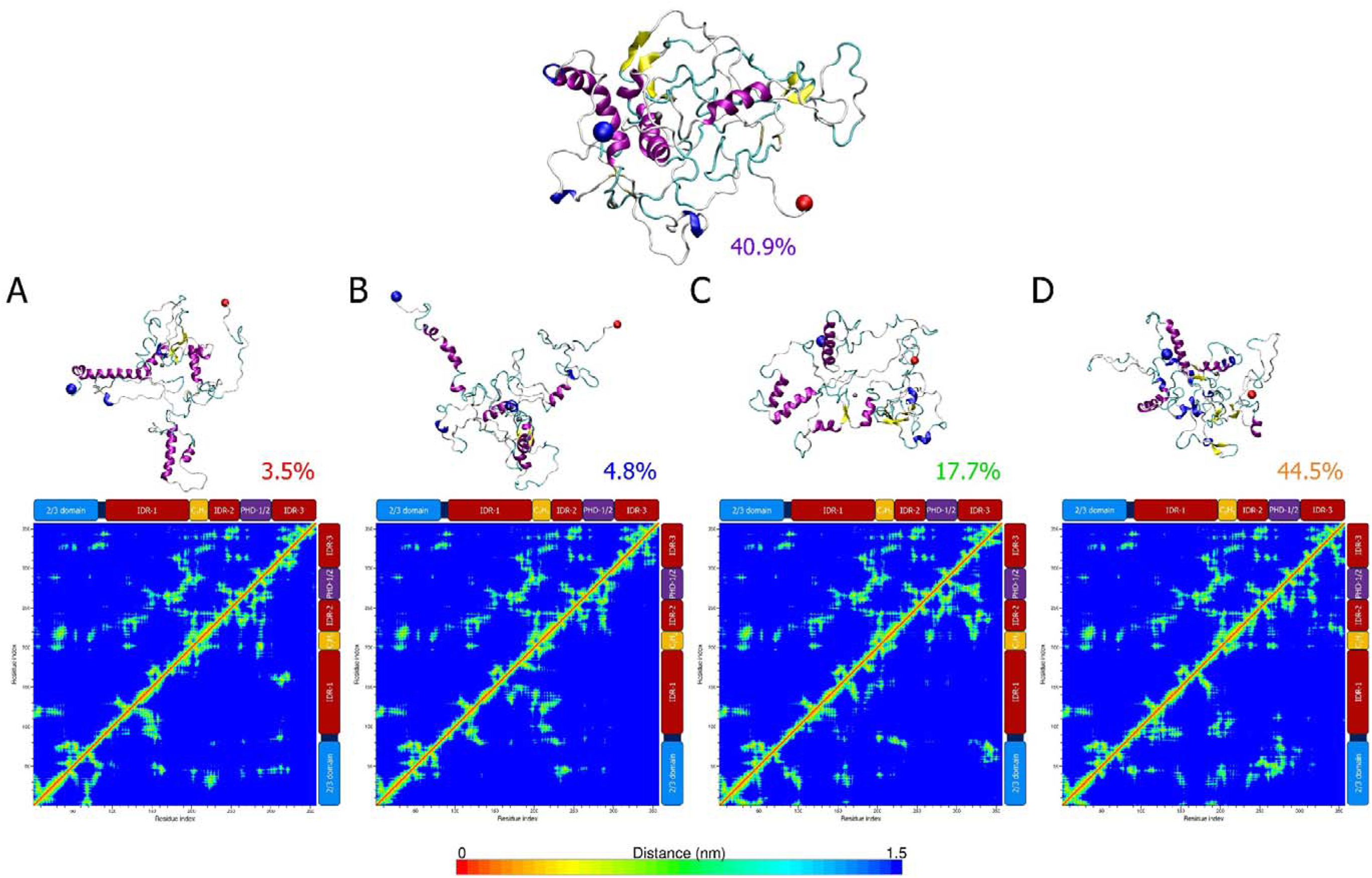
Conformational clustering analysis and residue pairs contact mapping of full-length DPF3a simulated for 1 µs at (A) pH 2, 8 (upper panel), and (B) 12 in 150 mM NaCl, as well as in (C) 300 and (D) 500 mM NaCl at pH 8. For each condition, the central protein structure of the most populated cluster amongst the triplicates (upper row) is shown in cartoon representation with the N- and C-termini respectively pinpointed as blue and red spheres, and the different secondary structure elements coloured according to STRIDE assignment: turn (turquoise), extended β-sheet (yellow), isolated β-bridge (tan), α-helix (pink), 3_10_-helix (blue), π-helix (red), and random coil (light grey). The coordinated Zn^2+^ cation is represented as a grey bead. Percentages indicate the population size of the first cluster over the selected trajectory. Minimum distance contact maps between residue pairs (bottom row) are compared between the different pH and ionic strength conditions (bottom half on each map) with respect to the pH 8 and 150 mM NaCl system taken as a reference (upper half on each map). On the sides of each contact map, the sequence organisation of DPF3a is displayed according to its constitutive domains: the N-terminal 2/3 domain (blue), intrinsically disordered regions (dark red), the Krüppel-like C_2_H_2_ zinc finger (yellow), and truncated PHD-1/2 (purple).

Because of the protonation of aspartate (Asp) and glutamate (Glu) carboxyl groups at pH 2, the protein acts more like a positively charged polyelectrolyte IDP, adopting swollen coil and extended conformers due to intramolecular repulsive interactions between positively charged residues, i.e. lysine (Lys) and Arg residues. This is well evidenced by the net charge per-residue distribution of DPF3a upon abrogating the negative charge of its Asp and Glu amino acids (Fig. S1B). In addition, intermolecular contacts between monomers are also inhibited due to charge repulsion, suppressing DPF3a amyloid propensity to rearrange and self-assemble into fibrillar aggregates. In this regard, high ionic strength could help recovering DPF3a ability to form amyloids, as ions will screen short- and long-range repulsive interactions, increasing the overall protein compactness and protein-protein affinity. Conversely, at pH 12, the protein tends to behave like a negatively charged polyelectrolyte IDP, but the permanence of the Arg positive charge partly compensates the repulsion between the negative charges, as exemplified by the net charge per-residue distribution (Fig. S1C). Therefore, while leading to an overall expansion of DPF3a conformational ensemble, alkaline conditions still allow the protein structure to collapse through attraction between patches of opposite charges. Such charge unbalance, driven by Arg compensation, while delaying the fibrillation process, does not prevent it from taking place. As such, it is expected that further alkalinisation, resulting in Arg deprotonation, will inhibit DPF3a amyloidogenic proneness, in way comparable to that of pH 2. Furthermore, given the higher-than-average content of Cys residues in the DPF3a sequence, alkaline-enhancing thiolate reactivity and disulphide scrambling could promote pro-amyloid structural rearrangement and association between protein molecules through disulphide bridging. In this intermolecular hypothesis, thiolate moieties need to be surface accessible and distanced from one another to avoid intramolecular bridges. Such requirements are *de facto* fulfilled at pH 12, simulations showing that Cys residues are significantly more solvent-exposed and away from each other vis-à-vis pH 8 (Fig. S18).

Although with a less pronounced effect, high ionic strength favours more swollen conformers with less contacts within the protein through screening attractive interactions between the scattered charged patches distributed amongst the different protein domains and IDRs. Comparatively, in the reference condition (pH 8 in 150 mM NaCl), DPF3a structure collapses via favourable long-range inter-domain/IDRs interactions, as well as by fostering intramolecular H bond networking and secondary structure formation into amyloidogenic conformations.

## 4. Conclusions

In the present study, we explored and deciphered the effect of pH and ionic strength on the structural and fibrillation properties of the amyloidogenic IDP DPF3a, by combining spectroscopy, microscopy, and molecular dynamics approaches. In a nutshell, we found out that DPF3a is a polyampholyte IDP that is highly responsive to variation in solution conditions, thus acting as an environmental sensor, which is perfectly in line with our previously discussed predictions categorising it as a context-dependent or Janus sequence [32]. Furthermore, we underlined that such adaptive behaviour arises from the sequence-embedded charge distribution and amino acid composition of DPF3a, parameters that have concordantly been identified as key tuners of aggregation and phase separation in other protein systems [95,97].

More detailedly, the protein amyloidogenic pathway was integrally inhibited upon extreme acidification, which was unveiled through a very low proportion of Trp-Tyr FRET-enabling conformers, highly solvent-exposed and associated to the permanence of disorder-turn conformation, as well as through the absence of photoactive fibrillar species (Fig. 14). MD data validated such results, showing that a highly protonated DPF3a is indeed enriched in coil elements, displays a low number of intramolecular H bonds and inter-domain contacts, and remarkably occupies an expanded and heterogeneous conformational space. The latter is mainly characterised by a rod-like extension of the C-terminal region, a globally elevated backbone flexibility, as well as high R_g_ and SASA values. In the light of positive charge enrichment at pH 2, such patterns were attributed to substantial intra- and intermolecular repulsive interactions, the protein behaving similarly to a polyelectrolyte IDP.

**Figure 14.**
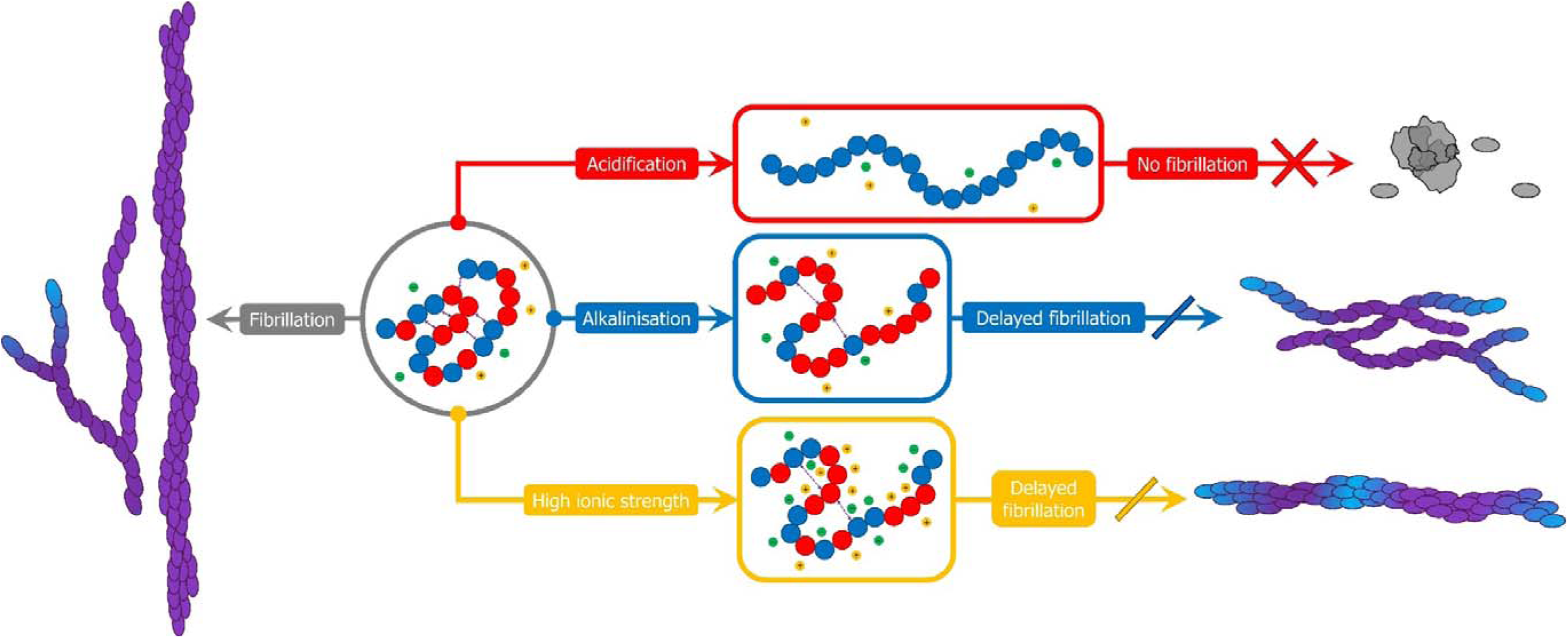
Modulation of the conformational preference and amyloidogenic pathway of the intrinsically disordered polyampholyte protein DPF3a through sensing physicochemical changes in its environment. (Grey route) In a physiological-like context, its charge distribution and sequence composition allow DPF3a to structurally rearrange by favourable chain collapse, as well as to self-assemble into long, mature, and vAF-emitting amyloid fibrils, resulting from the association of dbAF-emitting (proto)fibrillar intermediates. (Red route) Due to protonation-mediated abrogation of its negative charges, DPF3a harbours a polyelectrolytic behaviour in acidic medium, characterised by highly hydrated and extended conformations that are unable to fibrillate and sporadically cluster into photoinactive amorphous phases. (Blue route) Upon alkalinisation, DPF3a structure compaction and aggregation prone contacts are reduced due to repulsion between negatively charged residues, favouring dbAF prefibrillar emitters and impairing the fibrillation process, which is nonetheless rescued thanks to partial charge compensation by Arg sidechains and enhanced thiolate reactivity. (Yellow route) Under increased ionic strength, β-sheet conversion and fibril elongation steps become hindered through the screening of attractive intra- and intermolecular electrostatic interactions, maintaining DPF3a in a more swollen state and leading to the generation of short fibrils containing both vAF and dbAF fluorophores.

While not as impaired, DPF3a fibrillation process was delayed in alkaline condition, through the formation of less β-sheeted and more hydrated intermediates during the first steps of the protein amyloid conversion (Fig. 14). Furthermore, alkalinisation fostered the generation of dbAF-emitting populations, which were associated to thin and cobweb-like fibrous structures later maturing and assembling into fibrils fluorescing in the vAF region. Simulations confirmed that a higher negative net charge, via the deprotonation of Lys amino acids, resulted in more solvated and less compact conformers, presenting higher flexibility at the level of IDRs, as well as fewer intramolecular contacts over the sampled microsecond timescale. Nevertheless, the amyloid susceptibility was rescued at pH 12 not only thanks to partial charge compensation with positively charged Arg residues, leading to a more homogeneous conformational landscape, but also through the enhanced reactivity of thiolate moieties, the accessibility of which was ascertained by our simulations, thus likely enabling pro-amyloid disulphide bridging between monomers.

At high ionic strength, β-sheet conversion and fibrillation were also slowed down through the retention of swollen conformations, as well as the formation of constrained β-turns, which are less favourable to amyloid nucleation and elongation, resulting in a lower proportion of mature fibrils characterised by shorter length and less efficient intrinsic vAF-emitters (Fig. 14). Self-assembly hindrance was assigned to the screening effect of ions with respect to the distribution of charged residues in DPF3a sequence. Indeed, rather than extensively screening intramolecular contacts in 300 mM NaCl, ions could partially shield interactions between DPF3a molecules that are required to the assembly into oligomers and fibrils. At even higher ionic strength, the extent of the effect is exacerbated to the point of impairing attractive interactions within the protein, which consequently adopts a more expanded conformational ensemble, further delaying the generation of β-sheeted species. Such claims were substantiated by the MD simulations, highlighting that increasing the ionic strength induced more conformational heterogeneity, greater variability between replicated trajectories, structural swelling, as well as loss of long-range contacts between N- and C-terminal regions.

As both pH and ionic strength were proposed as defining factors in the disruption of intermolecular interactions between DPF3a molecules for explaining the delay in amyloid nucleation and fibrillation, such properties could also be investigated by MD in future studies, either exploiting coarse-grained models on several copies of the full-length protein or performing classical all-atom simulations on selected domains and aggregation prone regions. As the druggability of amyloidogenic IDPs is intricate by essence, by shedding light onto the factor-inducible conformational transitions of DPF3a, we hope opening new avenues for the development of innovative therapeutic strategies to target it in cancer and neurodegenerative diseases.

Altogether, our results unveiled that DPF3a is a context-dependent polyampholyte IDP the amyloidogenic pathway and propensity of which can be modulated by modifications of its physicochemical environment, shaping the protein conformational ensemble and optical-morphological properties via electrostatic attractive-repulsive interactions and screening effects with respect to the unique distribution of charge and amino acids encoded in its sequence.

## Abbreviations

APR: aggregation prone region
α-syn: α-synuclein
AD: Alzheimer’s disease
Aβ: amyloid β peptide
AF: autofluorescence
BAF: BRM/BRG1-associated factor
bgAF: blue-green autofluorescence
CD: circular dichroism
CMAP: correction map
dbAF: deep-blue autofluorescence
DPF3: double PHD fingers 3
d_ee_: end-to-end distance
EEM: excitation-emission matrix
HMR: hydrogen mass repartitioning
FRET: Förster resonance energy transfer
IDP: intrinsically disordered protein
IDR: intrinsically disordered region
IPF: intrinsic phenylalanine fluorescence
ITF: intrinsic tryptophan fluorescence
ITyrF: intrinsic tyrosine fluorescence
MD: molecular dynamics
PD: Parkinson’s disease
PBS: phosphate-buffered saline
PMT: photomultiplier tube
PHD: plant homeodomain
PDF: probability density function
R_g_: radius of gyration
RMSD: root-mean square deviation
RMSF: root-mean square fluctuation
sw: slit width
SASA: solvent accessible surface area
TOF: Tetralogy of Fallot
TEM: transmission electron microscopy
TBS: Tris-buffered saline
vAF: violet autofluorescence
ZnF: zinc finger.

## Data availability

Data will be made available from the corresponding author upon request.

## Acknowledgements and funding

The authors are grateful to the Research Unit in Biology of Microorganisms (URBM), as well as to the L.O.S. and Morph-Im platforms of the University of Namur. The authors are also appreciative of the PTCI high-performance computing resource of the University of Namur. This work benefited from computational resources provided by the Consortium des Équipements de Calcul Intensif (CÉCI), funded by the Belgian National Fund for Scientific Research (F.R.S.-FNRS) under grant n°2.5020.11 and by the Walloon Region, and made available on Lucia, the Tier-1 supercomputer of the Walloon Region, infrastructure funded by the Walloon Region under the grant agreement n°1910247. J. M. and T. L. thank the FNRS for their Research Fellow fellowship and Fund for Research training in Industry and Agriculture (FRIA) Doctoral grant, respectively. A. M. thanks ANR and CGI for their financial support of the present work through Labex SEAM ANR 11 LABEX 086, ANR 11 IDEX 05 02. The support of the IdEx “Université Paris 2019” ANR-18-IDEX-0001 is also acknowledged. D. M. and C. M. thank the FNRS for their Senior Research Associate position. This research did not receive any specific grant from funding agencies in the public, commercial, or not-for-profit sectors.

## CRediT authorship contributions

Julien Mignon: Conceptualisation; Methodology; Validation; Formal analysis; Investigation; Data acquisition and interpretation; Visualisation; Writing-original draft; Writing-reviewing & editing.

Tanguy Leyder: Validation; Writing-reviewing & editing.

Antonio Monari: Conceptualisation; Validation; Data interpretation; Writing-reviewing & editing.

Denis Mottet: Validation; Writing-reviewing & editing.

Catherine Michaux: Conceptualisation; Validation; Supervision; Writing-reviewing & editing.

## Author information

Julien Mignon

Tanguy Leyder

Antonio Monari

Denis Mottet

Catherine Michaux

