## Supporting Information for "Exploration of the influence of environmental changes on the conformational and amyloidogenic landscapes of the zinc finger protein DPF3a by combining biophysical and molecular dynamics approaches"

**–**

|  | pH 2  150 mM NaCl | pH 8  150 mM NaCl | pH 12  150 mM NaCl | pH 8  300 mM NaCl | pH 8  500 mM NaCl |
| --- | --- | --- | --- | --- | --- |
| Protein net charge | +59 | −5 | −37 | −5 | −5 |
| Box volume (nm^3^) | 2049.1 | 1751.4 | 1793.5 | 1751.4 | 1751.4 |
| Water molecule | 58908 | 50197 | 51376 | 49881 | 49459 |
| Water atom | 235632 | 200788 | 205504 | 199524 | 197836 |
| Na^+^ | 185 | 163 | 199 | 321 | 532 |
| Cl^−^ | 244 | 158 | 162 | 316 | 527 |
| Zn^2+^ | 1 | 1 | 1 | 1 | 1 |
| Ions | 430 | 322 | 362 | 638 | 1060 |
| Protein atom | 5583 | 5519 | 5487 | 5519 | 5519 |
| Total atom | 241645 | 206629 | 211353 | 205681 | 204415 |
| Ionic strength (M) | 0.18 | 0.15 | 0.17 | 0.30 | 0.50 |
| Minimisation (ns) | 3.654 | 2.637 | 3.207 | 3.298 | 3.552 |
| NVT equilibration (ns) | 0.200 | 0.200 | 0.200 | 0.200 | 0.200 |
| NPT equilibration (ns) | 0.200 | 0.200 | 0.200 | 0.200 | 0.200 |
| Total production MD (ns) | 3000 | 3000 | 3000 | 3000 | 3000 |
| Total simulation time (µs) | 15.02 | | | | |

Table S1 – Summary of the molecular dynamics metrics, parameters, and durations for the five simulated pH and ionic strength systems of full-length DPF3a protein.

|  |  | Secondary structure content (%) | | | | |  |
| --- | --- | --- | --- | --- | --- | --- | --- |
|  | Time (h) | α-helix | Antiparallel  β-sheet | Parallel  β-sheet | Turn | Coil | RMSD |
| pH 2  150 mM NaCl | 0 | 0.0 | 34.6 | 0.0 | 18.3 | 47.1 | 0.048 |
|  | 24 | 0.0 | 36.2 | 0.0 | 17.6 | 46.2 | 0.043 |
|  | 48 | 0.0 | 34.9 | 0.0 | 17.8 | 47.3 | 0.041 |
|  | 72 | 0.0 | 36.8 | 0.0 | 16.8 | 46.4 | 0.038 |
|  | 96 | 0.0 | 40.3 | 0.0 | 16.2 | 43.6 | 0.047 |
|  | 168 | 0.7 | 35.3 | 0.0 | 16.5 | 47.6 | 0.029 |
| pH 8  150 mM NaCl | 0 | 4.8 | 30.1 | 0.4 | 13.4 | 51.3 | 0.076 |
|  | 24 | 3.1 | 34.6 | 0.0 | 13.6 | 48.8 | 0.059 |
|  | 48 | 2.6 | 36.5 | 0.0 | 14.7 | 46.3 | 0.048 |
|  | 72 | 2.1 | 37.2 | 0.0 | 14.9 | 45.8 | 0.051 |
|  | 96 | 2.1 | 37.4 | 0.0 | 15.8 | 44.7 | 0.022 |
|  | 168 | 0.7 | 37.2 | 0.0 | 15.7 | 46.4 | 0.019 |
| pH 12  150 mM NaCl | 0 | 4.5 | 32.2 | 0.0 | 16.5 | 46.9 | 0.073 |
|  | 24 | 4.5 | 31.0 | 0.0 | 15.9 | 48.6 | 0.061 |
|  | 48 | 3.5 | 32.6 | 0.0 | 14.6 | 49.3 | 0.049 |
|  | 72 | 3.0 | 33.6 | 0.0 | 14.0 | 49.4 | 0.048 |
|  | 96 | 2.6 | 36.3 | 0.0 | 14.5 | 46.6 | 0.053 |
|  | 168 | 2.2 | 37.1 | 0.0 | 15.0 | 45.8 | 0.048 |
| pH 8  300 mM NaCl | 0 | 5.1 | 30.1 | 0.0 | 13.2 | 51.7 | 0.086 |
|  | 24 | 3.4 | 33.1 | 0.0 | 13.6 | 49.9 | 0.053 |
|  | 48 | 3.3 | 32.9 | 0.0 | 13.2 | 50.7 | 0.055 |
|  | 72 | 2.9 | 33.9 | 0.0 | 13.8 | 49.4 | 0.058 |
|  | 96 | 2.4 | 34.8 | 0.0 | 14.5 | 48.3 | 0.062 |
|  | 168 | 2.6 | 35.0 | 0.0 | 13.6 | 48.8 | 0.059 |
| pH 8  500 mM NaCl | 0 | 5.2 | 29.6 | 0.1 | 14.2 | 50.9 | 0.080 |
|  | 24 | 4.1 | 31.4 | 0.0 | 13.3 | 51.3 | 0.060 |
|  | 48 | 3.3 | 33.2 | 0.0 | 13.1 | 50.4 | 0.057 |
|  | 72 | 2.8 | 34.3 | 0.0 | 13.9 | 49.0 | 0.053 |
|  | 96 | 2.7 | 34.5 | 0.0 | 14.3 | 48.6 | 0.057 |
|  | 168 | 2.4 | 34.4 | 0.0 | 14.4 | 48.8 | 0.046 |

Table S2 – BeStSel secondary structure content estimations and fit-associated RMSD values of full-length DPF3a incubated in selected pH and ionic strength conditions at ~25 °C for 0, 24, 48, 72, 96, and 168 h.

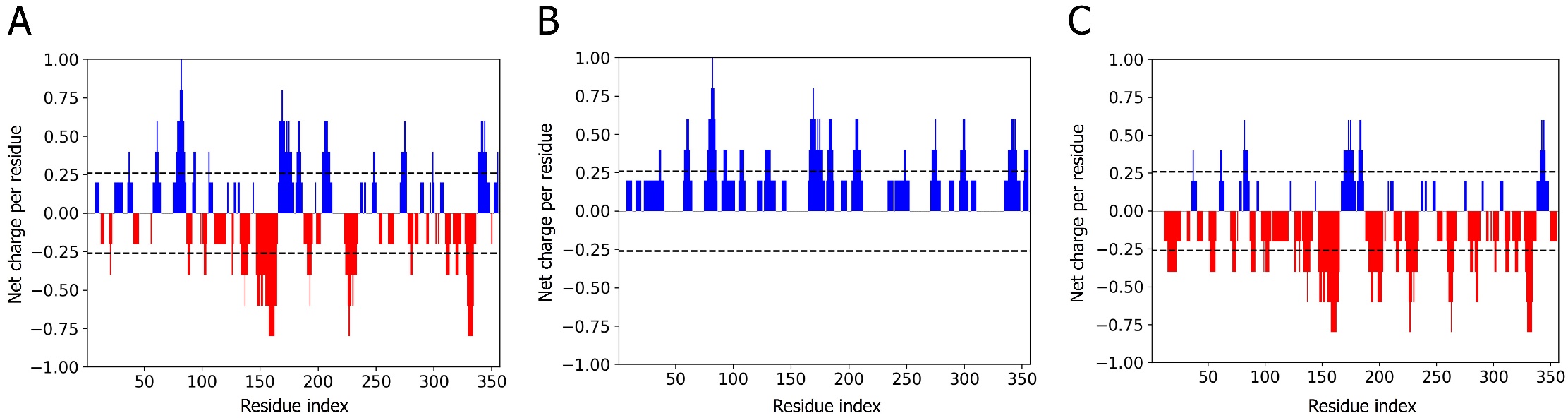

Figure S1 – Linear distribution of the net charge per residue of full-length DPF3a defined over a sliding window of five residues at (A) pH 8, (B) 2, and (C) 12. Data were generated and plotted using the CIDER (Classification of Intrinsically Disordered Ensemble Regions) web server.

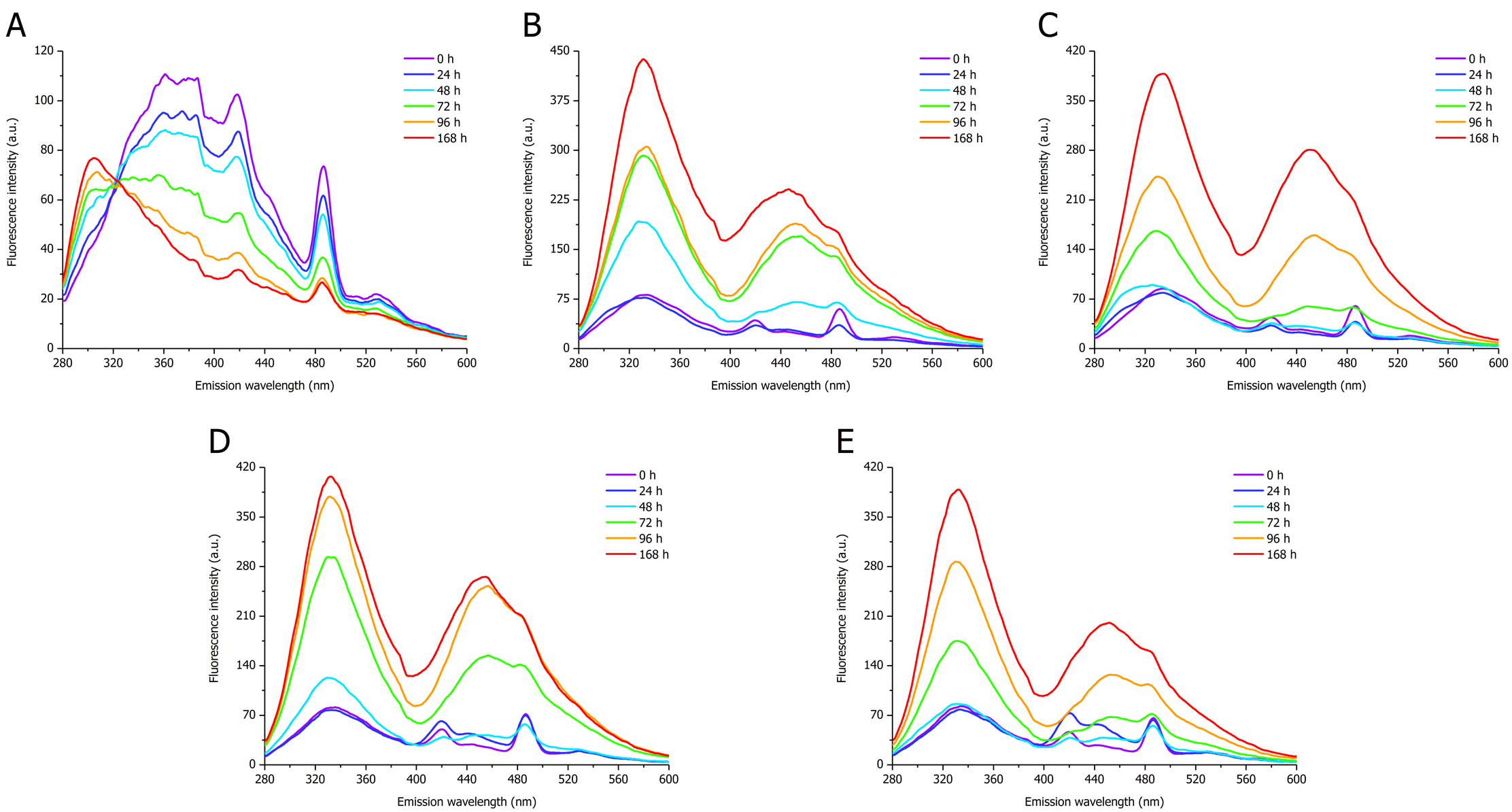

Figure S2 – Intrinsic IPF spectra (λ_ex_ = 260 nm, sw = 10 nm) of full-length DPF3a incubated at ~25 °C and (A) pH 2, (B) 8, and (C) 12 in 150 mM NaCl, as well as (D) 300 and (E) 500 mM NaCl at pH 8 for 0 h (purple), 24 h (dark blue), 48 h (light blue), 72 h (green), 96 h (orange), and 168 h (red).

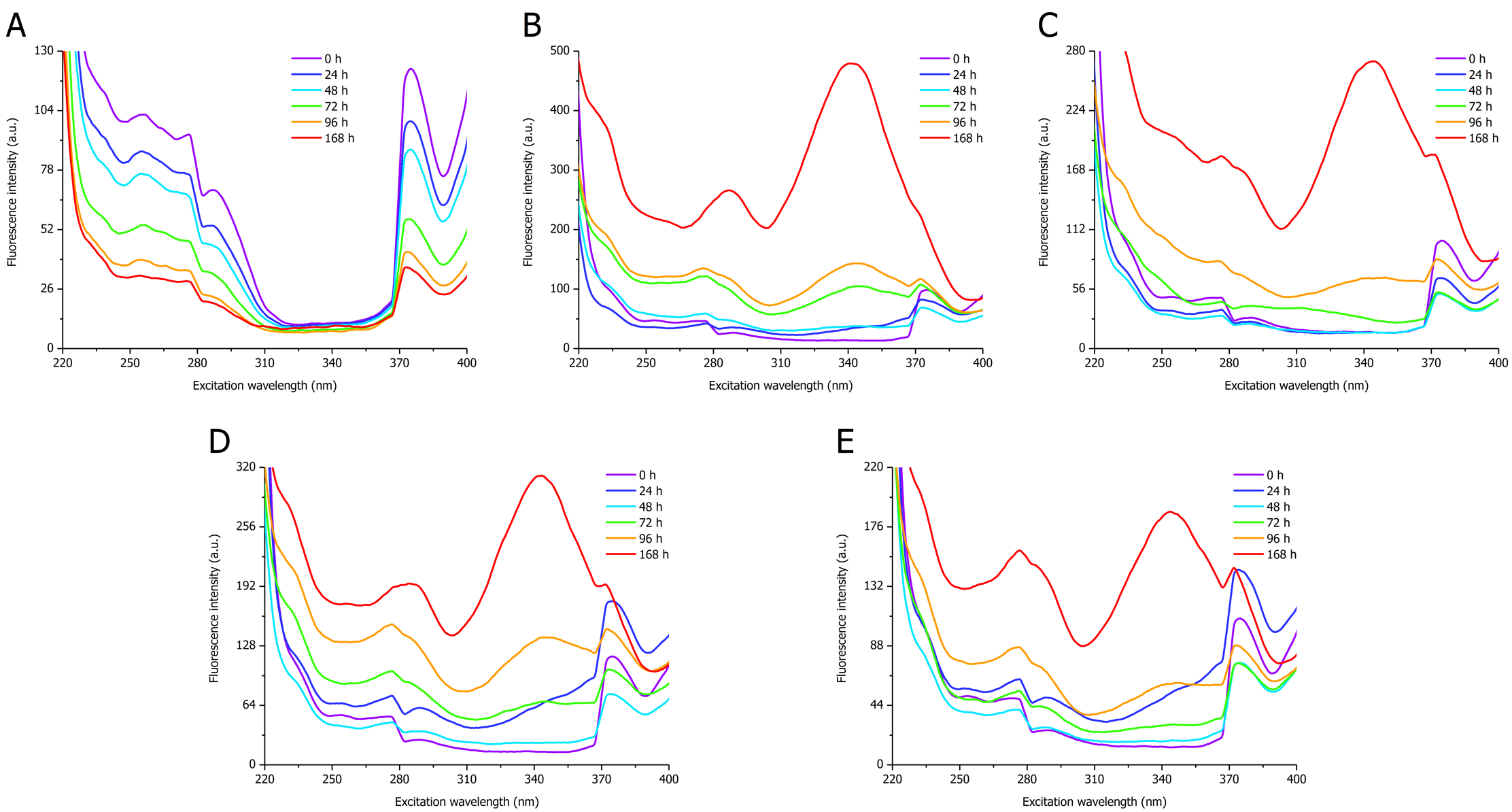

Figure S3 – AF excitation spectra (λ_em_ = 420 nm, sw = 10 nm) of full-length DPF3a incubated at ~25 °C and (A) pH 2, (B) 8, and (C) 12 in 150 mM NaCl, as well as (D) 300 and (E) 500 mM NaCl at pH 8 for 0 h (purple), 24 h (dark blue), 48 h (light blue), 72 h (green), 96 h (orange), and 168 h (red).

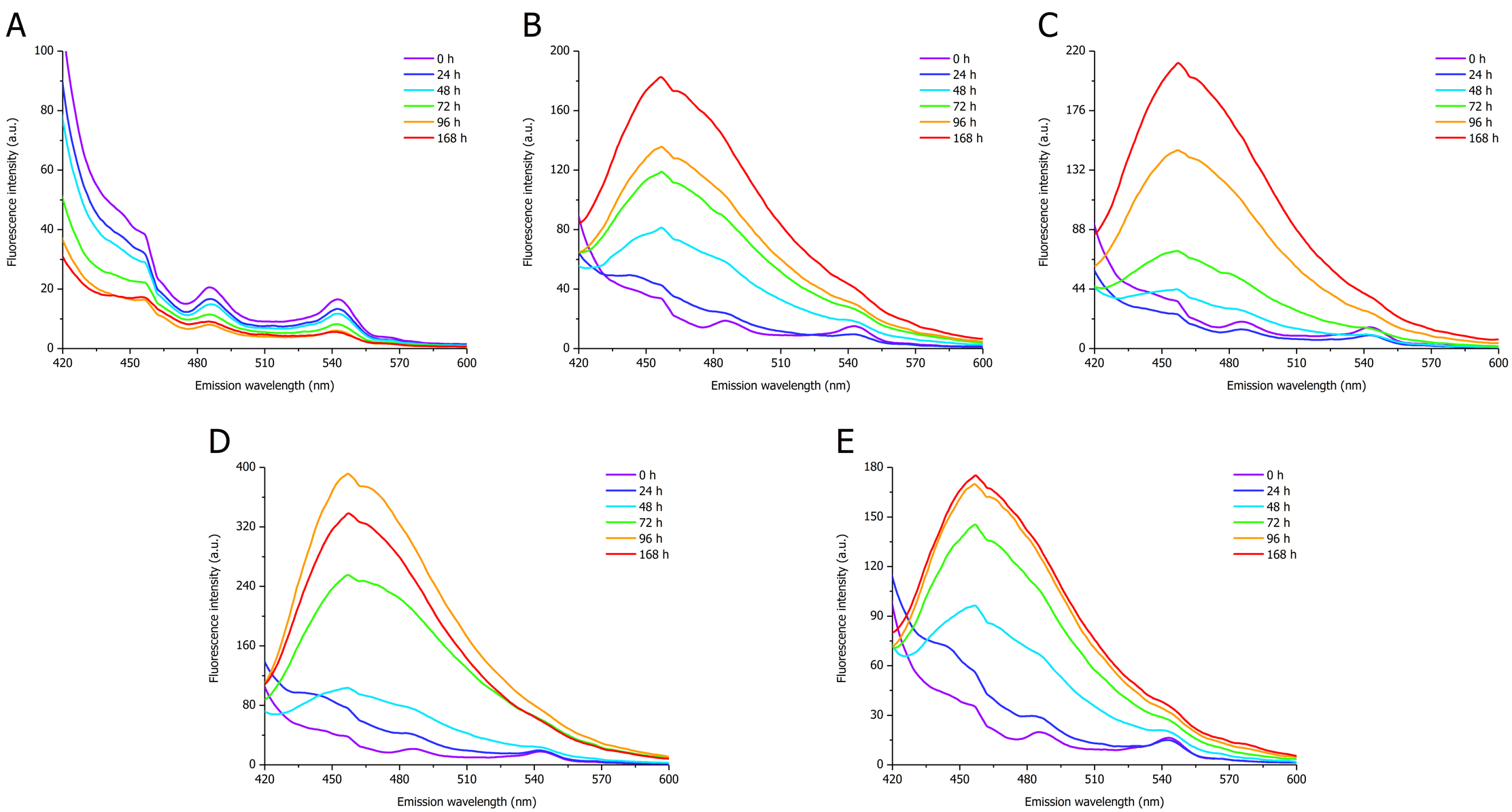

Figure S4 – AF spectra (λ_ex_ = 400 nm, sw = 10 nm) of full-length DPF3a incubated at ~25 °C and (A) pH 2, (B) 8, and (C) 12 in 150 mM NaCl, as well as (D) 300 and (E) 500 mM NaCl at pH 8 for 0 h (purple), 24 h (dark blue), 48 h (light blue), 72 h (green), 96 h (orange), and 168 h (red).

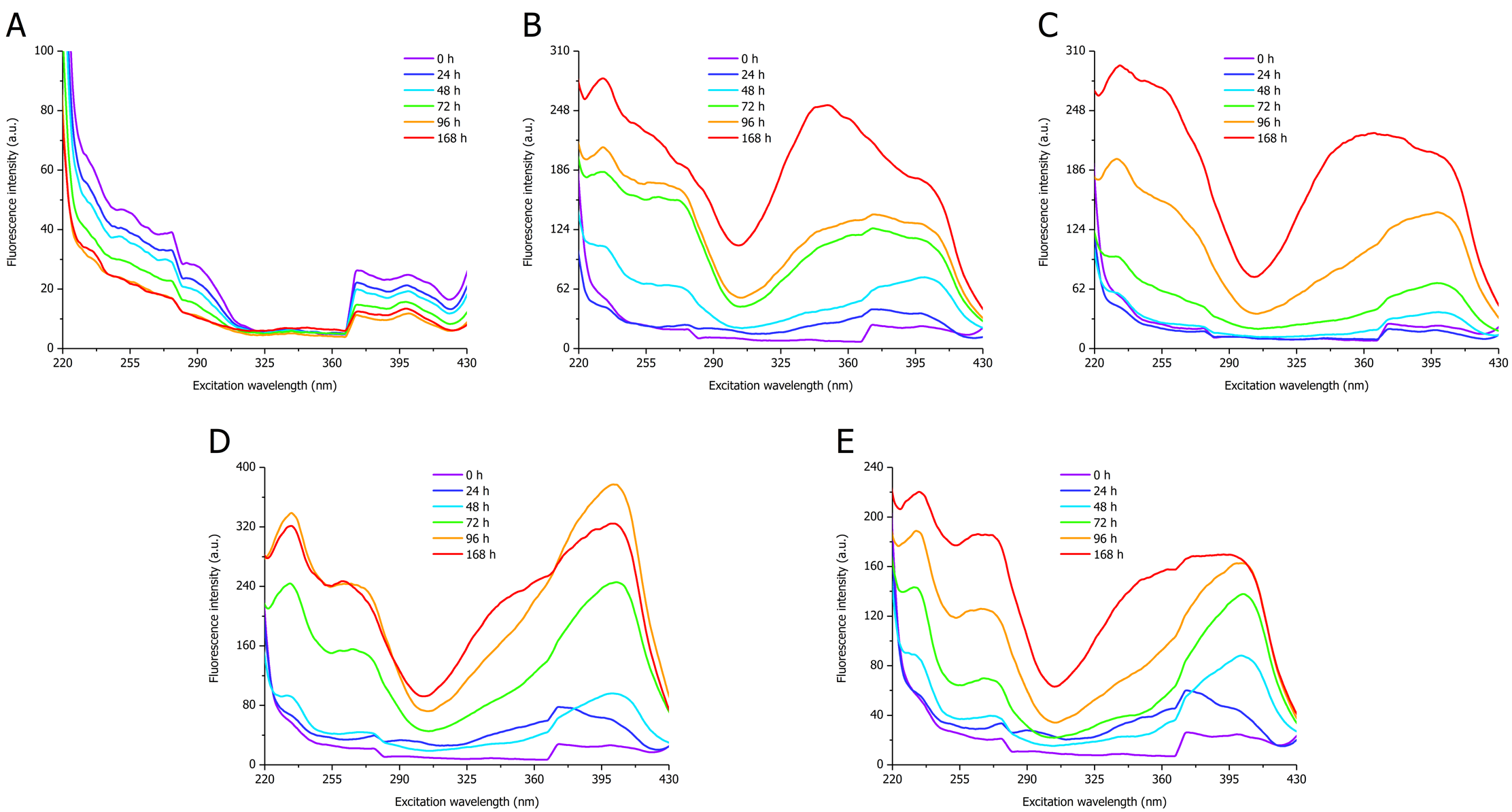

Figure S5 – AF excitation spectra (λ_em_ = 460 nm, sw = 10 nm) of full-length DPF3a incubated at ~25 °C and (A) pH 2, (B) 8, and (C) 12 in 150 mM NaCl, as well as (D) 300 and (E) 500 mM NaCl at pH 8 for 0 h (purple), 24 h (dark blue), 48 h (light blue), 72 h (green), 96 h (orange), and 168 h (red).

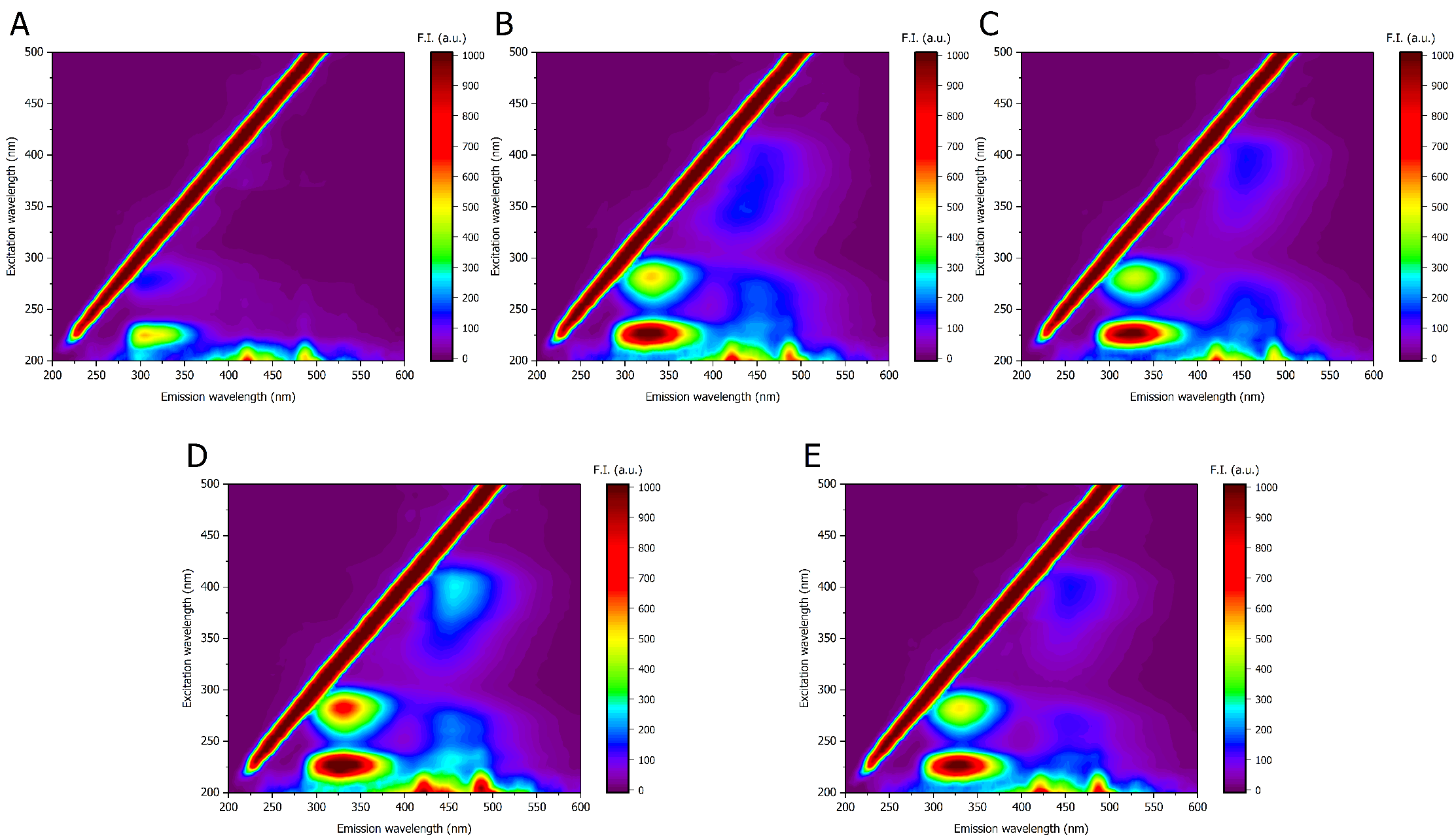

Figure S6 – EEM (sw = 10 nm) of full-length DPF3a incubated for 96 h at ~25 °C and (A) pH 2, (B) 8, and (C) 12 in 150 mM NaCl, as well as (D) 300 and (E) 500 mM NaCl at pH 8.

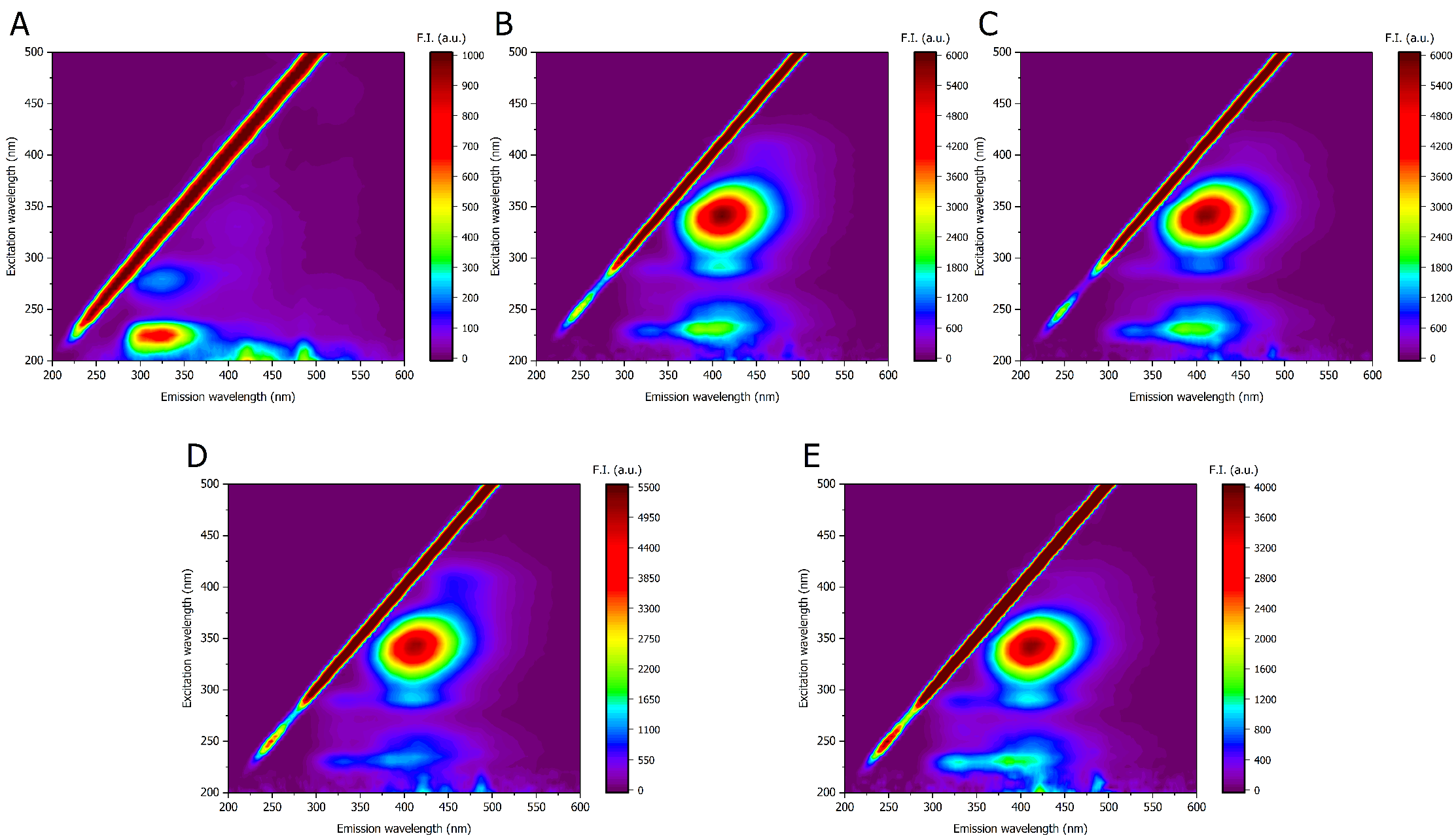

Figure S7 – EEM (sw = 10 nm) of full-length DPF3a incubated for 6 weeks at ~25 °C and (A) pH 2, (B) 8, and (C) 12 in 150 mM NaCl, as well as (D) 300 and (E) 500 mM NaCl at pH 8.

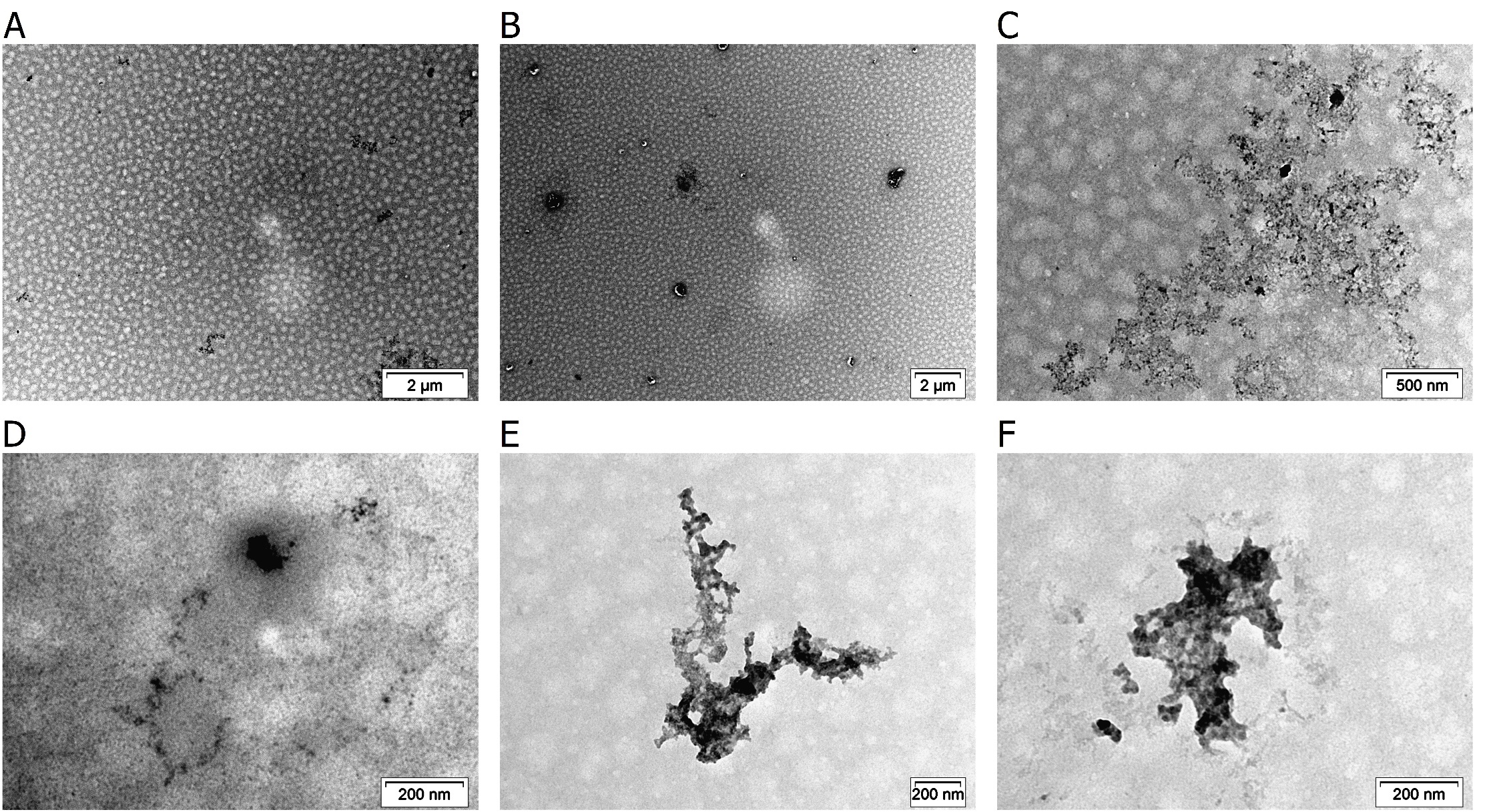

Figure S8 – Negatively stained TEM micrographs of full-length DPF3a incubated for 168 h at ~25 °C and pH 2 in 150 mM NaCl. (A-B) Representative overview of the grid particularly characterised by the absence of fibrillar aggregates and the sporadic presence of small amorphous clusters. (C-F) Amorphous phases and aggregates. The scale bar is provided at the bottom right of each micrograph.

s

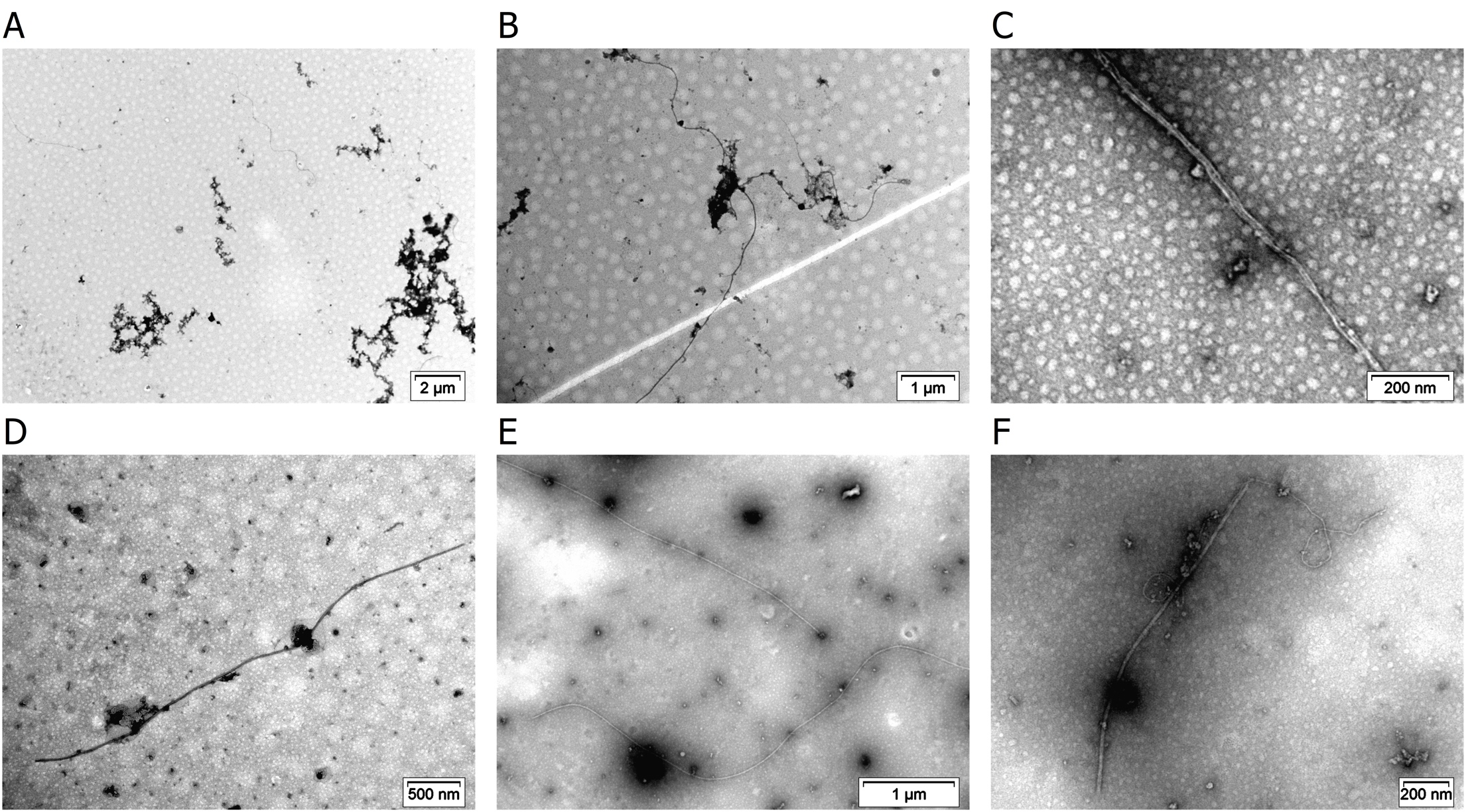

Figure S9 – Negatively stained TEM micrographs of full-length DPF3a incubated for 168 h at ~25 °C and pH 8 in 150 mM NaCl. (A) Representative overview of the grid characterised by the presence of fibrils (black arrow), spherical oligomers (red arrow), and curvilinear protofibrils or aggregates (blue arrow). (B) Most representative fibrillar morphology with respect to the condition (also refer to Figure 6B in the main text). Note the presence of cobweb-like fibrillar networks entangled with the fibril (black arrow). (C) Assembly of very thin fibrils. (D) Straight and isolated fibril. (E) Extensively long straight (black arrow) and curved (red arrow) fibrils. (F) Assembly of thin and curved fibril (black arrow) with straight fibril (red arrow). The scale bar is provided at the bottom right of each micrograph.

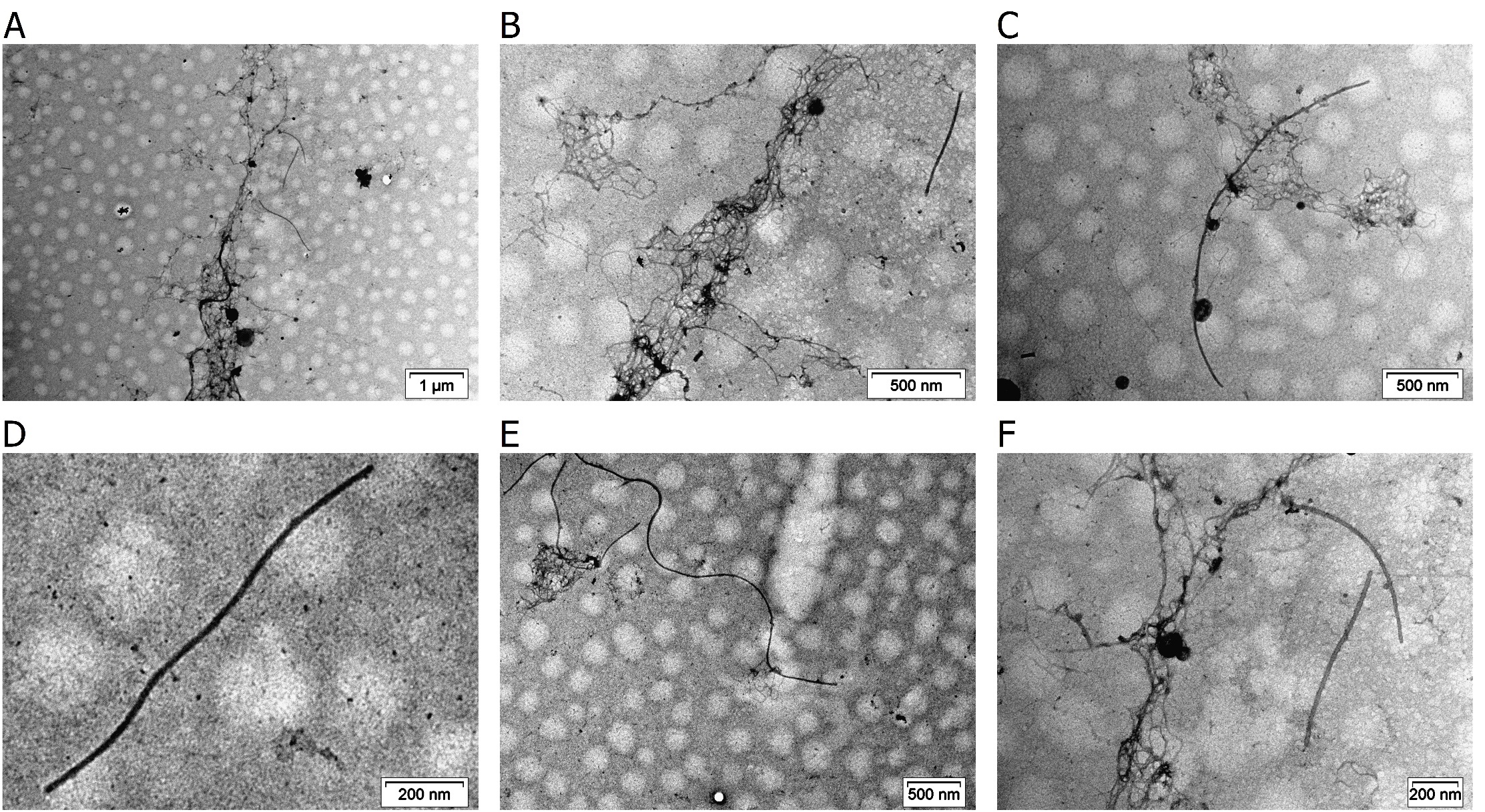

Figure S10 – Negatively stained TEM micrographs of full-length DPF3a incubated for 168 h at ~25 °C and pH 12 in 150 mM NaCl. (A) Representative overview of the grid characterised by the presence of fibrils (black arrow), spherical oligomers (red arrow), and dense network of very thin and cobweb-like fibrils (blue arrow). (B) Most representative fibrillar morphology with respect to the condition (also refer to Figure 6C in the main text). (C) Local association and coexistence of curve fibril (black arrow) with cobweb-like fibrillar networks (red arrow). (D) Straight and isolated fibril. (E) Long curved and undulating fibrils. (F) Coexistence of isolated straight fibril (black arrow) with cobweb-like fibrils (red arrow). The scale bar is provided at the bottom right of each micrograph.

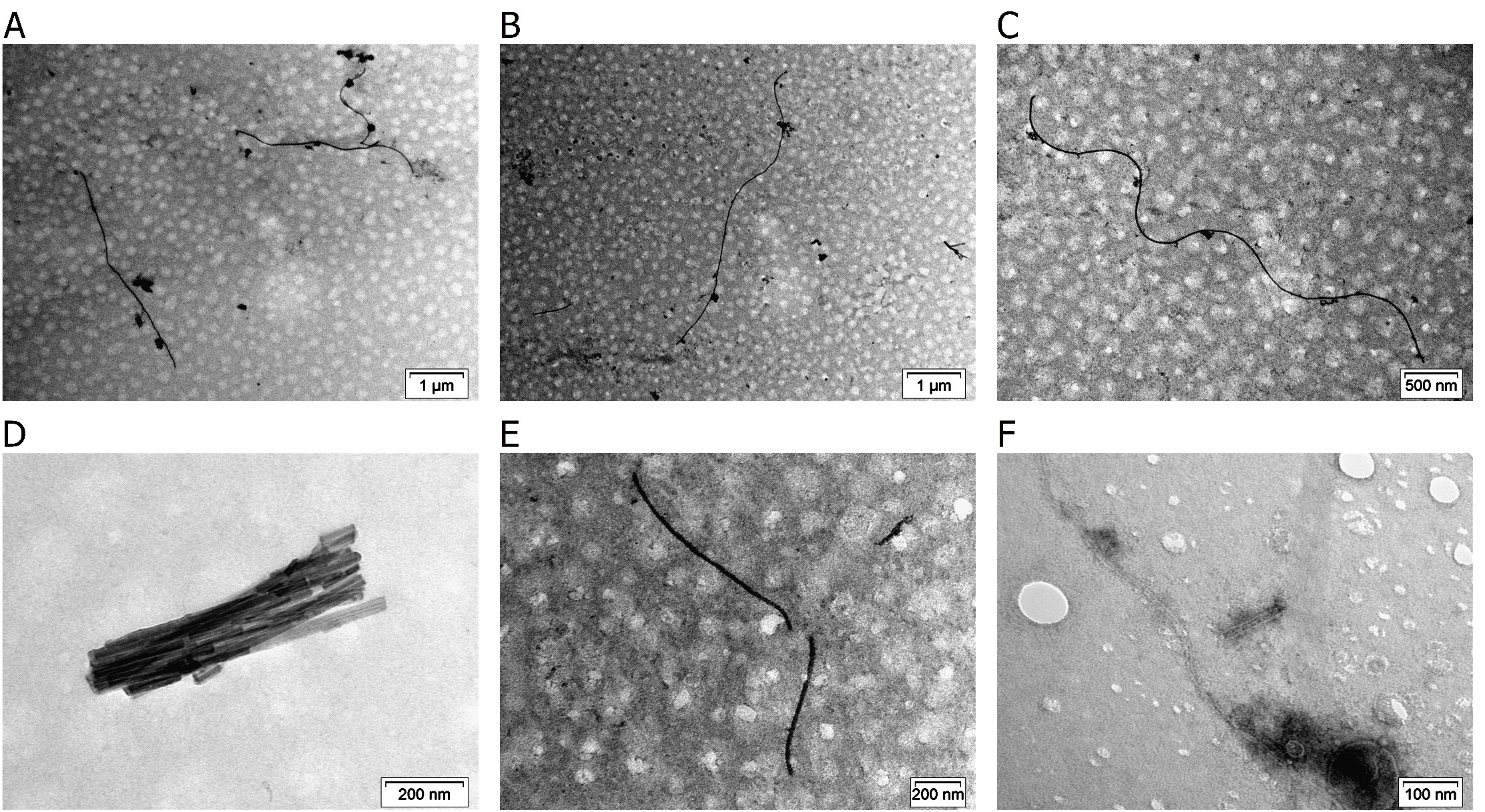

Figure S11 – Negatively stained TEM micrographs of full-length DPF3a incubated for 168 h at ~25 °C and pH 8 in 300 mM NaCl. (A) Representative overview of the grid characterised by the presence of straight (black arrow) and curved (red arrow) fibrils. (B) Most representative fibrillar morphology with respect to the condition (also refer to Figure 6D in the main text). (C) Long curved and undulating fibril. (D) Short and crystal-like fibrillar aggregates. (E) Short straight and isolated fibrils. (F) Assembly of very thin fibrils. The scale bar is provided at the bottom right of each micrograph.

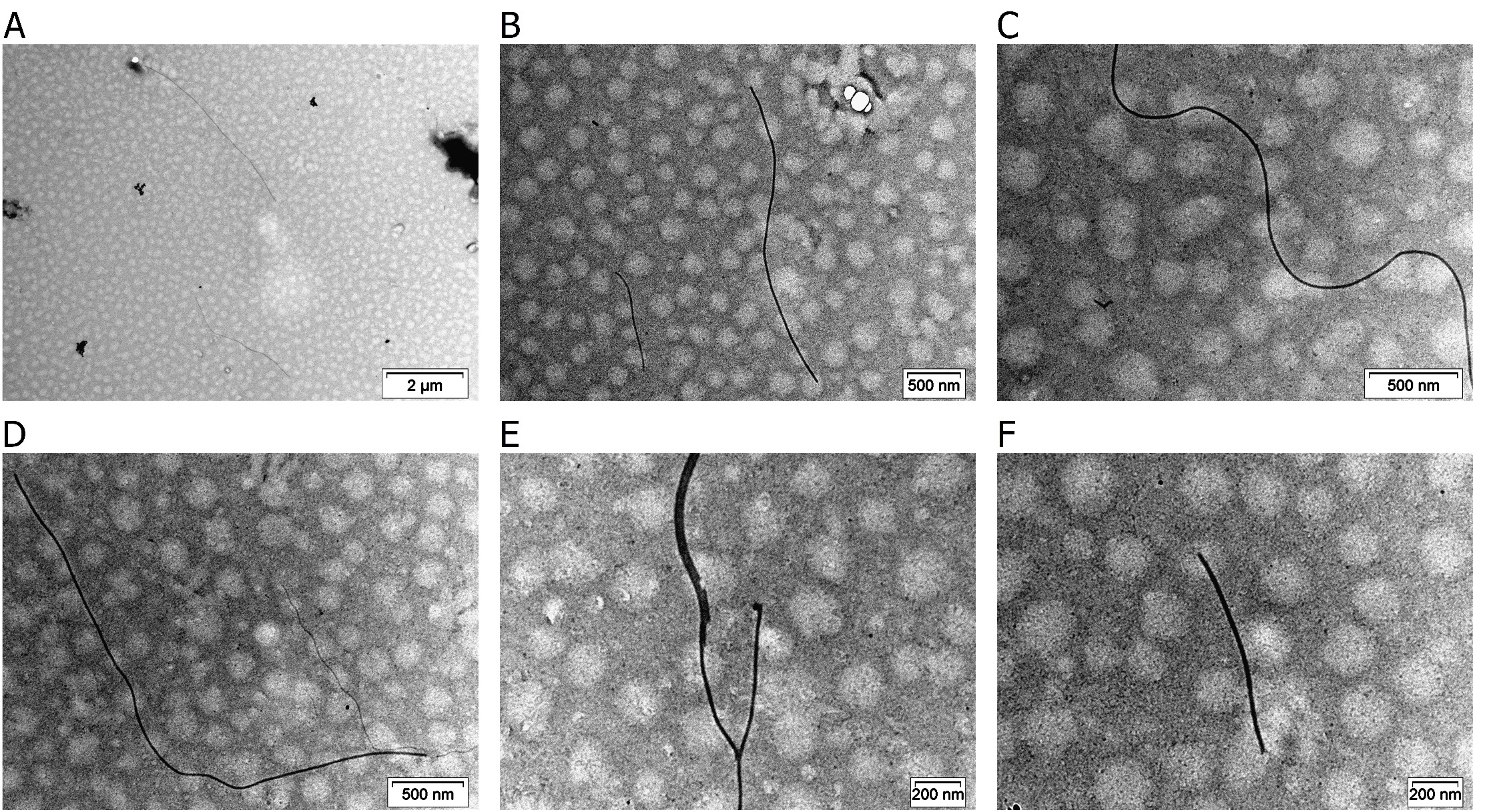

Figure S12 – Negatively stained TEM micrographs of full-length DPF3a incubated for 168 h at ~25 °C and pH 8 in 500 mM NaCl. (A) Representative overview of the grid characterised by the presence of straight (black arrow) fibrils. (B) Most representative fibrillar morphology with respect to the condition (also refer to Figure 6E in the main text). (C) Long curved and undulating fibril. (D) Coexistence of straight fibril (black arrow) with thinner ones (red arrow). (E) Assembly of thin fibrils. (F) Short straight and isolated fibril. The scale bar is provided at the bottom right of each micrograph.

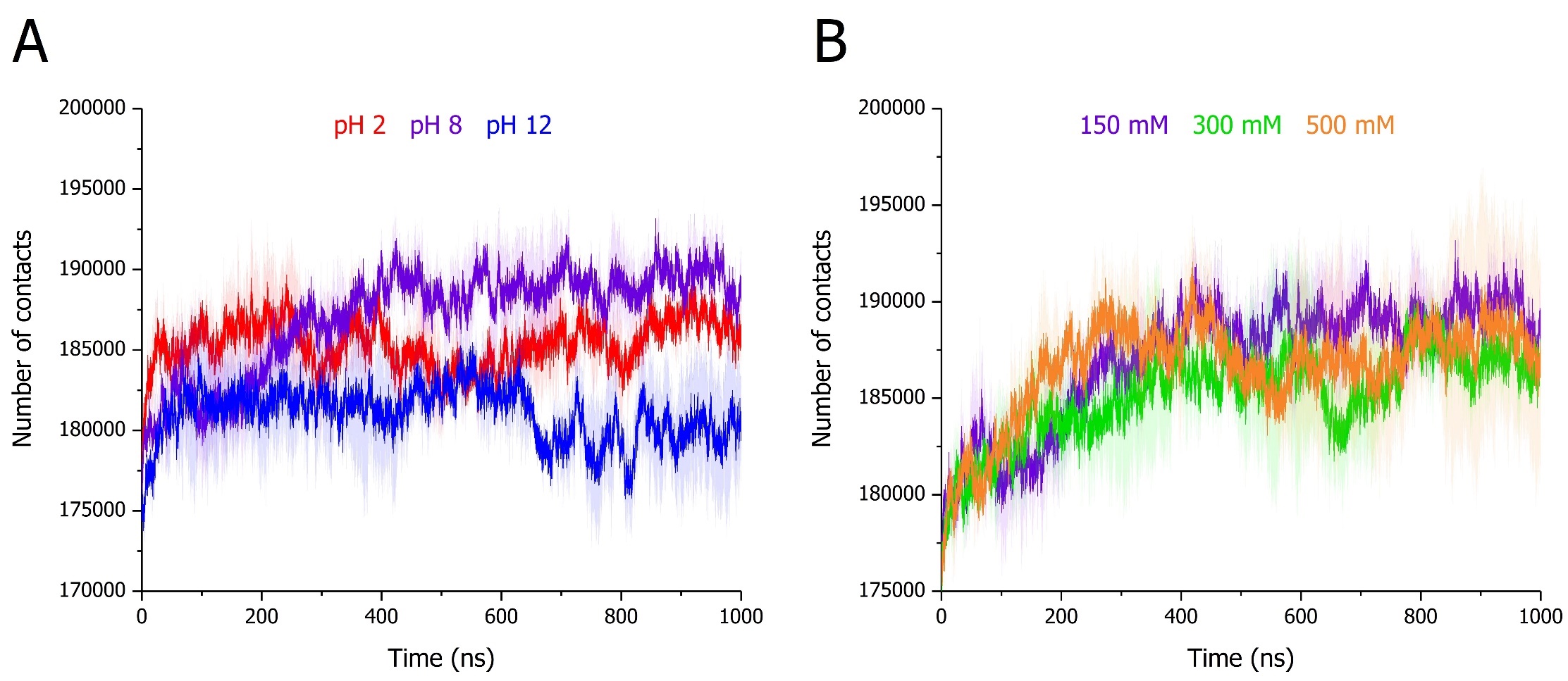

Figure S13 – Time-evolution of intramolecular number of contacts of full-length DPF3a simulated for 1 µs at (A) pH 2 (red), 8 (purple), and 12 (blue) in 150 mM NaCl, as well as (B) in 150 (purple), 300 (green), and 500 mM NaCl (orange) at pH 8. For each condition, curves correspond to the average of triplicates with the standard deviation represented as a trace (shaded area) in the condition-associated colour.

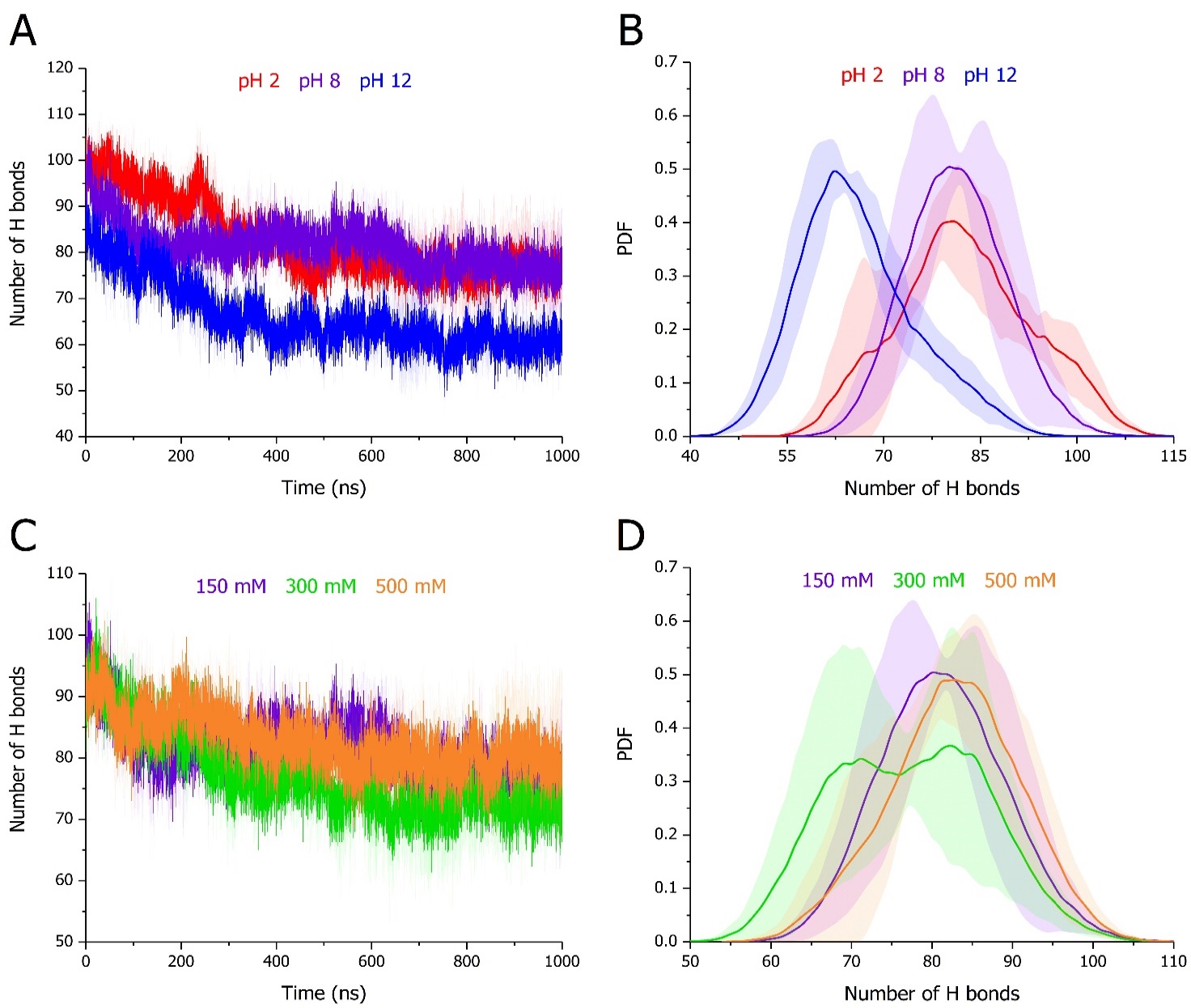

Figure S14 – Intramolecular backbone H bond (A,C) evolution over time and (B,D) distribution over the trajectories of full-length DPF3a simulated for 1 µs at (A-B) pH 2 (red), 8 (purple), and 12 (blue) in 150 mM NaCl, as well as (C-D) in 150 (purple), 300 (green), and 500 mM NaCl (orange) at pH 8. For each condition, curves correspond to the (A,C) time-evolution and (B,D) probability density function (PDF) average of triplicates with the standard deviation represented as a trace (shaded area) in the condition-associated colour.

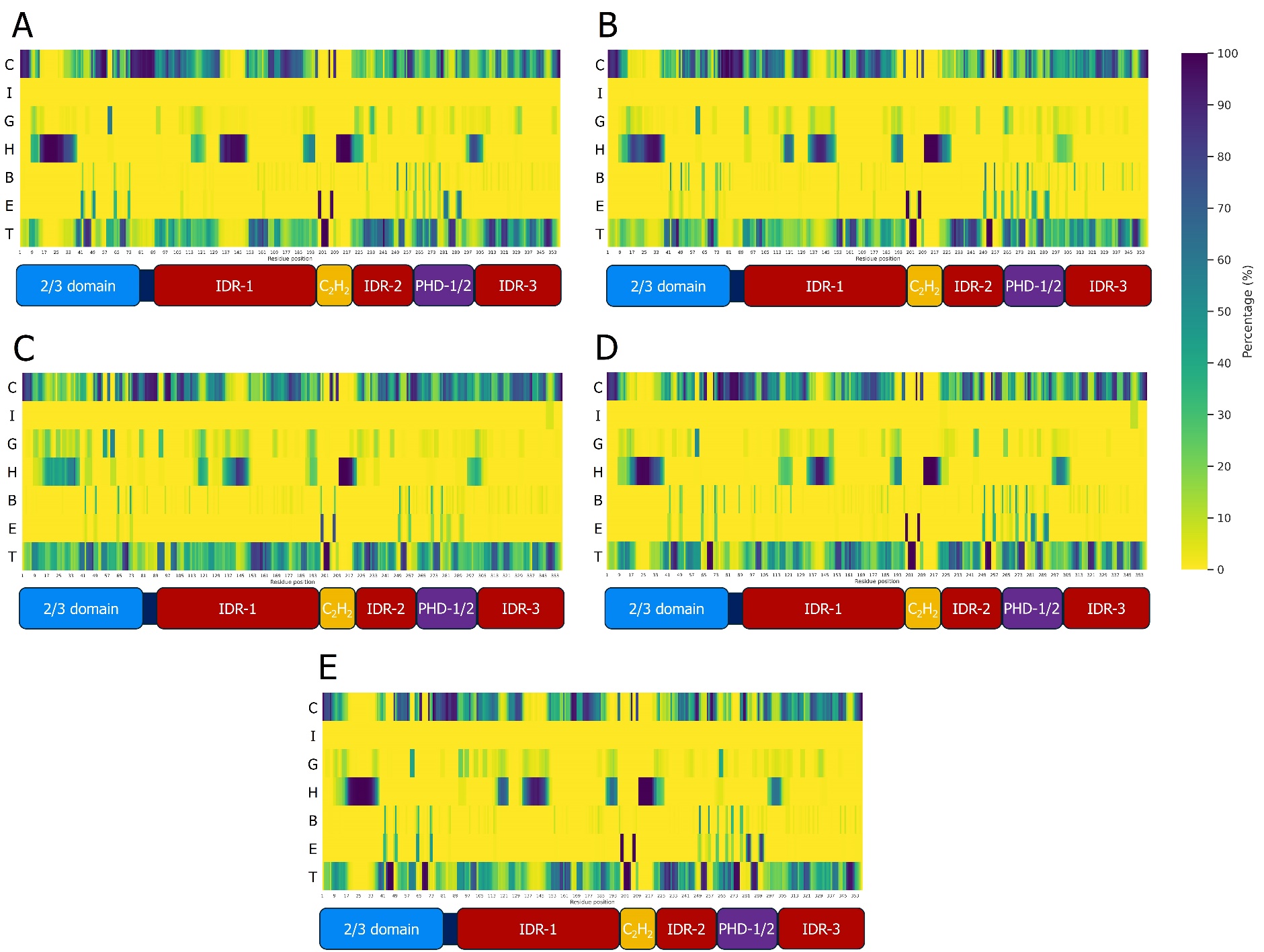

Figure S15 – Mean per-residue occurrence of secondary structure types over the trajectories of full-length DPF3a simulated for 1 µs at (A) pH 2, (B) 8, and (C) 12 in 150 mM NaCl, as well as (D) in 300 and (E) 500 mM NaCl at pH 8. The secondary structures are defined into seven classes after STRIDE assignment: turn (T), extended β-sheet (E), isolated β-bridge (B), α-helix (H), 3_10_-helix (G), π-helix (I), and random coil (C). For each condition, per-residue percentages correspond to the average of triplicates. The percentage scale goes from 0 (yellow) to 100% (dark blue) of occurrence. On each graph, the sequence organisation of DPF3a is displayed according to its constitutive domains: the N-terminal 2/3 domain (blue), intrinsically disordered regions (dark red), the Krüppel-like C_2_H_2_ zinc finger (yellow), and truncated PHD-1/2 (purple).

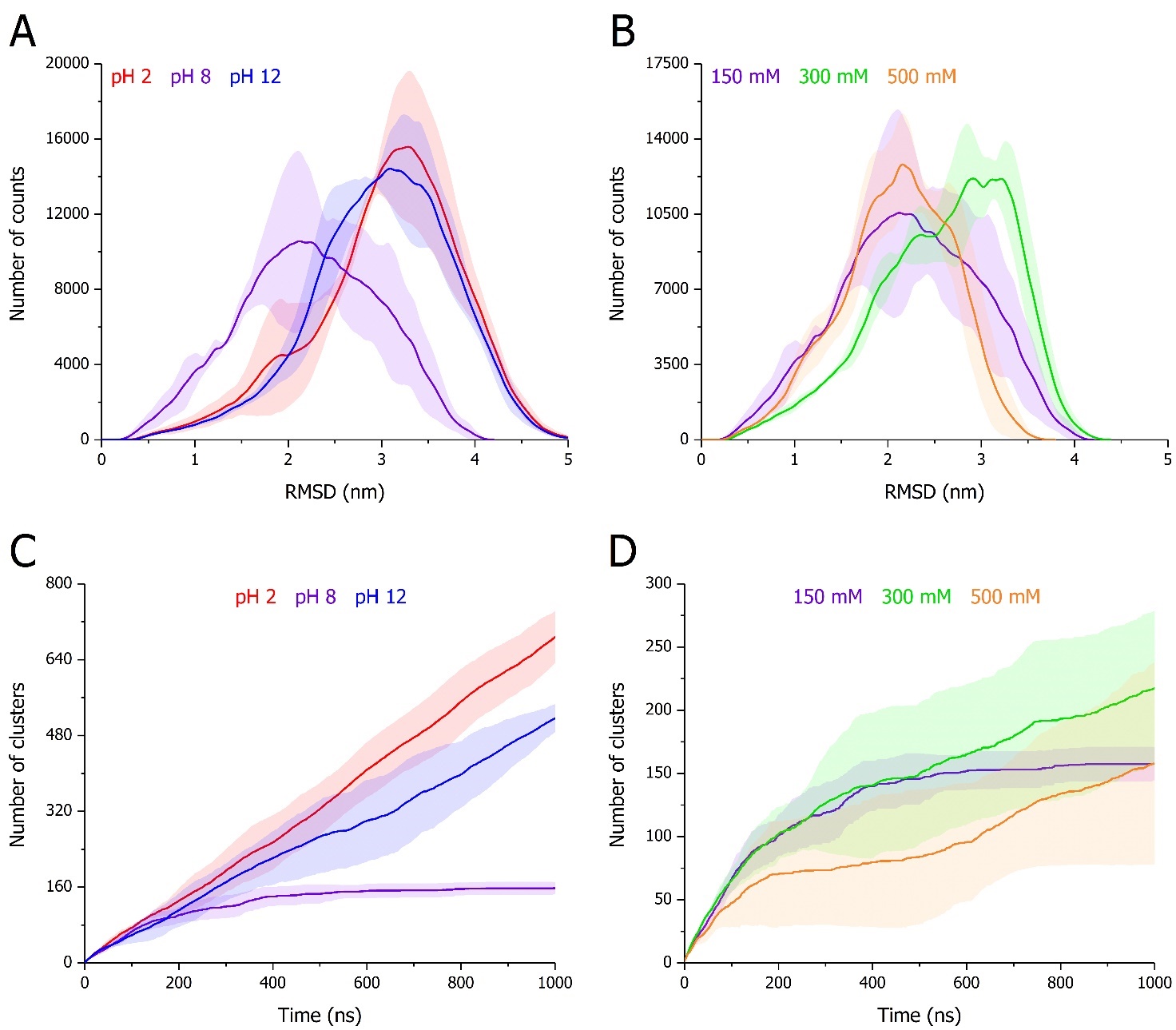

Figure S16 – Distribution of (A-B) RMSD and (C-D) time-evolution of number of clusters of full-length DPF3a simulated for 1 µs at (A,C) pH 2 (red), 8 (purple), and 12 (blue) in 150 mM NaCl, as well as (B,D) in 150 (purple), 300 (green), and 500 mM NaCl (orange) at pH 8. For each condition, curves correspond to the average of triplicates with the standard deviation represented as a trace (shaded area) in the condition-associated colour.

Figure S17 – Conformational clustering analysis and residue pairs contact mapping of full-length DPF3a simulated for 1 µs at (A) pH 2, 8 (upper panel), and (B) 12 in 150 mM NaCl, as well as in (C) 300 and (D) 500 mM NaCl at pH 8. For each condition, the protein structure of the final frame of the most representative trajectory (refer to Figure 13 in the main text) amongst the triplicates (upper row) is shown in cartoon representation with the N- and C-termini respectively pinpointed as blue and red spheres, and the different secondary structure elements coloured according to STRIDE assignment: turn (turquoise), extended β-sheet (yellow), isolated β-bridge (tan), α-helix (pink), 3_10_-helix (blue), π-helix (red), and random coil (light grey). The coordinated Zn^2+^ cation is represented as a grey bead. Percentages indicate the population size in which the final frame is found over the selected trajectory. Minimum distance contact maps between residue pairs (bottom row) are compared between the different pH and ionic strength conditions (bottom half on each map) with respect to the pH 8 and 150 mM NaCl system taken as a reference (upper half on each map). On the sides of each contact map, the sequence organisation of DPF3a is displayed according to its constitutive domains: the N-terminal 2/3 domain (blue), intrinsically disordered regions (dark red), the Krüppel-like C_2_H_2_ zinc finger (yellow), and truncated PHD-1/2 (purple).

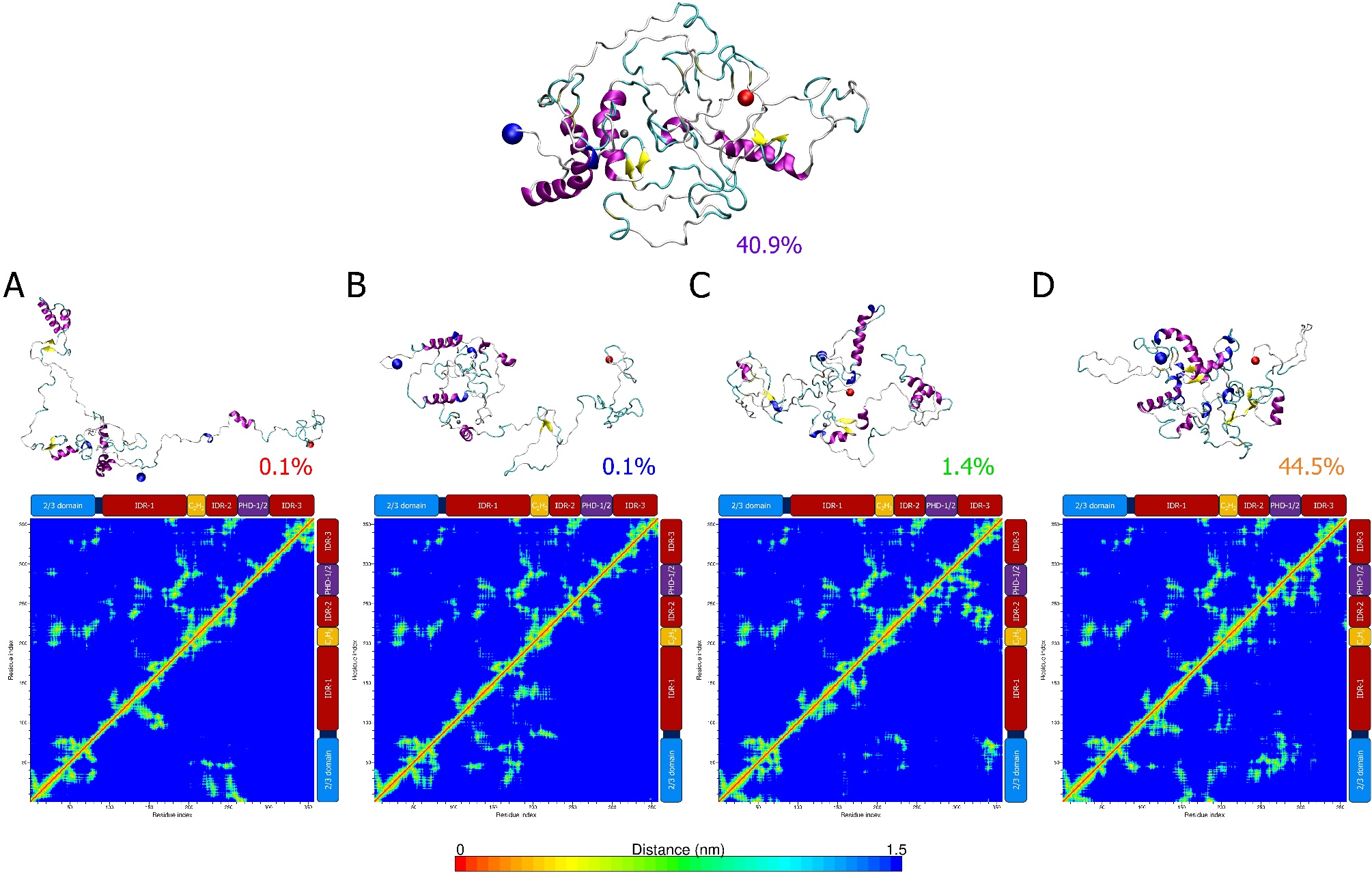

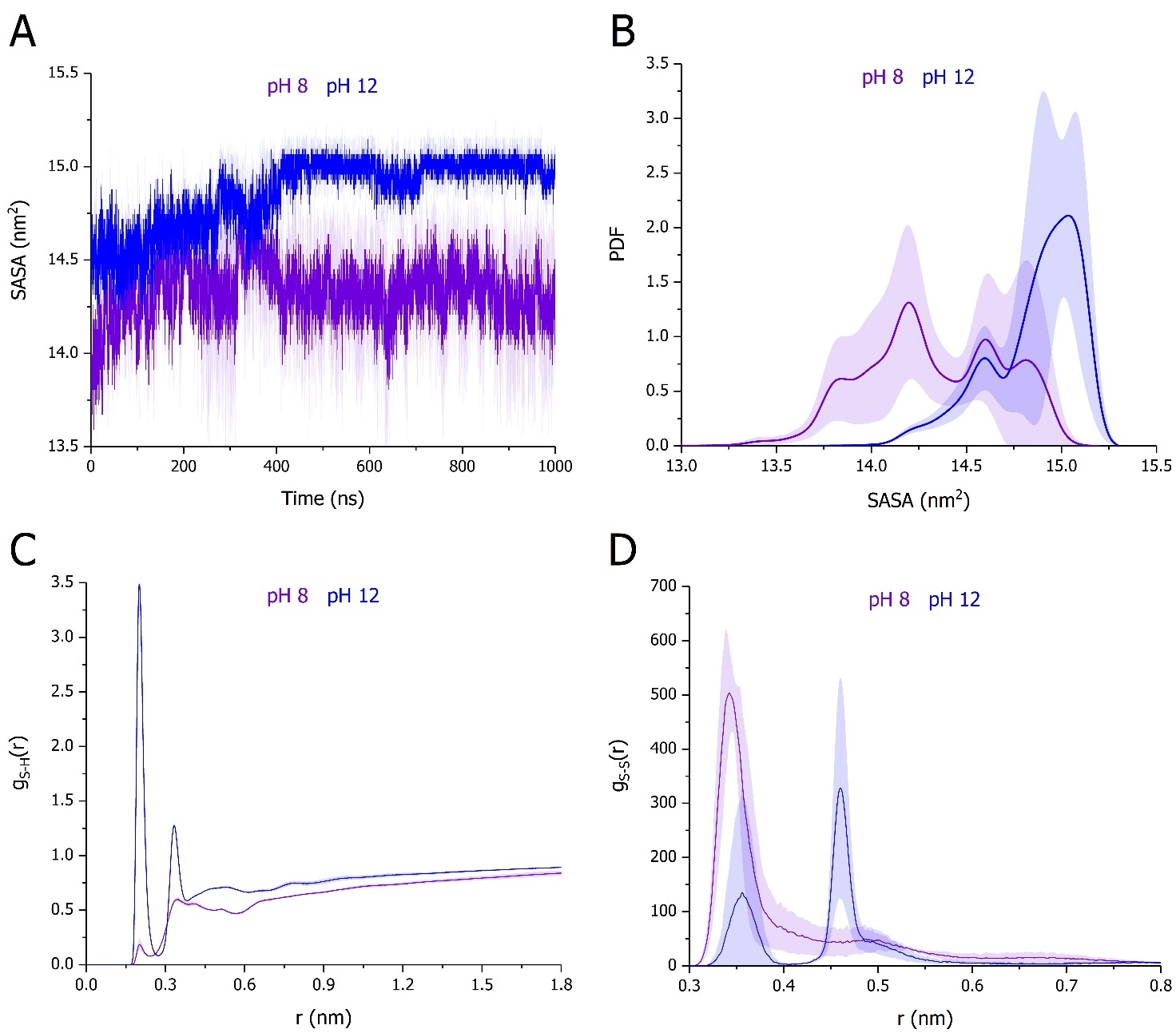

Figure S18 – SASA (A) evolution over time and (B) distribution over the trajectories of the cysteine Sγ atoms of full-length DPF3a simulated for 1 µs at pH 8 (purple) and 12 (blue) in 150 mM NaCl. For each condition, curves correspond to the (A) time-evolution and (B) probability density function (PDF) average of triplicates with the standard deviation represented as a trace (shaded area) in the condition-associated colour. Radial distribution function g(r) of (C) cysteine Sγ-water H atoms and (D) cysteine Sγ-Sγ atoms. For each condition, curves correspond to the average of triplicates with the standard deviation represented as a trace (shaded area) in the condition-associated colour.
